# Design, immunogenicity, and efficacy of a pan-sarbecovirus dendritic-cell targeting vaccine

**DOI:** 10.1101/2021.12.28.474244

**Authors:** Séverin Coléon, Aurélie Wiedemann, Mathieu Surénaud, Christine Lacabaratz, Sophie Hue, Mélanie Prague, Minerva Cervantes-Gonzalez, Zhiqing Wang, Jerome Ellis, Amandine Sansoni, Camille Pierini, Quentin Bardin, Manon Fabregue, Sarah Sharkaoui, Philippe Hoest, Léa Dupaty, Florence Picard, Mireille Centlivre, Jade Ghosn, French COVID Cohort Study Group, Rodolphe Thiébaut, Sylvain Cardinaud, Bernard Malissen, Gérard Zurawski, Ana Zarubica, Sandra M Zurawski, Véronique Godot, Yves Lévy

## Abstract

The emergence of SARS-CoV-2 variants of concern (VOCs) that escape pre-existing antibody neutralizing responses increases the need for vaccines that target conserved epitopes and induce cross-reactive B- and T-cell responses. We used a computational approach and sequence alignment analysis to design a new-generation subunit vaccine targeting conserved sarbecovirus B- and T-cell epitopes from Spike (S) and Nucleocapsid (N) to antigen-presenting cells expressing CD40 (CD40.CoV2). We demonstrate the potency of CD40.CoV2 to elicit high levels of cross-neutralizing antibodies against SARS-CoV-2, VOCs, and SARS-CoV-1 in K18-hACE2 transgenic mice, associated with improved viral control and survival after challenge. In addition, we demonstrate the potency of CD40.CoV2 *in vitro* to recall human multi-epitope, functional, and cytotoxic SARS-CoV-2 S- and N-specific T-cell responses that are unaffected by VOC mutations and cross-reactive with SARS-CoV-1 and, to a lesser extent, MERS epitopes. Overall, these findings provide a framework for a pan-sarbecovirus vaccine.

## Introduction

Severe acute respiratory syndrome coronavirus 2 (SARS-CoV-2), which emerged in late 2019 in the Hubei province of China, has caused devastating human and economic losses worldwide. Unprecedented mobilization of the scientific community has led to the implementation of viral diagnostics, immunological monitoring tools, and the rapid development of protective vaccines.

The current landscape of COVID-19 vaccines is based on the delivery of SARS-CoV-2 Spike (S) through various vaccine platforms that elicit neutralizing-antibody responses against the S protein, including the receptor binding domain (RBD). Most of these vaccines induce Th1 responses restricted to S epitopes, depending on the type of platform and variations in the S protein (Anderson et al., 2020; Angyal et al., 2021; Bange et al., 2021; Guccione; Kalimuddin et al.; Lederer et al., 2020; Mazzoni et al., 2021; Painter et al., 2021; Prendecki et al., 2021; Sahin et al., 2020; Stamatatos et al., 2021; Tarke et al., 2021a), but vary in their capacity to elicit CD8^+^ T-cell responses, known to be an important element for control of the infection (McMahan et al., 2021). Although neutralizing antibodies are a key component of a broadly protective vaccine, several recent studies have suggested that the induction of broad virus-specific CD4^+^ and CD8^+^ T cells could greatly augment antibody-based protection and the long-term durability of vaccine responses (Sette and Crotty, 2021).

Despite current progress, control of the ongoing SARS-CoV-2 pandemic is endangered by the emergence of viral variants, called variants of concern (VOCs). Among them, the B.1.1.7 Alpha (Funk et al., 2021), B.1.351 Beta (Tegally et al., 2021), P.1 Gamma (Voloch et al., 2021), B.1.617.2 Delta (Cherian et al., 2021), and recently emerged B.1.1.529 Omicron variants (Karim and Karim, 2021) exhibit several specific or shared mutations within the S sequences, raising substantial new concerns due to their increased transmissibility (Davies et al., 2021; Kumar et al., 2021) and ability to escape convalescent and vaccine-induced antibody responses (Garcia-Beltran et al., 2021; Hoffmann et al., 2021; Madhi et al., 2021; Wang et al., 2021; Wibmer et al., 2021). Recent studies showing a decrease in the effectiveness of mRNA vaccines against the new VOCs (Puranik et al.) report of breakthrough infections (Kustin et al., 2021), and concerns of reduced efficacy of vaccination in older patients (Israel et al., 2021) or immune-compromised individuals (Hadjadj et al., 2021) highlight the need to develop a new and complementary generation of vaccines as prophylaxis or boosters that include new T- and B-cell selected antigens that are potentially less affected by the mutations of VOCs.

Within the last 20 years, SARS-CoV-2 is the third major human infectious disease outbreak caused by zoonotic coronaviruses after SARS-CoV-1 in 2002-2003 and Middle East respiratory syndrome coronavirus (MERS-CoV) in 2012. The first available sequence of SARS-CoV-2 (Wu et al., 2020) identified this novel human pathogen as a member of the *Sarbecovirus* subgenus of *Coronaviridae* (Lu et al., 2020), the same subgenus as SARS-CoV-1. The high prevalence and diversity of viruses in bats and the fact that all zoonotic sarbecoviruses identified to date use hACE2 as their entry receptor raise major concerns about a future epidemic (Boni et al., 2020; Cohen, 2021; Frutos et al., 2021; Menachery et al., 2015, 2016). These additional concerns underscore the urgent need for new vaccine candidates that are able to enhance protection against VOCs and emerging coronaviruses.

Dendritic cells (DCs) are immune system controllers that can deliver differential signals to other immune cells through intercellular interactions and soluble factors, resulting in a variety of host immune responses of varying quality. Targeting vaccine antigens to DCs via surface receptors represents an appealing strategy to improve subunit-vaccine efficacy while reducing the amount of required antigen. This strategy, which allows the delivery of designed and selected antigens, in addition to an activation signal, may also evoke a danger signal that stimulates an immune response, with or without the need of additional immune stimulants, such as adjuvants. Among the various DC receptors tested, including lectins and scavenger receptors, we previously reported the superiority of vaccines targeting diverse viral antigens to CD40-expressing antigen-presenting cells to evoke strong antigen-specific T- and B-cell responses (Bouteau et al., 2019; Chatterjee et al., 2012; Cheng et al., 2018; Flamar et al., 2013, 2018; Godot et al., 2020; Yin et al., 2016; Zurawski et al., 2017).

We have recently reported results on the efficacy of a new generation of subunit vaccines targeting the RBD of the SARS-CoV-2 spike protein to the CD40 receptor (aCD40.RBD) (Marlin et al., 2021). We demonstrated that a single dose of the aCD40.RBD vaccine, injected without adjuvant, is sufficient to boost a rapid increase in neutralizing antibodies in convalescent non-human primates (NHPs) infected six months previously with SARS-CoV-2. Interestingly, the aCD40.RBD vaccine-elicited antibodies cross-neutralized D614G SARS-CoV-2 and the VOCs Alpha (B1.1.7) and, to a lesser extent, Beta (B1.351). This vaccination significantly improved protection against a new high-dose virulent challenge versus that in non-vaccinated convalescent animals (Marlin et al., 2021).

Drawing from this knowledge, we used *in silico* approaches to design a next-generation CD40-targeting vaccine, CD40.CoV2, including new T- and B-cell epitopes spanning sequences from S and nucleocapsid (N) proteins from SARS-CoV-2 and highly homologous to 38 sarbecoviruses, including SARS-CoV-2 VOCs. We report here the immunogenicity and antiviral efficacy of this vaccine in a preclinical model.

## Results

### *In silico* down-selection of T- and B-cell polyepitope regions from SARS-CoV-2 for an improved dendritic cell-targeting vaccine platform

We first screened three structural proteins (S, N, and M) of SARS-CoV-2 for the identification of T-cell epitopes using NetMHC 4.0 (Andreatta and Nielsen, 2016) and NetMHCII 2.3 (Jensen et al., 2018) software, which predict peptides that bind to a large panel of class-I and -II HLAs, respectively. Linear B-cell epitopes were predicted using BepiPred 2.0 (Jespersen et al., 2017). We mapped a set of 9-mer epitopes binding to 80 HLA-class I molecules and 15-mer epitopes binding to 54 HLA-class II molecules, as well as all linear B-cell epitopes. We evaluated amino-acid (aa) regions encompassing both the highest number of predicted epitopes and the largest HLA coverage. Selected epitope-enriched regions were further screened for their sequence homology with other β coronaviruses, including SARS (designed thereafter as SARS-CoV-1), focusing on described T-cell epitopes and B-cell epitopes that generate neutralizing antibodies. Down-selected regions containing epitopes were also compared to early predicted and described SARS-CoV-2 T- and B-cell epitopes (Ahmed et al., 2020; Baruah and Bose, 2020; Dahlke et al., 2020; Fast et al., 2020; Grifoni et al., 2020; Hyun-Jung Lee and Koohy, 2020; Le Bert et al., 2020; Mateus et al., 2020; Nelde et al., 2021; Peng et al., 2020; Prachar et al., 2020). Thus, four T-and B-cell epitope-enriched regions were selected as vaccine (v) regions, one from N: vN2 (aa 276-411) and three from S: vS1 (aa 125-250), vRBD (aa 318-541), and vS2 (aa 1056-1209) (Figure 1A-B). These epitope-enriched regions contain a total of 640 aa with 2,313 predicted CD8^+^ T-cell epitopes covering 100% of HLA-Class I haplotypes, 2,985 predicted CD4^+^ T-cell epitopes covering 100% of HLA-Class II haplotypes, and 17 predicted SARS-CoV-2 linear B-cell epitopes (Figure 1 A-B, Table S1). Then, we examined whether selected vaccine sequences significantly matched all described currently known SARS-CoV-2 T-cell epitopes (Ferretti et al., 2020; Gao et al., 2021; Hu et al., 2021; Kared et al., 2021; Le Bert et al., 2020; Mateus et al., 2020, 2020; Motozono et al., 2021; Nelde et al., 2021; Peng et al., 2020; Poran et al., 2020; Rha et al., 2021; Schulien et al., 2021; Shomuradova et al., 2020; Tarke et al., 2021b). We found the vaccine sequences to contain 71/171 (42%) and 21/44 (48%) described CD8^+^ T-cell epitopes for S and N, respectively (Table S2). These values were 57/123 (46%) and 21/53 (40%) for described CD4^+^ T-cell epitopes (Table S3).

**Figure 1.**
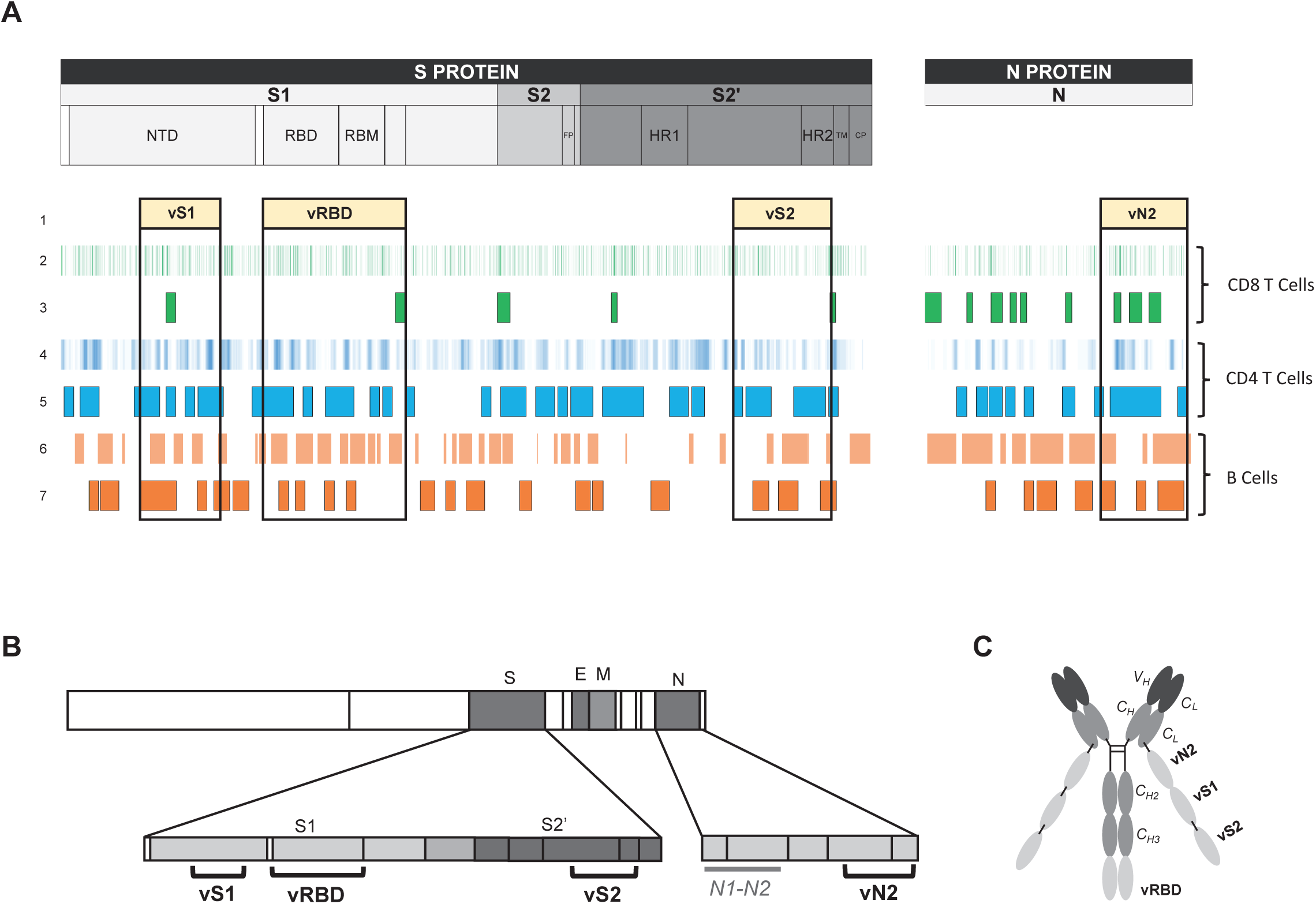
Selection of SARS-CoV-2 T- and B-cell polyepitope regions for an improved dendritic cell-targeting vaccine platform. (A) Mapping of selected SARS-CoV-2 epitope-enriched regions. (1) Four selected vaccine regions. (2) Predicted SARS-CoV-2 CD8^+^ T-cell epitopes (NetMHC 4.0). (3) Described SARS-CoV-2 CD8^+^ T-cell epitopes at the time of vaccine region selection. (4) Predicted SARS-CoV-2 CD4^+^ T-cell epitopes (NetMHCII 2.3). (5) Described SARS-CoV-2 CD4^+^ T-cell epitopes at the time of vaccine region selection. (6) Predicted linear B-cell epitopes (BepiPred 2.0). (7) Described SARS-CoV-2 IgM, IgA, and IgG epitopes at the time of vaccine region selection. (B) Vaccine (v) regions (vS1, vRBD, vS2 and vN2) and control region not included in the vaccine (N1-N2) and (C) CD40.CoV2 vaccine construct.

We then used Cobalt (https://www.ncbi.nlm.nih.gov/tools/cobalt/re_cobalt.cgi) to perform the alignment of sequences from SARS-CoV-2, four SARS-CoV-2 VOCs (α, β, γ, δ), SARS-CoV-1, and 32 recently described SARS-CoV-related coronaviruses (Lam et al., 2020; Wacharapluesadee et al., 2021; Zhou et al., 2020), which include 30 viruses of bat origin and two of pangolin origin (all from the *Sarbecovirus* subgenus). Globally, the mean [min-max] percentage of homology between these 38 sarbecoviruses for vS1, vRBD, vS2, and vN2 vaccine sequences was 53.5 [36.5 to 99.2], 73.6 [63.8 to 99.6], 94.3 [86.4 to 100], and 93.5 [89.7 to 100]%, respectively (Table S4). As expected, homology between vaccine sequences and members of *Embecovirus*, *Merbecovirus*, *Setracovirus*, and *Duvinacovirus* subgenus coronaviruses was lower and varied from 6 to 38%. Beyond sequence homology across sarbecoviruses, we observed the vaccine T-cell epitopes to be highly conserved between SARS-CoV-2 and SARS-CoV-1 and the 32 sarbecoviruses, reaching 75 to 100% homology (Tables S2 and S3). More in-depth analysis showed that among all CD8^+^ T-cell epitopes, 62% (n = 57) differed between SARS-CoV-2 and CoV-1 by at least one mutation, but these mutations did not affect HLA-Class I binding for a large majority of them (81%), as predicted by NetMHC4.0 (Table S2). Moreover, two CD4^+^ T-cell epitopes included in the vN2 sequence (N301-315, WPQIAQFAPSASAFF and N306-320, QFAPSASAFFGMSRI) and nine CD8^+^ T-cell epitopes from the vS2 and vN2 sequences (S1056-1063, APHGVVFL; S1089-1096, FPREGVFV; S1137-1145, VYDPLQPEL) and vN2 (N305-314, AQFAPSASAF; N306-315, QFAPSASAF; N307-315, FAPSASAFF; 308-317, APSASAFFGM; 310-319, SASAFFGMSR; 311-319, ASAFFGMSR) were 100% homologous across all sarbecoviruses (Table S5).These results confirm that vaccine sequences, particularly vS2 and vN2, are theoretically suitable for the design of a pan-sarbecovirus vaccine aimed at eliciting broad cross-reactive specific T-cell responses.

Next, we engineered plasmids expressing the vaccine sequence vRBD fused to the C-terminus of the Heavy (H) chain of anti-human CD40 humanized 12E12 IgG4 antibody, whereas the vN2, vS1, and vS2, sequences were fused sequentially to the Light (L) chain C-terminus to generate the CD40.CoV2 vaccine (Figure 1C). We have previously shown that 12E12 anti-CD40 fused to various viral antigens enhances CD40-mediated internalization and antigen-presentation by mononuclear cells and *ex-vivo* generated monocyte-derived DCs (Flamar et al., 2013; Yin et al., 2016).

### The CD40.CoV2 vaccine improves protection after viral challenge and induces cross-neutralizing antibody responses in the hCD40/K18-hACE2 transgenic mouse model

Transgenic mice expressing both the human (h) ACE2 receptor, the receptor of SARS-CoV-2 (Hoffmann et al., 2020), and hCD40 receptor (hCD40/K18-hACE2) were vaccinated with two intraperitoneal injections of CD40.CoV2 vaccine (10 μg) supplemented with polyinosinic-polycytidylic acid (Poly(IC) (50 μg) three weeks apart and challenged with Wuhan/D614G SARS-CoV-2 (Figure 2A). Poly(IC) was selected as an adjuvant due to its ability to increase antigen-presenting cell maturation (Cheng et al., 2017). In contrast to the vaccinated animals, controls exhibited significant weight loss from day 5 post-infection (pi), lasting until day 12 pi (Figure 2B). This was associated with the development of clinical symptoms in the controls from day 7 pi (Figure S1), leading to death of 67% of the animals by day 12 pi, whereas the vaccinated animals showed no symptoms and none died (Figures 2B and S1). Accordingly, the SARS-CoV-2 viral replication (genome equivalent/μg RNA) and viral infectious particles (PFU/mg of tissue) were lower in the lungs of the vaccinated mice than the controls or, indeed, undetectable (Figure 2C). We next assessed the antibody responses elicited *in vivo* by the CD40.CoV2 vaccine. One week after the booster injection (d28 post-vaccination [dpv]), the SARS-CoV-2 RBD- and S-specific IgG (Wuhan strain) binding levels were significantly higher in the vaccinated than mock-vaccinated mice (P = 0.0004 for both, Mann Whitney U test) (Figure 2D-2E). The CD40.CoV2 vaccine was also able to elicit RBD-specific IgG with cross-reactivity against VOCs (α, β, γ, δ) or the variant of interest (VOI) κ (P = 0.0004 for all comparisons between vaccinated and mock-vaccinated animals, Wilcoxon U test) (Figure 2D). Moreover, vaccine-elicited IgG highly cross-reacted with the SARS-CoV-1 spike protein (Figure 2E), but not the S protein of MERS or common cold coronaviruses, of which the sequences show less homology (Figure S2). By day 12 pi (corresponding to 40 dpv), cross-reactive IgG levels had increased in the control animals, but the response remained significantly lower than in the vaccinated animals (P = 0.0238 between vaccinated and mock-vaccinated animals, Wilcoxon U test) (Figure 2D-2E). Overall, the CD40.CoV2 vaccine elicited cross-neutralizing antibody responses against RBD from SARS-CoV-2 Wuhan and VOCs (Figure 2F) and S from both SARS-CoV-2 and SARS-CoV-1 (Figure 2G). These results were confirmed in a second replicate animal experiment (Figure S3).

**Figure 2.**
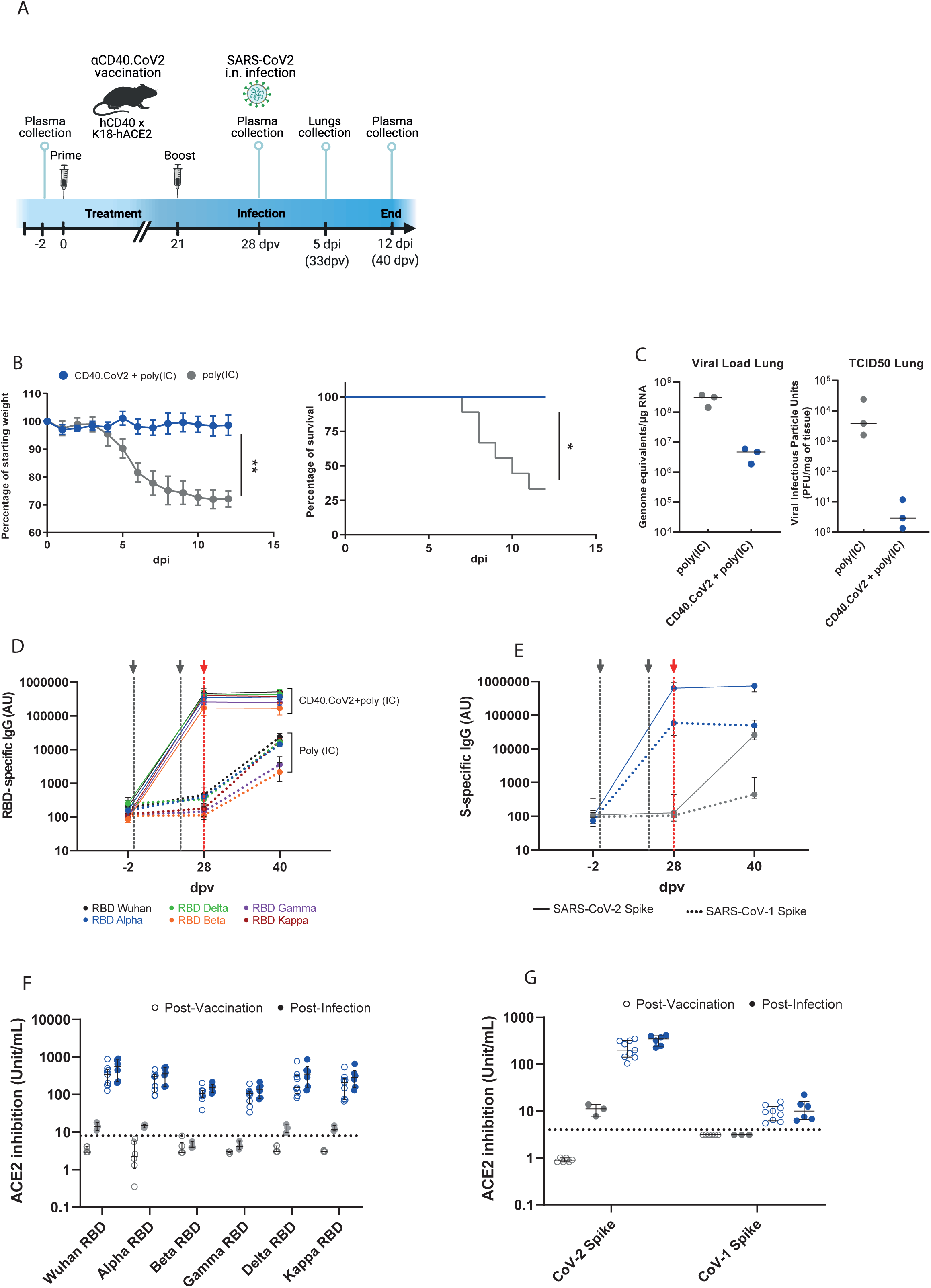
CD40.CoV2-vaccinated animals survive SARS-CoV-2 infection and show neutralizing and cross-reactive antibody responses. (A) Design of the CD40.CoV2 vaccination strategy before SARS-CoV2 infection. (B) Relative weight and survival of mock-vaccinated (grey) and vaccinated (blue) hCD40/K18-hACE2 transgenic mice after SARS-CoV-2 inoculation. Both parameters were recorded from days 0 to 12 post infection (pi). The mean ± SD is presented. A Mann-Whitney U test was conducted to compare differences in weight between the two groups on day 12 (n = 9-12 animals per group) (**P < 0.01). Kaplan-Meier survival curves were generated (n = 6-9 animals per group) and the P value was calculated using the log-rank (Mantel-Cox) test (*P < 0.05). (C) Viral load (genome equivalent/μg RNA) and viral infectious particle units (PFU/mg of tissue) in the lungs of mock-vaccinated (grey) and vaccinated (blue) hCD40/K18-hACE2 transgenic mice (n = 3 animals per group) on day 5 pi with the median plotted as a line. (D) Levels of IgG antibodies (AU) binding to Wuhan and VOCs SARS-CoV2 RBD proteins before vaccination (baseline, -2 days post-vaccination (dpv), n = 9-12 animals per group), after the completion of the vaccination schedule (28 dpv, n = 9-12 animals per group), and 40 dpv (i.e., day 12 pi time point, n = 3-5 animals per group). (E) Levels of IgG antibodies (AU) binding to SARS-CoV-2 (solid line) and SARS-CoV-1 (dashed line) S proteins in mock-vaccinated (grey) and vaccinated (blue) animals at -2, 28, and 40 dpv. Medians [Min-Max] are shown. The grey dashed lines represent prime and boost vaccines. The red dashed line represents SARS-CoV2 inoculation. Neutralizing activity of (F) anti-RBD antibodies (units/mL) and (G) anti-S antibodies (units/mL) in mock-vaccinated (grey) and vaccinated (blue) animals post-vaccination (open circles) and post-infection (solid circles). Medians ± Interquartile ranges (IQRs) are shown. Thirty plasma samples from unvaccinated mice were used to determine the threshold for positivity, defined as the whole units/mL value immediately above the concentration of the highest sample for RBD (i.e., 8 units/mL) and Spike (i.e., 4 units/mL) proteins. These results were reproduced in a second independent experiment (Figure S3).

In conclusion, the vaccine-elicited immune responses provided protection against SARS-CoV-2 challenge, with 100% survival, no clinical symptoms, and significant viral load control *in vivo.* These results significantly add to our previous vaccine studies in various animal models (hCD40transgenic mice, Hu-mice, or NHP), in which aCD40 targeting vaccines were able to induce potent humoral and cellular immune responses against Influenza virus, HIV, and, more recently, SARS-CoV-2 RBD (Flamar et al., 2018; Godot et al., 2020; Graham et al., 2016; Marlin et al., 2021).

### The CD40.CoV2 vaccine recalls cross-reactive functional SARS-CoV-2 T-cell responses *in vitro*

We next investigated the potency of the CD40.CoV2 vaccine in recalling T-cell responses *in vitro* using PBMCs collected from individuals who had experienced a viral infection, as previously demonstrated. Peripheral blood mononuclear cells (PBMCs) from 39 convalescent COVID-19 patients (between one to six months following infection) from the French COVID cohort (Yazdanpanah et al., 2021) were collected. The median [IQR] age of the patients was 56 [47-64], of whom 67% were male. First, we evaluated the frequency of CD4^+^- and CD8^+^- specific T cells, as assessed by the expression of the activation markers CD69^+^ and CD137^+^ (Figure 3A-3B) (Weiskopf et al., 2020). PBMCs (n = 5 donors) were stimulated with various doses of CD40.CoV2 vaccine, ranging from 10 to 10^-4^ nM, or an equimolar concentration of a combination of overlapping peptides (OLPs) spanning the full-length sequence of the vaccine antigens (vS1+vS2+vRBD+ vN2), referred as vOLPmix. The effective range of vaccine potency eliciting SARS-CoV-2-specific CD4^+^ T cells was between 1 to 10 nM, with maximal activity at 1 nM. At these concentrations, the recall of specific CD4^+^ T cells was 10-fold higher than with vOLPmix stimulation (Figure 3 B) (P = 0.0062, Wilcoxon U test). We observed a similar activation profile for CD8^+^ T cells, with the highest potency at 1 nM. Next, we confirmed the functionality of these cells, showing that the CD40.CoV2 vaccine induced robust and significantly higher proliferation of specific CD4^+^ and CD8^+^ T cells and CD19^+^ B cells than cells stimulated with vOLPmix or a control aCD40 vaccine fused to the Ebola glycoprotein (CD40.Gpz) (Figure 3C and 3D).

**Figure 3.**
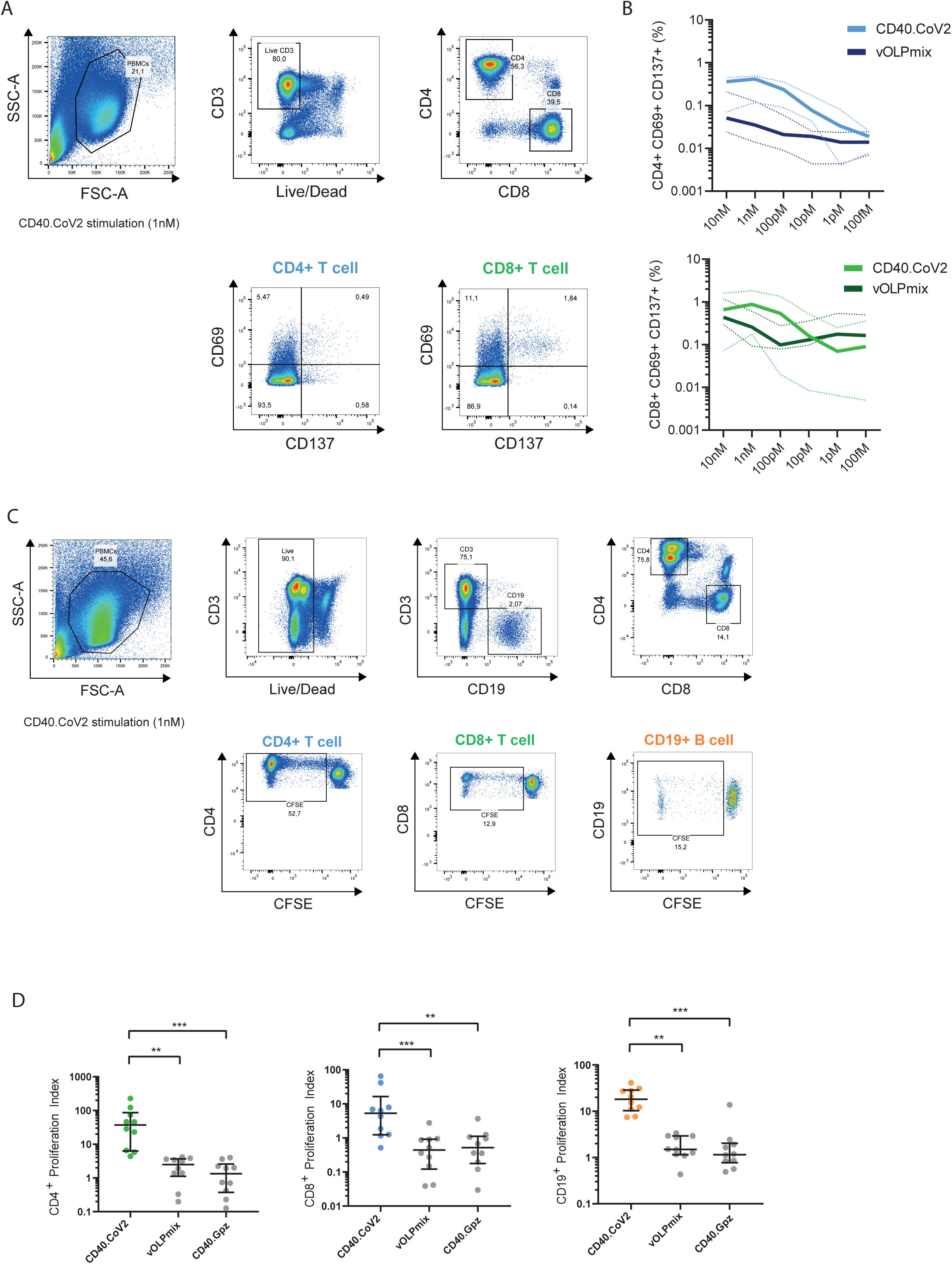
Determination of the optimal immunogenic CD40.CoV2 vaccine concentration and proliferation of specific T cells induced by the CD40.CoV2 vaccine. (A) Gating strategy for specific T cells that upregulate activation-induced markers (AIM). (B) Antigen specific activation of CD4^+^ (blue) and CD8^+^ (green) T cells from COVID-19 convalescent patients (n = 5) stimulated with various concentrations of CD40.CoV2 vaccine or a combination of OLPs covering the full-length sequence of the vaccine antigens (vOLPmix). Activation of SARS-CoV-2 specific CD4^+^ and CD8^+^ T cells is shown as the percentage of CD69^+^ CD137^+^ cells within the CD4^+^ or CD8^+^ subset after background subtraction. Median values (solid line) ± interquartile ranges (IQRs) (dashed lines) are shown. (C) Gating strategy for assessing the proliferation of specific T and B cells after seven days of CD40.CoV2 stimulation. (D) Proliferation of CD4^+^ T-cells, CD8^+^ T-cells, and B-cells from COVID-19 convalescent patients (n = 10) induced by the CD40.CoV2 vaccine, an irrelevant vaccine (αCD40 Gpz [1 nM]), or an equimolar concentration of vOLPmix. Data are expressed as a proliferation index obtained by dividing the frequency of proliferating cells after specific stimulation over background. Median values ± IQRs are shown. Friedman and Dunn’s multiple comparison tests were used for statistical analysis (**P < 0.01, ***P < 0.001).

These responses were likely favored by the targeting of vaccine epitopes through the anti-CD40 vehicle, as demonstrated by the broad and high levels of secretion of soluble factors produced by PBMCs from convalescent COVID-19 patients (n = 15) stimulated for two days with either the CD40.CoV2 vaccine (1 nM) or vOLPmix (Figure 4). Vaccine stimulation induced the production of chemokines involved in monocyte, macrophage, and DC chemotaxis (MCP-1, IP-10), as well as those associated with T-cell (CCL5) and neutrophil (IL-8) recruitment. Moreover, the level of cytokines produced by activated monocytes/macrophages and DCs (TNF, MIP-1α, MIP-1β, and IL-12p70) and those specific to cytotoxic activity (Granzyme B) also increased. Interestingly, Th1 (IFN-γ, IL-2), Th2 (IL-4, IL-13), and Th17 (IL-17A) cytokines were also detected. By contrast, stimulation with the matched vOLPmix only significantly induced the production of IL-2 (Figure 4).

**Figure 4.**
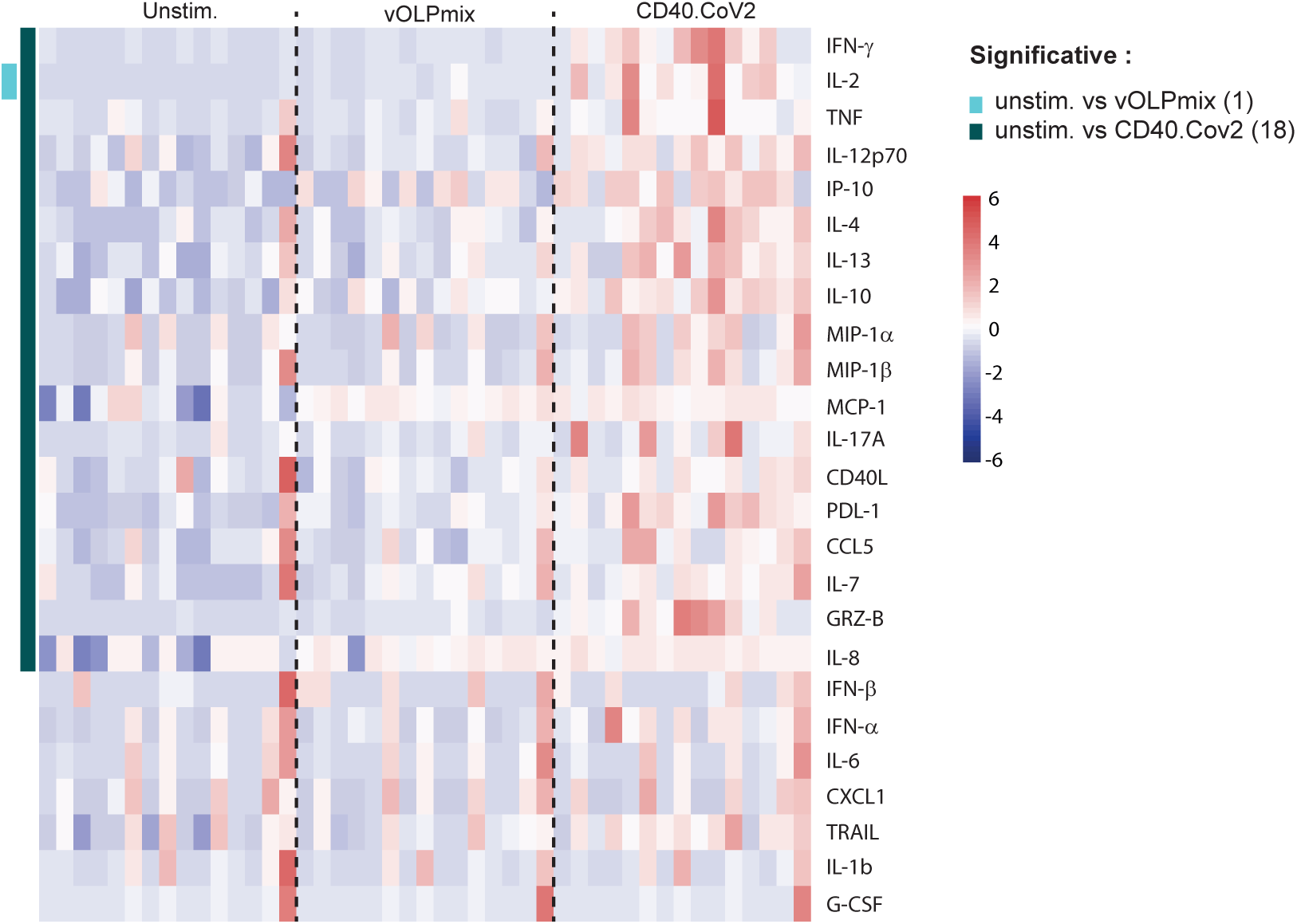
Heatmap of standardized biomarker expression in culture supernatants induced by the CD40.CoV2 vaccine. Supernatants from convalescent COVID-19 patient PBMCs collected on day 2 after stimulation with the CD40.CoV2 vaccine (1 nM) or an equimolar concentration of vOLPmix (n = 15). The colors represent standardized expression values centered around the mean, with variance equal to 1. Biomarker hierarchical clustering was computed using the Euclidean distance and Ward’s method (Ward, 1963).

Overall, these results demonstrate the potency of minute amounts of CD40.CoV2 vaccine to promote the recall of functional specific memory T and B cells.

### The CD40.CoV2 vaccine elicits multiepitope and cross-reactive specific T-cell responses against SARS-related sequences

We next confirmed the potency of the CD40.CoV2 vaccine construct to elicit cross-reactive functional memory T cells against individual vaccine regions from SARS-CoV-2 Wuhan or those harboring VOC/VOI mutations within these regions and SARS-CoV-1 S1/RBD/N2 epitopes.

First, PBMCs from convalescent COVID-19 patients (M1-M6 post-infection, n = 14) were stimulated with the CD40.CoV2 vaccine (1 nM) and restimulated on day8 either with one of the vOLPs (vRBD, vS1, vS2, or vN2) or control OLPs (cont.OLP), defined as SARS-CoV-2 regions either not contained in the CD40.CoV2 vaccine (SARS-CoV-2 Nucleocapsid [N1-N2] or SARS-CoV-2 Matrix [M]) or irrelevant peptides, such as Ebola glycoprotein (Gpz). The CD40.CoV2 vaccine recalled polyfunctional SARS-CoV-2-specific CD4^+^ T cells simultaneously producing up to three cytokines (IFN-γ ± IL-2 ± TNF) (Figure 5A-5C and Figure S4). Specific CD4^+^ and, to a lesser extent, CD8^+^ T cells produced IFN-γ against all vaccine antigens but not against control antigens (non-significant P value for control antigens vs the unstimulated condition, Wilcoxon U test). The CD40.CoV2 vaccine recalled polyepitope IFN-γ^+^ responses ranked from vS1 > vRBD > vN2 > vS2 for CD4^+^ T cells and vN2 > vRBD > vS1 for CD8^+^ T cells (Figure 5C). We also observed significant specific TNF^+^ and IL-2^+^ CD4^+^ T-cell responses against various vaccine antigens, whereas only the IL-2^+^ CD8^+^ T-cell response was significant after stimulation with vRBD (Figure S4). Interestingly, the strongest CD8^+^ T-cell response was directed against vN2, known to be important for long-term immunity (Lineburg et al., 2021) (Figure 5C). Finally, vaccine-expanded specific memory CD8^+^ T cells from three different patients showed cytotoxic activity against autologous cells pulsed with different vOLPs (vN2 or vRBD), whereas there was no cytotoxic activity when target cells were pulsed with cont.OLP (N1-N2 or M) (Figure 5D).

**Figure 5.**
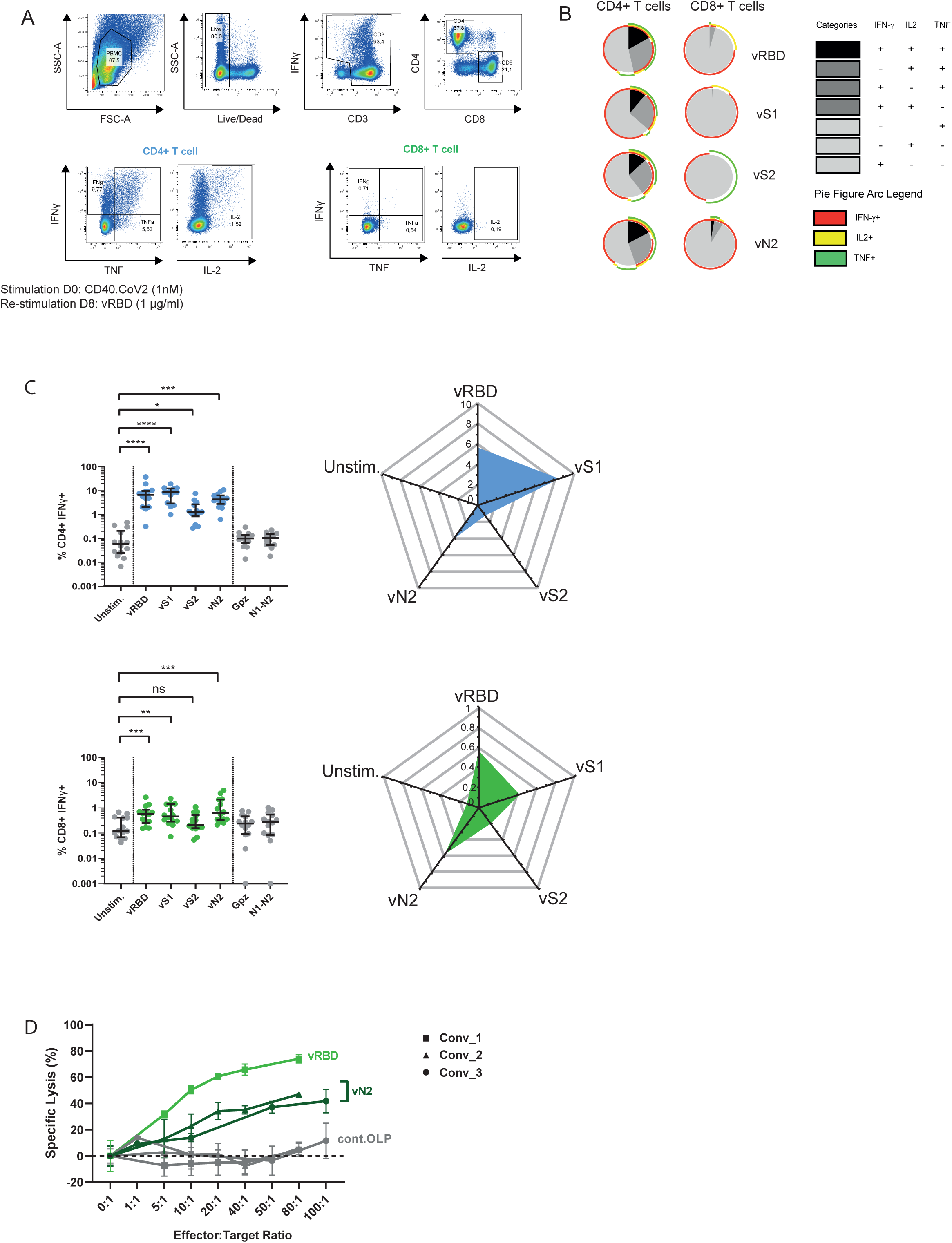
Polyfunctional and cytotoxic specific T-cell responses of convalescent COVID-19 patients after *in-vitro* stimulation with the CD40.CoV2 vaccine. (A) Representative dot plots of SARS-CoV-2-specific CD4^+^ and CD8^+^ T-cell responses after in vitro stimulation of patient PBMCs with the CD40.CoV2 vaccine (1 nM) on D0 and re-stimulation with various vOLPs (vRBD, vS1, vS2 or vN2) (1 µg/ml) on D8. (B) Functional composition of SARS-CoV-2-specific CD4^+^ and CD8^+^ T-cell responses induced by the CD40.CoV2 vaccine and various vOLPs (vRBD, vS1, vS2 or vN2) (1 µg/ml). Responses are color coded according to the combination of cytokines produced. The arcs identify cytokine-producing subsets (IFN-γ, IL-2, and TNF) within the CD4^+^ or CD8^+^ T cell populations. (C) Frequency and radar charts of the merged median of IFN-γ^+^ SARS-CoV-2-specific CD4^+^ (blue) or CD8^+^ (green) T cells from convalescent COVID-19 patients (n = 14) stimulated or not with the CD40.CoV2 vaccine (1 nM) on D0 and re-stimulated with various vOLPs (vRBD, vS1, vS2 or vN2), cont.OLP (SARS-CoV2 N1-N2, or Ebola Gpz) (grey) on D8 (1 µg/mL). Median values ± IQRs are shown. Friedman and Dunn’s multiple comparison tests were used for statistical analysis (*P < 0.05, **P < 0.01, ***P < 0.001, ****P < 0.0001, ns: not significant). (D) Specific lysis of CD8^+^ T cells stimulated with the CD40.CoV2 vaccine (1 nM) against autologous PHA-blasted PBMCs from three different convalescent COVID-19 patients, pulsed with either vRBD (light green), vN2 (dark green), or cont.OLP (SARS-CoV2 N1-N2 or M) (grey). The means of triplicate values ± the standard deviation (SD) are shown. Each symbol represents a different patient.

We also observed that CD40.CoV2 vaccine-expanded *in-vitro* CD4^+^ and CD8^+^ T-cell responses were not affected when the cells were stimulated with RBD OLPs containing common mutations of β/γ, δ (VOCs) and κ (VOI) (Figure 6A). The cross-reacting CD8^+^ and CD4^+^ T-cell responses were polyfunctional, simultaneously producing up to 2 or 3 cytokines, respectively, with no major differences between VOCs (Figure 6B).

**Figure 6.**
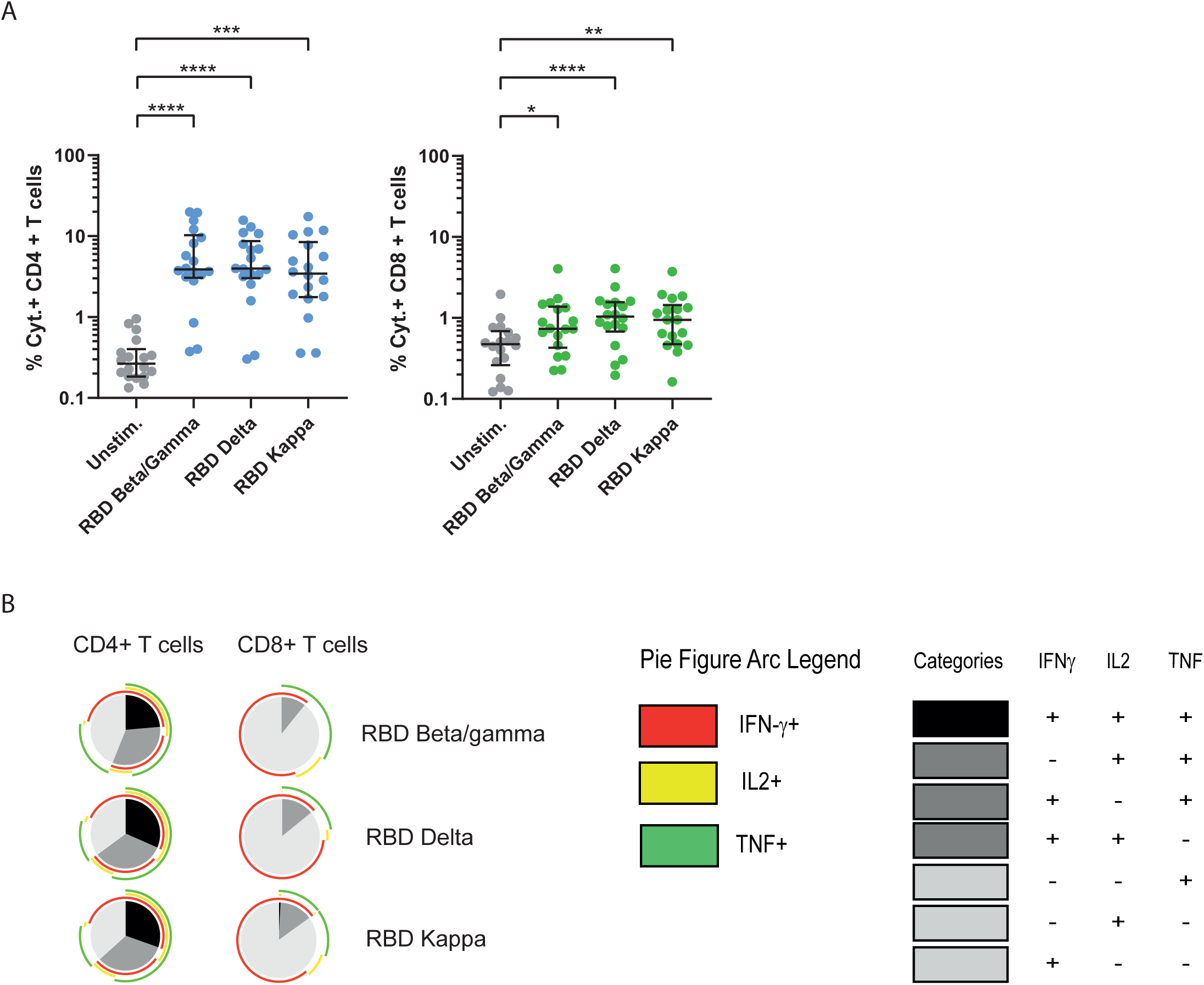
Polyfunctional specific T-cell responses against SARS-CoV-2 VOCs after stimulation with the CD40.CoV2 vaccine. (A) Frequency of total cytokines (IFN-γ ± IL-2 ± TNF) produced by specific CD4^+^ (blue) or CD8^+^ (green) T cells from convalescent COVID-9 patients (n = 18) after i*n-vitro* stimulation with the CD40.CoV2 vaccine (1 nM) on D0 and re-stimulation with RBD OLP from various VOCs or VOI (1 µg/mL). (B) Functional composition of SARS-CoV-2-specific CD4^+^ and CD8^+^ T-cell responses induced by the CD40.CoV2 vaccine against VOCs/VOI. Responses are color coded according to the combination of cytokines produced. The arcs identify cytokine-producing subsets (IFN-γ, IL-2, and TNF) within the CD4^+^ and CD8^+^ T-cell populations. Median values ± IQRs are shown. Friedman’s test was used for comparisons (*P < 0.05, **P < 0.01, ***P < 0.001, ****P < 0.0001).

Finally, we re-stimulated CD40.CoV2-stimulated PBMCs from convalescent COVID-19 patients with OLPs covering the S1 (S1-CoV-1), vRBD (vRBD-CoV-1), and vN2 (vN2-CoV-1) regions from SARS-CoV-1 and S1 from MERS. The vaccine elicited a high frequency of cross-reactive SARS-CoV-1 CD4^+^ and CD8^+^ T cells and, to a lower extent, cross-reactive MERS-S1 CD8^+^ T cells (Figure 7). Interestingly, the CD40.CoV2 vaccine-induced SARS-CoV-1- and SARS-CoV-2-specific T-cell responses were highly correlated for all corresponding antigen sequences (Figure S5). Overall, we confirm that the breadth of recall responses induced by the CD40.CoV2 vaccine is not affected by RBD mutations from SARS-CoV-2 VOCs and fully recognizes SARS-CoV-1 epitopes.

**Figure 7.**
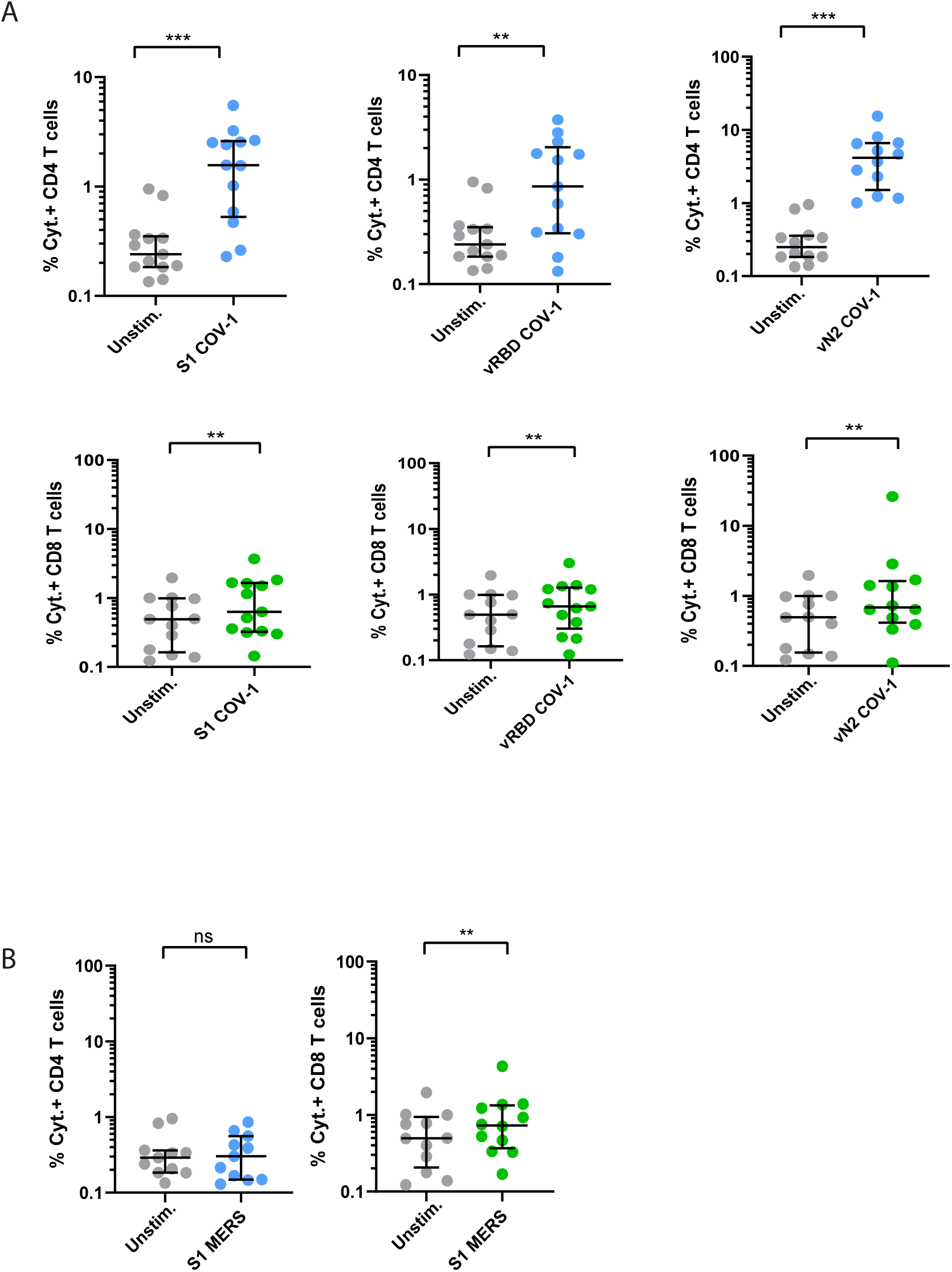
Cross-reactive specific T-cell responses against SARS-CoV-1 and MERS of convalescent COVID-19 patients after *in-vitro* stimulation with the CD40.CoV2 vaccine. Frequency of total cytokines (IFN-γ ± IL-2 ± TNF) produced by specific CD4^+^ (blue) or CD8^+^ (green) T cells after *in-vitro* stimulation with the CD40.CoV2 vaccine (1 nM) on D0 and re-stimulation with OLPs representing the sequences of S1, vRBD, and vN2 from (A) SARS-CoV-1 and (B) S1 from MERS (1 µg/mL). Median values ± IQRs are shown. The Wilcoxon U test was used for comparisons (**P < 0.01, ***P < 0.001, ns: not significant).

## Discussion

Despite the rapid development of several effective vaccines against SARS-CoV-2, recent observations from vaccine campaigns in the general population have shown that the antibody response is waning, with reduced efficacy against VOCs. Most are characterized by mutations found in areas that are likely targeted by neutralizing antibodies, leading to vaccine escape and compromising the first line of immunological defense against SARS-CoV-2. Moreover, whether the current first-generation vaccines based on the original virus strain would still protect against emerging VOCs or pre-emergent coronaviruses, which may be responsible for future pandemics, is unknown. Thus, the development of new-generation vaccines that can induce B- and T-cell responses to a broad range of epitopes, less prone to variation, are warranted.

Here, we demonstrate the immunogenicity and anti-viral efficacy of a new protein vaccine composed of three regions from S (aa 125-250, 318-541, 1056-1209) and one from N (aa 276-411) of SARS-CoV-2, accumulating a large set of predicted CD4^+^ and CD8^+^ T- and B-cell epitopes that are highly homologous to those of SARS-CoV-1 and 32 recently described SARS-CoV-2-related coronaviruses. The CD40.CoV2 vaccine elicited potent SARS-CoV-2- specific cross-reactive and neutralizing antibodies associated with anti-viral and protective activity against SARS-CoV-2 challenge in the hCD40/K18-hACE2 mouse model. Furthermore, vaccinated mice developed high neutralizing antibody levels against the RBD region from not only SARS-CoV-2 Wuhan but also several SARS-CoV-2 VOCs/VOI (α, β, γ, δ and κ) and S from SARS-CoV-1. These results confirm and extended our previous observations of the antiviral efficacy of the CD40.RBD vaccine in convalescent macaques (Marlin et al., 2021).

Our vaccine design was also driven by the need to include conserved epitopes to generate robust memory CD4^+^ or CD8^+^ T cells to provide early control of acute infection with a novel SARS-CoV-2 VOC or closely related virus in the absence of pre-existing cross-protective antibodies (Tarke et al., 2021a). The *in-silico* definition of vaccine sequences was supported by several observations. First, various CD40.CoV2 vaccine epitopes have already been shown by others, through structure-based network analysis and assessment of HLA class-I peptide stability, to be structurally constrained, thus limiting genetic variation across SARS-CoV-2, SARS-CoV-1, and sarbecoviruses (Nathan et al., 2021). For example, the CD40.CoV-2 vaccine contains 6/28 S (S319-329, RVQPTESIVRF; S321-329, QPTESIVRF; S386-395, KLNDLCFTNV; S386-396, KLNDLCFTNVY; S515-524, FELLHAPATV; S1093-1102, GVFVSNGTHW) and 6/11 N (N276-286, RRGPEQTQGNF; N305-313, AQFAPSASA; N306-314, QFAPSASAF; N306-315, QFAPSASAFF; N308-315, APSASAFF; N308-317, APSASAFFGM) of highly networked constrained regions with stabilizing CD8^+^ T-cell epitopes with global HLA coverage (Nathan et al., 2021). Second, we showed the reactivity of CD4^+^ and CD8^+^ T cells to nanomolar concentrations of the CD40.CoV2 vaccine using an approach combining the expression of activation markers, cytokine production, T-cell proliferation, and cytotoxic function. These results demonstrate the immunogenicity of vaccine epitopes and are consistent with those of prior studies describing a number of them in several patient cohorts (Tables S3 and S4) (Grifoni et al., 2020, 2021; Le Bert et al., 2020; Li et al., 2006). Third, based on the recent review of all SARS-CoV-2 CD4^+^ and CD8^+^ T-cell epitopes reported in 25 studies (Grifoni et al., 2021), it appears that the CD40.CoV2 vaccine contains 42% of the described immunodominant CD8^+^ T-cell epitopes for S and 39% for N. The respective values are 54% (S) and 35% (N) for the immunodominant CD4^+^ T-cell epitopes. Moreover, all vaccine regions contain dominant epitopes. For example, the vS1 and vRBD regions of the CD40.CoV2 vaccine closely fit or overlap the two immunodominant S regions for CD4^+^ T cells (S154-254, S296-370). Similarly, the vN2 sequence overlaps the described CD4^+^ and CD8^+^ T-cell immunodominant region of the nucleocapsid (201-371) (Grifoni et al., 2021).

A crucial question for the development of vaccines to counteract the escape of the virus from neutralizing antibodies is whether SARS-CoV-2 VOCs can evade T-cell immunity. However, even if SARS-CoV-2 does mutate, analysis of mutations associated with the various VOCs shows the vast majority of CD40.CoV2 vaccine epitopes to be conserved in SARS-CoV-2 variants (Tables S2 and S3). Accordingly, we found that *in-vitro* stimulation with the CD40.CoV-2 vaccine elicited specific cross-reactive polyfunctional CD4^+^ and CD8^+^ T-cell responses against RBD from VOCs/VOI. Similarly, we found that the breadth of vaccine-elicited CD4^+^ and CD8^+^ T-cell responses extended to S1, RBD, and N sequences from SARS-CoV-1. Indeed, expanded CD8^+^ T cells were even cross reactive to S1 peptides from MERS, despite low homology with SARS-CoV-2 (15%).

While it is critical to determine to what extent VOCs may or may not be susceptible to evading existing humoral responses, T-cell associated immunity is, in general, significantly more difficult for viruses to overcome, due to the broad and adaptable response generated in a given individual and because of the variety of HLA haplotypes. In this regard, the new SARS-CoV-2 B.1.1.529 (Omicron) VOC, which emerged in November 2021, is characterized by the presence of 32 mutations in Spike, located mostly in the N-terminal domain (NTD) and the RBD. Recent results have shown that this VOC significantly escapes from neutralizing antibodies, either therapeutic or those from convalescent or vaccinated individuals at various levels (Cele et al., 2021; Planas et al., 2021; Rössler et al., 2021). However, preliminary studies have shown minimal cross-over between mutations associated with the Omicron variant of SARS-CoV-2 and CD8^+^ T-cell epitopes identified in convalescent COVID-19 individuals (Redd et al., 2021). Of note, the vN2 vaccine sequence is 100% homologous with that of most VOCs, including the Omicron variant. Overall, concerning the objective to develop a vaccine with a broader range of protection, our results confirm previous observations that single amino-acid substitutions or deletions across large peptidomes do not significantly affect polyclonal memory T-cell responses (Tarke et al., 2021b).

The design of the CD40.CoV-2 vaccine benefited from the high number of genetically conserved SARS-CoV-2 S and N sequences across VOCs, as well as those of SARS-CoV-related viruses, with the goal of inducing broad immune cross reactivity, a key component for the development of a next-generation pan-sarbecovirus vaccine. We show that the vaccine T-cell epitopes are highly conserved with those of SARS-CoV-2 VOCs, SARS-CoV-1, and, more generally, all 38 sarbecoviruses tested, with up to 80-100% homology for the most highly conserved T-cell epitopes (Tables S2 and S3). Moreover, we show that nine CD8^+^ T-cell epitopes, from S and N, contained in the vaccine are 100% homologous among all sarbecoviruses (Table S5). Globally, these results confirm that vaccine sequences, particularly vS2 and vN2, are theoretically suitable for the design of a pan-sarbecovirus vaccine aiming to elicit broad cross-reactive T-cell responses.

We further propose in this vaccination strategy to deliver the antigens through a DC-targeting platform, as the antigens were fused to a humanized anti-CD40 monoclonal antibody. This platform has already been tested *in vitro*, in various preclinical animal models, and is currently in phase I/II clinical development for a prophylactic HIV vaccine (NCT04842682). Thus, by targeting epitopes of the S and N proteins, the CD40.CoV2 vaccine may represent an excellent booster of pre-existing immunity, induced either by previous priming with available vaccines or by natural infection, as we recently demonstrated that a single dose of the CD40.RBD vaccine, injected without adjuvant, is sufficient to elicit neutralizing antibodies that protect macaques from a new viral challenge (Marlin et al., 2021).

Because SARS-CoV2 humoral responses decline rapidly over time, repeated vaccinations at short time intervals are required to maintain high neutralizing responses, at least with the currently available vaccines, which all target the S protein (Barda et al., 2021). However, aside from antibody responses, early induction of functional SARS-CoV-2-specific T cells is observed in patients with mild disease and rapid viral clearance (Tan et al., 2021). In addition, recent studies have highlighted the significant cross-protective advantage of a heterologous boost, even if the vaccine antigens do not fully match the viral challenge (Dangi et al., 2021) and the interest to target also conserved regions of the spike protein, outside the RBD domain, for induction of cross-neutralizing antibodies (Cameroni et al.). Thus, a significant advantage of our vaccine may be to extend the breadth of the responses of current vaccines.

Our study had several limitations. These included the absence of characterization of cross-neutralizing antibodies in vaccinated mice against the B.1.1.529 Omicron variant, which emerged during the writing of this manuscript. However, we could expect that our vaccine would elicit T-cell responses against N epitopes because of the high homology (100%) between vaccine sequences and this VOC. Due to the limited availability of hCD40/K18-hACE2 mice, we did not evaluate various VOC challenges in mice. Finally we favored the analysis of T cell responses using samples from recovered individuals instead of *in vivo* preclinical models. Although these responses may be dependent on the “clinical history” of patients and their HLA haplotypes, they are less biased than those that would be observed in an animal model. Our results show that the *in-vitro* vaccine responses are directed against all vaccine proteins, which confirmed the broad HLA coverage of the vaccine sequences.

In conclusion, it is becoming urgent to develop a “pan-sarbecovirus vaccine”. The development of a new protein-based vaccine with expected improved tolerability suitable for people with specific vulnerabilities and children would extend the portfolio of current vaccines and be instrumental in controlling the circulation of the virus and the emergence of new variants. By selecting a narrow range of immunodominant epitopes, presented by a wide variety of HLA alleles and less prone to genetic variations across sarbecoviruses, we provide a rationale for the development of a global T cell-based vaccine to counteract emerging SARS-CoV-2 variants and future SARS-like coronaviruses.

## Acknowledgments

We thank the patients who donated their blood. We thank F. Mentre, S. Tubiana, the French COVID cohort, and REACTing (REsearch & ACtion emergING infectious diseases) for cohort management. We thank the scientific advisory board of the French COVID-19 cohort composed of Dominique Costagliola, Astrid Vabret, Hervé Raoul, and Laurence Weiss. We thank Corinne Krief, Lydia Guillaumat, and Marie Déchenaud for the biobanking of convalescent COVID-19 patient samples. We also thank Cathleen Lutz and The Jackson Laboratory for providing the K18-hACE2 mice and Pr. Sylvie van der Werf, Dr. X.Lescure, and Pr. Y. Yazdanpanah for the BetaCoV/France/IDF0372/2020 strain. The following reagents were obtained through BEI Resources, NIAID, NIH: Peptide Array, SARS Coronavirus Nucleocapsid (N) Protein, NR-52419, and Peptide Array, SARS Coronavirus Spike (S) Protein, NR-52418.

## Author Contributions

Y.L, V.G, and S.C conceived and designed the study. S.Co, A.W, M.S, C.L, A.Z, B.M, Q.B, M.F, S.H, S.C, G.Z, S.Z, V.G, and Y.L analyzed and interpreted the data. S.Co, M.S, L.D, F.P, A.S, C.P, Q.B, M.F, S.S and P.H performed the experiments. S.Z, Z.W, S.C, C.L, M.S., and Y.L. designed and produced the CD40.CoV2 vaccine. A.Z, B.M, and V.G supervised the animal studies. J.G and M.C.G participated in sample and clinical data collection. M.C administered the project. Y.L, V.G, S.Co, AW, M.S, A.Z, S.Z, and G.Z drafted the first version and wrote the final version of the manuscript. All authors approved the final version.

## STAR Methods

### Resource availability

#### Lead contact

Further information and requests for resources should be directed to and will be fulfilled by the lead contact, Yves Lévy.

#### Materials availability

The CD40.CoV2 vaccine generated in this study was deposited in GenBank: anti-human CD40 12E12 antibody IgG4 H chain (GenBank ID: AJD85779.1 residues 20-467) fused to SARS_CoV_2RBD (GenBank ID: UEP92470.1 residues 17-240) followed by EPEA (C-tag) and the anti-human CD40 12E12 antibody kappa L chain (GenBank ID: AJD85780.1 residues 21-236) fused sequentially to a linker (GenBank ID: AJD85777.1 residues 699-725), nucleocapsid phosphoprotein, partial [Severe acute respiratory syndrome coronavirus 2] (GenBank ID: QWE63393.1 residues 95-230), linker residues AR, Chain A, Spike protein S1 [Severe acute respiratory syndrome coronavirus 2] (GenBank ID: 7M8J_A residues 113-237), linker residues TR, and Sequence 12 from patent US 8518410 (Genbank ID: AGU17682.1 residues 3-27), surface glycoprotein, partial [Severe acute respiratory syndrome coronavirus 2] (GenBank ID: UET03776.1 residues 195-348)

#### Data and code availability

Any additional information required to reanalyze the data reported in this paper is available from the lead contact upon request.

### Experimental model and subject details

#### Animals

Animal housing and experimental procedures were conducted according to the French and European Regulations (*Parlement Européen et du Conseil du 22 septembre 2010, Décret n° 2013-118 du 1er février 2013 relatif à la protection des animaux utilisés à des fins scientifiques*) and the National Research Council Guide for the Care and Use of Laboratory Animals (*National Research Council (U.S.), Institute for Laboratory Animal Research (U.S.), and* National Academies Press (U.S.), Eds., Guide for the care and use of laboratory animals, 8th ed. Washington, D.C: National Academies Press, 2011). The animal BSL3 facility is authorized by the French authorities (Agreement N° B 13 014 07). All animal procedures (including surgery, anesthesia, and euthanasia, as applicable) used in the current study were submitted to the Institutional Animal Care and Use Committee of the CIPHE approved by the French authorities (CETEA DSV – APAFIS#26484-2020062213431976 v6). All CIPHE BSL3 facility operations are overseen by a biosecurity/biosafety officer and accredited by the Agence Nationale de Sécurité du Médicament (ANSM). Heterozygous K18-hACE C57BL/6J mice (strain: 2B6.Cg-Tg (K18-ACE2)2Prlmn/J) were obtained from The Jackson Laboratory. The hCD40-OST transgenic mice expressed a human *Cd40* gene in place of the mouse *Cd40* gene. They were derived at CIPHE under CIPHE-Sanofi Research Collaborative program n° 171137A10 and kindly provided by Sanofi under the agreement MTA #209012. All breeding, genotyping, and production of hCD40/K18-hACE2 was performed at the CIPHE. Animals were housed in groups and fed standard chow diets.

#### COVID-19 convalescent patients

We enrolled a subgroup of COVID-19 patients of the prospective French COVID cohort (registered at clinicaltrials.gov NCT04262921) in this study. Ethics approval was given on February 5, 2020, by the French Ethics Committee CPP-Ile-de-France VI (ID RCB: 2020-A00256-33). Eligible patients were those who were hospitalized with virologically confirmed COVID-19. Convalescent follow-up visits were performed between one, three, and six months after infection. The study was conducted with the understanding and consent of each participant or their surrogate covering the sampling, storage, and use of biological samples.

### Method details

#### Cloning and production of the CD40.CoV2 vaccine

The vaccine was produced using the expression plasmids described in materials availability via transient transfection (TransIT-PRO® Transfection Kit, Mirus) into mammalian CHO-S cells (ThermoFisher) followed by Protein A-affinity purification (Ceglia et al., 2021; Cheng et al., 2018). The eluted product (2.2 mg/ml, 0.3 ng LPS/mg) was stored at -80°C in PBS with 125 mM hydroxypropyl β-cyclodextrin (Cavitron W7 HP5). CD40 binding was validated by ELISA as previously described (Flamar et al., 2013).

#### Wuhan/D614 SARS-CoV-2 virus production

Vero E6 cells (CRL-1586; American Type Culture Collection) were cultured at 37°C in Dulbecco’s modified Eagle’s medium (DMEM) supplemented with 10% fetal bovine serum (FBS), 10 mM HEPES (pH 7.3), 1 mM sodium pyruvate, 1X non-essential amino acids, and 100 U/ mL penicillin–streptomycin. The strain BetaCoV/France/IDF0372/2020 was supplied by the National Reference Centre for Respiratory Viruses hosted by the Institut Pasteur (Paris, France). The human sample from which strain BetaCoV/France/IDF0372/2020 was isolated was provided by the Bichat Hospital, Paris, France. Infectious stocks were grown by inoculating Vero E6 cells and collecting supernatants upon observation of the cytopathic effect. Debris was removed by centrifugation and passage through a 0.22-μm filter. Supernatants were stored at -80°C.

#### Vaccination and infection of hCD40/K18-hACE2 transgenic mice

Mice of 8 to 12 weeks of age of both sexes received two intraperitoneal injections of the CD40.CoV2 vaccine (10 µg) plus polyinosinic-polycytidylic acid (Poly-IC; Oncovir) (50 µg) or poly(IC) alone three weeks apart. Mice were further infected with Wuhan/D614 SARS-CoV-2 at week 4. Vaccinated and mock-vaccinated mice were administered 2.5 x 10^4^ PFU of SARS-CoV-2 via intranasal administration. Mice were monitored daily for morbidity (body weight) and mortality (survival). During the monitoring period, mice were scored for clinical symptoms (weight loss, eye closure, appearance of the fur, posture, and respiration). Mice obtaining a clinical score defined as reaching the experimental end-point were humanely euthanized. Two independent experiments of CD40.CoV2 vaccination followed by SARS-CoV2 inoculation were performed. Blood was collected on day -2 (before vaccination), day 28 (before viral infection), and day 40 post-vaccination.

The clinical and immunological monitoring of Experiment 1 are reported in the manuscript and principal figures. The antibody responses monitored in Experiment 2 to confirm the main results are presented in Supplemental Figure S3.

#### Measurement of SARS-CoV-2 viral load by RT-qPCR and TCID50 (50% of tissue-culture infective dose)

For viral titration by RT-qPCR, tissues were homogenized with ceramic beads in a tissue homogenizer (Precellys – Bertin Instruments) in 0.5 mL RLT buffer. RNA was extracted using the RNeasy Mini Kit (QIAGEN) and reverse transcribed using the High-Capacity cDNA Reverse Transcription Kit (Thermo Fisher Scientific). Amplification was carried out using OneGreen Fast qPCR Premix (OZYME) according to the manufacturer’s recommendations. The number of copies of the SARS-CoV2 RNA-dependent RNA polymerase (RdRp) gene in samples was determined using the following primers: forward primer – catgtgtggcggttcactat, reverse primer – gttgtggcatctcctgatga. This region was included in a cDNA standard to allow the copy number determination down to ∼100 copies per reaction. The copies of SARS-CoV2 were compared and quantified using a standard curve and normalized to total RNA levels. An external control (mock-infected wildtype animal, nondetectable in the assay) and a positive control (SARS-CoV-2 cDNA containing the targeted region of the RdRp gene at a concentration of 10^4^ copies/µl [1.94 x 10^4^ copies/µl detected in the assay]) were used in the RT-qPCR analysis to validate the assay. The median tissue-culture infectious dose (TCID50) represents the dilution of a virus-containing sample at which half of the inoculated cells show signs of infection. To perform the assay, lung tissue was weighed and homogenized using ceramic beads in a tissue homogenizer (Precellys – Bertin Instruments) in 0.5 ml RPMI media supplemented with 2% FCS and 25 mM HEPES. Tissue homogenates were then clarified by centrifugation and stored at −80°C until use. Forty-thousand cells per well were seeded in 96-well plates containing 200 µl DMEM + 4% FCS and incubated for 24 h at 37°C. Tissue homogenates were serially diluted (1:10) in RPMI media and 50 µl of each dilution was transferred to the plate in six replicates for titration at five-days post-inoculation. Plates were read for the CPE (cytopathology effect) using microscopy reader and the data were recorded. Viral titers were then calculated using the Spearman-Karber formula and expressed as TCID50/mg of tissue.

#### Antibody measurement

Two multiplexed MesoScale Discovery immunoassays (V-PLEX Coronavirus Panel 3 and V-PLEX SARS-CoV-2 Panel 11 [IgG] Kits, MesoScale Discovery, Rockville, MD, USA) were used on all available plasma samples to measure plasma IgG antibodies to SARS-CoV-2, SARS-CoV, MERS-CoV, and HCoVs. Coronavirus Panel 3 plates are MULTI-SPOT 96-well, 10 Spot, coated with three SARS-CoV-2 antigens (spike, receptor binding domain [RBD], and nucleocapsid), and spike proteins from SARS-CoV, MERS-CoV, and seasonal HCoVs OC43, HKU1, 229E, and NL63. SARS-CoV-2 Panel 11 plates are MULTI-SPOT 96-well, 10 Spot, coated with RBD proteins from various SARS-CoV-2 lineages: Wuhan; Alpha; Beta and Botswana; Gamma; Delta sub-lineages and Vietnam; Epsilon, California, and New York; Eta, Iota, India, Zeta, and Kentucky; New York; U.K. and Philippines; Kappa and India. Assays were performed according to the manufacturer’s instructions, with samples diluted 1:50 000. The electro-chemiluminescence (ECL) signal was recorded and the results are expressed as arbitrary units (AU).

Two alternative immunoassays (V-PLEX Coronavirus Panel 3 and V-PLEX SARS-CoV-2 Panel 11 [ACE-2] Kits, MesoScale Discovery) were used to measure the ability of mouse plasma samples to inhibit angiotensin-converting enzyme 2 (ACE2) binding to the Spike protein of different coronaviruses and different variants of SARS-CoV-2 RBD proteins. The assays were performed according to the manufacturer’s instructions with samples diluted 1:33 to 1:3333. Antibody concentrations were quantified using a reference standard (ACE2 Calibration Reagent) and are expressed as units/mL (one unit per mL concentration of ACE2 Calibration Reagent corresponds to neutralizing activity of 1 µg/mL monoclonal antibody to SARS-CoV-2 Spike protein). For V-PLEX Coronavirus Panel 3, the lower limit of quantification (LLOQ) was calculated to be between 0.68 and 0.82 unit/mL for Spike CoV-2 and between 3.10 and 3.24 units/mL for Spike CoV-1. For V-PLEX SARS-CoV-2 Panel 11, the LLOQ was calculated to be between 2.83 and 2.94 units/mL for RBD Wuhan, 2.43 and 3.20 units/mL for RBD Alpha, 2.09 and 2.87 units/mL for RBD Beta, 2.56 and 2.99 units/mL for RBD Gamma, 2.1 and 3.02 units/mL for RBD Delta, and between 2.51 and 3.12 units/mL for RBD Kappa. Values under the LLOQ were imputed at the LLOQ. Thirty plasma samples from unvaccinated mice were used to determine the threshold for positivity, defined as the whole units/mL value immediately above the concentration of the highest sample for RBD (i.e., 8 units/mL) and Spike (i.e., 4 units/mL) proteins.

#### Specific antigens

Various peptide pools from reference strain Human 2019-nCoV HKU-SZ-005b, from JPT Peptide Technologies (Berlin, Germany) or BEI Resources, were used, as mentioned, for the various assays. A set of four pools of 15-mer peptides, overlapping by 11 amino acids, covering the four regions of S and N sequences of SARS-CoV2 included in the CD40.CoV2 vaccine: vS1 (29 peptides), vRBD (54 peptides), vS2 (37 peptides), and vN2 (32 peptides). In certain experiments, vS1, vRBD, vS2, and vN2 were pooled to provide a combination of all sequences included in the CD40.CoV2 vaccine (vOLPmix). A pool of 54 peptides (15-mers overlapping by 11 amino acids) encompassing the three RBD mutations K417N (four peptides), E484K (three peptides), and N501Y (four peptides): RBD SARS-CoV-2 beta/gamma. Two PepMix pools of RBD SARS-CoV-2 delta and RBD SARS-CoV-2 kappa (15-mer peptides, overlapping by 11 amino acids). A set of two pools of 15/20-mer peptides, overlapping by 10 amino acids, covering the two regions of RBD and N sequences of SARS-CoV-1 corresponding to the CD40.CoV2 vaccine sequences: vRBD-CoV1 (30 peptides), vN2-CoV-1 (18 peptides). A pool of 15-mer peptides, overlapping by 11 amino acids, covering the S1 region sequence of SARS-CoV-1: S1-CoV-1 (156 peptides). A pool of 15-mer peptides, overlapping by 11 amino acids, covering the S1 region sequence of MERS: S1-MERS (168 peptides). As controls (cont.OLP), we used a set of two pools of 15-mer peptides, overlapping by 11 amino acids, of N and M sequences of SARS-CoV2 not included in the vaccine: N1-N2 (34 peptides) and M (53 peptides) or an irrelevant pool of overlapping 15-mer peptides (11-amino acid overlaps) from the Ebola virus Mayinga variant glycoprotein (Gpz: 77 peptides).

#### Quantification of culture supernatant analytes

We quantified 25 analytes in supernatants from convalescent COVID-19 PBMCs on day 2 after CD40.CoV2 vaccine (1 nM) or vOLP (equimolar concentration) stimulation using the Human XL Cytokine Luminex® Performance Panel Premixed Kit: CCL2/MCP-1, CCL3/MIP-1α, CCL4/MIP-1β, CCL5/RANTES, CD40 Ligand/TNFSF5, CXCL1/GROα, CXCL10/IP-10, GCSF, Granzyme B, IFN-α, IFN-β, IFN-γ, IL-1β, IL-2, IL-4, IL-6, IL-7, IL-8/CXCL8, IL-10, IL-12 p70, IL-13, IL-17/IL-17A, PD-L1/B7-H1, TNF, and TRAIL/TNFSF10 (R&D Systems/Bio-Techne), according to the manufacturers’ instructions.

#### Characterization of SARS-COV-2-specific immune responses in convalescent COVID-19 patients

Cellular responses to CD40.CoV2 vaccine were assessed using the Activation Induced Marker assay (AIM) and EpiMax technology (Chujo et al., 2013). For the AIM assay, PBMCs were stimulated *in vitro* with various concentrations of the CD40.CoV2 vaccine or an equimolar combination of 15-mer overlapping peptide pools covering the full-length sequence of vaccine antigens (vS1 + vS2 + vRBD + vN2) referred to as vOLPmix. PBMCs (1 x 10^6^) were incubated in 300 µl RPMI supplemented with 10% human serum AB (SAB) for 24 h at 37°C in 5% CO_2_. T-cell activation was assessed by detection of the extracellular activation markers CD69 and CD137, in addition to a viability marker and CD3, CD4, and CD8 to determine the T-cell lineage. For the EpiMax technology, PBMCs were stimulated *in vitro* with 1 nM CD40.CoV2 vaccine on D0 and restimulated on D8 with 1 µg/ml of various vOLPs (vS1, vRBD, vS2, or vN2). Cell functionality was assessed by intracellular cytokine staining (ICS), with Boolean gating. The flow cytometry panel included a viability marker, CD3, CD4, and CD8 to determine the T-cell lineage, and IFN-γ, TNF, and IL-2 antibodies. Distributions were plotted using SPICE version 5.22, downloaded from http://exon.niaid.nih.gov/spice (Roederer et al., 2011). T-cell proliferation was evaluated using the CellTrace™ CFSE Cell Proliferation Kit (Invitrogen) as previously described (Tario et al., 2011). PBMCs were stimulated *in vitro* with 1 nM CD40.CoV2 vaccine or an equimolar amount of vOLPmix for seven days without IL-2. The medium (RPMI 10% SAB) was changed 48 h after stimulation.

#### Cytotoxicity assay

PBMCs from convalescent COVID-19 patients were stimulated with 1 nM CD40.CoV2 vaccine (effector cells) or 2.5 µg/mL phytohemagglutinin-L (PHA-L) (ThermoFischer Scientific) (target-Blast cells) in RPMI 10% SAB medium replenished every 2 to 3 days with fresh medium supplemented with IL-2 (100 U/mL) (Myltenyi Biotec). After seven days, effector-CD8^+^ T cells were isolated using a CD8^+^ T-cell isolation kit from Myltenyi Biotec following the manufacturers’ instructions and target-Blast cells were pulsed for 1 h at 37°C in 5% CO_2_, either with 0.4% DMSO (control) or 2 µg/mL vOLP. After two washes, cells were labeled using various combinations of carboxyfluorescein succinimidyl ester (CFSE) (0.1 µM), Cell Trace Violet (CTV) (0.5 µM), and/or Cell Trace Far Red (CTFR) (0.02 µM) (Thermofisher Scientific) for 15 min at 37°C. Target-Blast specific populations were then mixed at a 1:1 ratio and co-cultured with effector-CD8^+^ T-cells at various ratios in triplicate. To measure basal apoptosis, three wells were seeded with target-Blast cells alone. After a 24 h-incubation, cells were stained with LIVE/DEAD Fixable Near-IR stain (Thermofisher Scientific) and analyzed using an LSR II-3 laser flow cytometer (405, 488, and 640 nm) (Becton Dickinson). The percentage of specific cytotoxicity among live cells was calculated as follows: specific lysis (%) = 100*[(average [count vOLP pulsed/count DMSO pulsed] target-Blast cells alone - [count vOLP pulsed/count.DMSO pulsed]) target-Blast cells + CD8^+^ effector T cells / (average [count vOLP pulsed/count DMSO pulsed] target-Blast cells alone)].

#### Statistical analysis

Graphpad Prism software version 8 was used for nonparametric statistics and plots, as described in the figure legends. Heatmaps were generated using the heatmap function from package NMF in R software, version 4.0.0. R: A language and environment for statistical computing. R Foundation for Statistical Computing, Vienna, Austria. URL: https://www.R-project.org. Statistical differences in the expression of standardized biomarkers were determined using the nonparametric Wilcoxon test, adjusting for multiple testing using the Benjamini & Hochberg correction.

## Supplemental information titles and legends

- **French Cohort Study Group**
- **Table S1. CD40.CoV2 vaccine region characteristics**
- **Table S4. Vaccine region homology between β and α coronaviruses**
- **Table S5. Vaccine CD4 and CD8 T-cell epitopes with 100% homology with 38 sarbecoviruses**
- **Figure S1. Clinical symptom scores for CD40.CoV2 vaccinated and mock-vaccinated hCD40/K18-hACE2 mice from days 0 to 12 post-inoculation with SARS-CoV-2.** The clinical score, ranging from 0 to 4, was monitored daily until day 12 post-inoculation (dpi) and was calculated according to the presence of four symptoms: eye closure, ruffled fur, hunched posture, and labored breathing.
- **Figure S2. IgG responses against MERS and common cold coronaviruses in CD40.CoV2 vaccinated and mock-vaccinated hCD40/K18-hACE2 mice.** Level of IgG antibodies (AU) binding to spike proteins from MERS and common cold coronaviruses in vaccinated and mock-vaccinated animals before vaccination (baseline, -2 days post-vaccination [dpv], n = 9-12 animals per group), after completion of the vaccination schedule (28 dpv, n = 9-12 animals per group), and 40 dpv (i.e., day 12 pi time point, n = 3-5 animals per group). The medians [Min-Max] are shown. Grey dashed lines represent the prime and boost vaccination. The red dashed line represents SARS-CoV2 inoculation.
- **Figure S3. Induction of cross-reactive and neutralizing IgG antibody responses by the CD40.CoV2 vaccine in a second replicated animal experiment.** Level of IgG antibodies (AU) binding to Wuhan and VOC SARS-CoV-2 RBD proteins (A) and Spike protein from MERS and common cold coronaviruses (B) in mock-vaccinated and vaccinated animals before vaccination (baseline, -2 days post-vaccination [dpv], n = 10-12 animals per group), after completion of the vaccination schedule (28 dpv, n = 10-12 animals per group), and 40 dpv (i.e., day 12 pi time point, n = 3 animals per group). (C) Levels of IgG antibodies (AU) binding to SARS-CoV-2 (solid lines) and SARS-CoV-1 (dashed lines) S proteins in mock-vaccinated (grey) and vaccinated (blue) animals at -2, 28, and 40 dpv. Medians [Min-Max] are shown. Grey dashed lines represent the prime and boost vaccination. The red dashed line represents SARS-CoV-2 inoculation. Neutralizing activity of RBD antibodies (Unit/mL) (D) and S antibodies (Unit/mL) (E) in mock-vaccinated (grey) and vaccinated (blue) animals post-vaccination (open circle) and post-infection (solid circle). Because mock-vaccinated animals have no SARS-CoV-2 antibody responses before infection, neutralization activity was evaluated in sera from only three mock-vaccinated animals for which samples before and after infection were available. Medians ± IQRs are shown. Thirty plasma samples from unvaccinated mice were used to determine the positivity threshold, defined as the whole units/mL value immediately above the concentration of the highest sample for RBD (i.e., 8 units/mL) and Spike (i.e., 4 units/mL) proteins.
- **Figure S4. Production of TNF and IL-2 by SARS-CoV-2-specific CD4^+^ and CD8^+^ T cells from convalescent COVID-19 patients after *in-vitro* stimulation with the CD40.CoV2 vaccine.** Frequency of TNF^+^ and IL-2^+^ SARS-CoV2 specific CD4^+^ T cells (A) or specific CD8^+^ T cells (B) from convalescent COVID-19 patients (n = 14) stimulated or not with the CD40.CoV2 vaccine (1 nM) on D0 and re-stimulated with various vOLPs (vRBD, vS1, vS2, or vN2) or cont.OLPs (Gpz or N1-N2, grey) on D8 (1 µg/ml). Median values ± IQRs are shown. Friedman and Dunn’s multiple comparison tests were used for statistical analysis (*P < 0.05, **P < 0.01, ***P < 0.001, ****P < 0.0001, ns: not significant).
- **Figure S5. Simple linear regression between SARS-CoV-1- and SARS-CoV-2- specific CD4^+^ and CD8^+^ T-cell responses.** Correlation between the frequency of total cytokines (IFN-γ ± IL-2 ± TNF) secreted by specific CD4^+^ (A) or CD8^+^ (B) T cells after stimulation with the CD40.CoV2 vaccine (1 nM) on D0 and re-stimulation with OLPs representing the sequence of S1, vRBD, and vN2 from either the SARS CoV-1 or SARS CoV-2 virus (1 µg/mL). Pearson’s test was used for statistical significance. Simple linear regression is represented as a solid line and the 95% confidence interval as dashed lines.

## Supplemental items (provided as Excel files)

- **Table S2. Conservation between SARS-CoV-2 CD8^+^ T-cell epitopes from vaccine sequences with 38 sarbecoviruses (related to** **Figure 1**). Mutations and deletions from the original Wuhan SARS-CoV-2 sequence are highlighted in red. Mutated CD8^+^ T-cell epitopes no longer able to bind to MHC-restriction molecule for SARS-CoV-2 VOCs or SARS-CoV-1 are colored in grey. The color gradient from yellow to green represents the increasing percentage of homology between vaccine antigen sequences and those of SARS-CoV1 (column J) and sarbecoviruses (column K).
- **Table S3. Conservation between SARS-CoV-2 CD4^+^ T-cell epitopes from vaccine sequences with 38 sarbecoviruses (related to** **Figure 1**). Mutations and deletions from the original Wuhan SARS-CoV-2 sequence are highlighted in red. The color gradient from yellow to green represents the increasing percentage of homology between vaccine antigen sequences and those of SARS-CoV1 (column J) and sarbecoviruses (column K).

## References

Ahmed, S.F., Quadeer, A.A., and McKay, M.R. (2020). Preliminary Identification of Potential Vaccine Targets for the COVID-19 Coronavirus (SARS-CoV-2) Based on SARS-CoV Immunological Studies. Viruses 12, 254.

Anderson, E.J., Rouphael, N.G., Widge, A.T., Jackson, L.A., Roberts, P.C., Makhene, M., Chappell, J.D., Denison, M.R., Stevens, L.J., Pruijssers, A.J., et al. (2020). Safety and Immunogenicity of SARS-CoV-2 mRNA-1273 Vaccine in Older Adults. N Engl J Med 383, 2427–2438.

Andreatta, M., and Nielsen, M. (2016). Gapped sequence alignment using artificial neural networks: application to the MHC class I system. Bioinformatics 32, 511–517.

Angyal, A., Longet, S., Moore, S., Payne, R.P., Harding, A., Tipton, T., Rongkard, P., Ali, M., Hering, L.M., Meardon, N., et al. (2021). T-Cell and Antibody Responses to First BNT162b2 Vaccine Dose in Previously SARS-CoV-2-Infected and Infection-Naive UK Healthcare Workers: A Multicentre, Prospective, Observational Cohort Study. SSRN Journal.

Bange, E.M., Han, N.A., Wileyto, P., Kim, J.Y., Gouma, S., Robinson, J., Greenplate, A.R., Hwee, M.A., Porterfield, F., Owoyemi, O., et al. (2021). CD8+ T cells contribute to survival in patients with COVID-19 and hematologic cancer. Nat Med 27, 1280–1289.

Barda, N., Dagan, N., Cohen, C., Hernán, M.A., Lipsitch, M., Kohane, I.S., Reis, B.Y., and Balicer, R.D. (2021). Effectiveness of a third dose of the BNT162b2 mRNA COVID-19 vaccine for preventing severe outcomes in Israel: an observational study. The Lancet 398, 2093–2100.

Baruah, V., and Bose, S. (2020). Immunoinformatics-aided identification of T cell and B cell epitopes in the surface glycoprotein of 2019-nCoV. J Med Virol 92, 495–500.

Boni, M.F., Lemey, P., Jiang, X., Lam, T.T.-Y., Perry, B.W., Castoe, T.A., Rambaut, A., and Robertson, D.L. (2020). Evolutionary origins of the SARS-CoV-2 sarbecovirus lineage responsible for the COVID-19 pandemic. Nat Microbiol 5, 1408–1417.

Bouteau, A., Kervevan, J., Su, Q., Zurawski, S.M., Contreras, V., Dereuddre-Bosquet, N., Le Grand, R., Zurawski, G., Cardinaud, S., Levy, Y., et al. (2019). DC Subsets Regulate Humoral Immune Responses by Supporting the Differentiation of Distinct Tfh Cells. Front. Immunol. 10, 1134.

Cameroni, E., Bowen, J.E., Rosen, L.E., Saliba, C., Zepeda, S.K., and Culap, K. 1 Broadly neutralizing antibodies overcome SARS-CoV-2 Omicron 2 antigenic shift. 53.

Ceglia, V., Zurawski, S., Montes, M., Bouteau, A., Wang, Z., Ellis, J., Igyártó, B.Z., Lévy, Y., and Zurawski, G. (2021). Anti-CD40 Antibodies Fused to CD40 Ligand Have Superagonist Properties. J.I. 207, 2060–2076.

Cele, S., Jackson, L., Khoury, D.S., Khan, K., Moyo-Gwete, T., Tegally, H., San, J.E., Cromer, D., Scheepers, C., Amoako, D., et al. (2021). Omicron extensively but incompletely escapes Pfizer BNT162b2 neutralization. Nature d41586-021-03824–03825.

Chatterjee, B., Smed-Sörensen, A., Cohn, L., Chalouni, C., Vandlen, R., Lee, B.-C., Widger, J., Keler, T., Delamarre, L., and Mellman, I. (2012). Internalization and endosomal degradation of receptor-bound antigens regulate the efficiency of cross presentation by human dendritic cells. Blood 120, 2011–2020.

Cheng, L., Zhang, Z., Li, G., Li, F., Wang, L., Zhang, L., Zurawski, S.M., Zurawski, G., Levy, Y., and Su, L. (2017). Human innate responses and adjuvant activity of TLR ligands in vivo in mice reconstituted with a human immune system. Vaccine 35, 6143–6153.

Cheng, L., Wang, Q., Li, G., Banga, R., Ma, J., Yu, H., Yasui, F., Zhang, Z., Pantaleo, G., Perreau, M., et al. (2018). TLR3 agonist and CD40-targeting vaccination induces immune responses and reduces HIV-1 reservoirs. Journal of Clinical Investigation 128, 4387–4396.

Cherian, S., Potdar, V., Jadhav, S., Yadav, P., Gupta, N., Das, M., Rakshit, P., Singh, S., Abraham, P., Panda, S., et al. (2021). Convergent evolution of SARS-CoV-2 spike mutations, L452R, E484Q and P681R, in the second wave of COVID-19 in Maharashtra, India (Molecular Biology).

Chujo, D., Foucat, E., Nguyen, T.-S., Chaussabel, D., Banchereau, J., and Ueno, H. (2013). ZnT8-Specific CD4+ T Cells Display Distinct Cytokine Expression Profiles between Type 1 Diabetes Patients and Healthy Adults. PLoS ONE 8, e55595.

Cohen, J. (2021). The dream vaccine. Science 372, 227–231.

Dahlke, C., Heidepriem, J., Kobbe, R., Santer, R., Koch, T., Fathi, A., Ly, M.L., Schmiedel, S., Seeberger, P.H., ID-UKE COVID-19 study group, et al. (2020). Distinct early IgA profile may determine severity of COVID-19 symptoms: an immunological case series (Infectious Diseases (except HIV/AIDS)).

Dangi, T., Palacio, N., Sanchez, S., Park, M., Class, J., Visvabharathy, L., Ciucci, T., Koralnik, I.J., Richner, J.M., and Penaloza-MacMaster, P. (2021). Cross-protective immunity following coronavirus vaccination and coronavirus infection. Journal of Clinical Investigation 131, e151969.

Davies, N.G., Abbott, S., Barnard, R.C., Jarvis, C.I., Kucharski, A.J., Munday, J.D., Pearson, C.A.B., Russell, T.W., Tully, D.C., Washburne, A.D., et al. (2021). Estimated transmissibility and impact of SARS-CoV-2 lineage B.1.1.7 in England. Science 372, eabg3055.

Fast, E., Altman, R.B., and Chen, B. (2020). Potential T-cell and B-cell Epitopes of 2019-nCoV (Microbiology).

Ferretti, A.P., Kula, T., Wang, Y., Nguyen, D.M.V., Weinheimer, A., Dunlap, G.S., Xu, Q., Nabilsi, N., Perullo, C.R., Cristofaro, A.W., et al. (2020). Unbiased Screens Show CD8+ T Cells of COVID-19 Patients Recognize Shared Epitopes in SARS-CoV-2 that Largely Reside outside the Spike Protein. Immunity 53, 1095–1107.e3.

Flamar, A.-L., Xue, Y., Zurawski, S.M., Montes, M., King, B., Sloan, L., Oh, S., Banchereau, J., Levy, Y., and Zurawski, G. (2013). Targeting concatenated HIV antigens to human CD40 expands a broad repertoire of multifunctional CD4+ and CD8+ T cells. AIDS 27, 2041–2051.

Flamar, A.-L., Bonnabau, H., Zurawski, S., Lacabaratz, C., Montes, M., Richert, L., Wiedemann, A., Galmin, L., Weiss, D., Cristillo, A., et al. (2018). HIV-1 T cell epitopes targeted to Rhesus macaque CD40 and DCIR: A comparative study of prototype dendritic cell targeting therapeutic vaccine candidates. PLoS ONE 13, e0207794.

Frutos, R., Serra-Cobo, J., Pinault, L., Lopez Roig, M., and Devaux, C.A. (2021). Emergence of Bat-Related Betacoronaviruses: Hazard and Risks. Front. Microbiol. 12, 591535.

Funk, T., Pharris, A., Spiteri, G., Bundle, N., Melidou, A., Carr, M., Gonzalez, G., Garcia-Leon, A., Crispie, F., O’Connor, L., et al. (2021). Characteristics of SARS-CoV-2 variants of concern B.1.1.7, B.1.351 or P.1: data from seven EU/EEA countries, weeks 38/2020 to 10/2021. Eurosurveillance 26.

Gao, A., Chen, Z., Amitai, A., Doelger, J., Mallajosyula, V., Sundquist, E., Pereyra Segal, F., Carrington, M., Davis, M.M., Streeck, H., et al. (2021). Learning from HIV-1 to predict the immunogenicity of T cell epitopes in SARS-CoV-2. IScience 24, 102311.

Garcia-Beltran, W.F., Lam, E.C., St. Denis, K., Nitido, A.D., Garcia, Z.H., Hauser, B.M., Feldman, J., Pavlovic, M.N., Gregory, D.J., Poznansky, M.C., et al. (2021). Multiple SARS-CoV-2 variants escape neutralization by vaccine-induced humoral immunity. Cell 184, 2372–2383.e9.

Godot, V., Tcherakian, C., Gil, L., Cervera-Marzal, I., Li, G., Cheng, L., Ortonne, N., Lelièvre, J.-D., Pantaleo, G., Fenwick, C., et al. (2020). TLR-9 agonist and CD40-targeting vaccination induces HIV-1 envelope-specific B cells with a diversified immunoglobulin repertoire in humanized mice. PLoS Pathog 16, e1009025.

Graham, J.P., Authie, P., Yu, C.I., Zurawski, S.M., Li, X.-H., Marches, F., Flamar, A.-L., Acharya, A., Banchereau, J., and Palucka, A.K. (2016). Targeting dendritic cells in humanized mice receiving adoptive T cells via monoclonal antibodies fused to Flu epitopes. Vaccine 34, 4857–4865.

Grifoni, A., Weiskopf, D., Ramirez, S.I., Mateus, J., Dan, J.M., Moderbacher, C.R., Rawlings, S.A., Sutherland, A., Premkumar, L., Jadi, R.S., et al. (2020). Targets of T Cell Responses to SARS-CoV-2 Coronavirus in Humans with COVID-19 Disease and Unexposed Individuals. Cell 181, 1489–1501.e15.

Grifoni, A., Sidney, J., Vita, R., Peters, B., Crotty, S., Weiskopf, D., and Sette, A. (2021). SARS-CoV-2 human T cell epitopes: Adaptive immune response against COVID-19. Cell Host & Microbe 29, 1076–1092.

Guccione, E. Differential effects of the second SARS-CoV-2 mRNA vaccine dose on T cell immunity in naïve and COVID-19 recovered individuals. 9.

Hadjadj, J., Planas, D., Ouedrani, A., Buffier, S., Delage, L., Nguyen, Y., Bruel, T., Stolzenberg, M.-C., Staropoli, I., Ermak, N., et al. (2021). Immunogenicity of BNT162b2 vaccine Against the Alpha and Delta Variants in Immunocompromised Patients (Infectious Diseases (except HIV/AIDS)).

Hoffmann, M., Kleine-Weber, H., Schroeder, S., Krüger, N., Herrler, T., Erichsen, S., Schiergens, T.S., Herrler, G., Wu, N.-H., Nitsche, A., et al. (2020). SARS-CoV-2 Cell Entry Depends on ACE2 and TMPRSS2 and Is Blocked by a Clinically Proven Protease Inhibitor. Cell 181, 271–280.e8.

Hoffmann, M., Arora, P., Groß, R., Seidel, A., Hörnich, B.F., Hahn, A.S., Krüger, N., Graichen, L., Hofmann-Winkler, H., Kempf, A., et al. (2021). SARS-CoV-2 variants B.1.351 and P.1 escape from neutralizing antibodies. Cell 184, 2384–2393.e12.

Hu, C., Shen, M., Han, X., Chen, Q., Li, L., Chen, S., Zhang, J., Gao, F., Wang, W., Wang, Y., et al. (2021). Identification of cross-reactive CD8+ T cell receptors with high functional avidity to a SARS-CoV-2 immunodominant epitope and its natural mutant variants. Genes & Diseases S2352304221000842.

Hyun-Jung Lee, C., and Koohy, H. (2020). In silico identification of vaccine targets for 2019-nCoV. F1000Res 9, 145.

Israel, A., Merzon, E., Schäffer, A.A., Shenhar, Y., Green, I., Golan-Cohen, A., Ruppin, E., Magen, E., and Vinker, S. (2021). Elapsed time since BNT162b2 vaccine and risk of SARS-CoV-2 infection in a large cohort (Infectious Diseases (except HIV/AIDS)).

Jensen, K.K., Andreatta, M., Marcatili, P., Buus, S., Greenbaum, J.A., Yan, Z., Sette, A., Peters, B., and Nielsen, M. (2018). Improved methods for predicting peptide binding affinity to MHC class II molecules. Immunology 154, 394–406.

Jespersen, M.C., Peters, B., Nielsen, M., and Marcatili, P. (2017). BepiPred-2.0: improving sequence-based B-cell epitope prediction using conformational epitopes. Nucleic Acids Research 45, W24–W29.

Kalimuddin, S., Qui, M., Eong, E., Bertoletti, A., and Low, J.G. Early T cell and binding antibody responses are associated with COVID-19 RNA vaccine efficacy onset. OPEN ACCESS 13.

Kared, H., Redd, A.D., Bloch, E.M., Bonny, T.S., Sumatoh, H., Kairi, F., Carbajo, D., Abel, B., Newell, E.W., Bettinotti, M.P., et al. (2021). SARS-CoV-2–specific CD8+ T cell responses in convalescent COVID-19 individuals. Journal of Clinical Investigation 131, e145476.

Karim, S.S.A., and Karim, Q.A. (2021). Omicron SARS-CoV-2 variant: a new chapter in the COVID-19 pandemic. The Lancet 398, 2126–2128.

Kumar, S., Chandele, A., and Sharma, A. (2021). Current status of therapeutic monoclonal antibodies against SARS-CoV-2. PLoS Pathog 17, e1009885.

Kustin, T., Harel, N., Finkel, U., Perchik, S., Harari, S., Tahor, M., Caspi, I., Levy, R., Leshchinsky, M., Ken Dror, S., et al. (2021). Evidence for increased breakthrough rates of SARS-CoV-2 variants of concern in BNT162b2-mRNA-vaccinated individuals. Nat Med 27, 1379–1384.

Lam, T.T.-Y., Jia, N., Zhang, Y.-W., Shum, M.H.-H., Jiang, J.-F., Zhu, H.-C., Tong, Y.-G., Shi, Y.-X., Ni, X.-B., Liao, Y.-S., et al. (2020). Identifying SARS-CoV-2-related coronaviruses in Malayan pangolins. Nature 583, 282–285.

Le Bert, N., Tan, A.T., Kunasegaran, K., Tham, C.Y.L., Hafezi, M., Chia, A., Chng, M.H.Y., Lin, M., Tan, N., Linster, M., et al. (2020). SARS-CoV-2-specific T cell immunity in cases of COVID-19 and SARS, and uninfected controls. Nature 584, 457–462.

Lederer, K., Castaño, D., Gómez Atria, D., Oguin, T.H., Wang, S., Manzoni, T.B., Muramatsu, H., Hogan, M.J., Amanat, F., Cherubin, P., et al. (2020). SARS-CoV-2 mRNA Vaccines Foster Potent Antigen-Specific Germinal Center Responses Associated with Neutralizing Antibody Generation. Immunity 53, 1281–1295.e5.

Li, T., Xie, J., He, Y., Fan, H., Baril, L., Qiu, Z., Han, Y., Xu, W., Zhang, W., You, H., et al. (2006). Long-Term Persistence of Robust Antibody and Cytotoxic T Cell Responses in Recovered Patients Infected with SARS Coronavirus. PLoS ONE 1, e24.

Lineburg, K.E., Grant, E.J., Swaminathan, S., Chatzileontiadou, D.S.M., Szeto, C., Sloane, H., Panikkar, A., Raju, J., Crooks, P., Rehan, S., et al. (2021). CD8+ T cells specific for an immunodominant SARS-CoV-2 nucleocapsid epitope cross-react with selective seasonal coronaviruses. Immunity 54, 1055–1065.e5.

Lu, M., Uchil, P.D., Li, W., Zheng, D., Terry, D.S., Gorman, J., Shi, W., Zhang, B., Zhou, T., Ding, S., et al. (2020). Real-Time Conformational Dynamics of SARS-CoV-2 Spikes on Virus Particles. Cell Host & Microbe 28, 880–891.e8.

Madhi, S.A., Baillie, V., Cutland, C.L., Voysey, M., Koen, A.L., Fairlie, L., Padayachee, S.D., Dheda, K., Barnabas, S.L., Bhorat, Q.E., et al. (2021). Efficacy of the ChAdOx1 nCoV-19 Covid-19 Vaccine against the B.1.351 Variant. N Engl J Med 384, 1885–1898.

Marlin, R., Godot, V., Cardinaud, S., Galhaut, M., Coleon, S., Zurawski, S., Dereuddre-Bosquet, N., Cavarelli, M., Gallouët, A.-S., Maisonnasse, P., et al. (2021). Targeting SARS-CoV-2 receptor-binding domain to cells expressing CD40 improves protection to infection in convalescent macaques. Nat Commun 12, 5215.

Mateus, J., Grifoni, A., Tarke, A., Sidney, J., Ramirez, S.I., Dan, J.M., Burger, Z.C., Rawlings, S.A., Smith, D.M., Phillips, E., et al. (2020). Selective and cross-reactive SARS-CoV-2 T cell epitopes in unexposed humans. Science 370, 89–94.

Mazzoni, A., Di Lauria, N., Maggi, L., Salvati, L., Vanni, A., Capone, M., Lamacchia, G., Mantengoli, E., Spinicci, M., Zammarchi, L., et al. (2021). First-dose mRNA vaccination is sufficient to reactivate immunological memory to SARS-CoV-2 in subjects who have recovered from COVID-19. Journal of Clinical Investigation 131, e149150.

McMahan, K., Yu, J., Mercado, N.B., Loos, C., Tostanoski, L.H., Chandrashekar, A., Liu, J., Peter, L., Atyeo, C., Zhu, A., et al. (2021). Correlates of protection against SARS-CoV-2 in rhesus macaques. Nature 590, 630–634.

Menachery, V.D., Yount, B.L., Debbink, K., Agnihothram, S., Gralinski, L.E., Plante, J.A., Graham, R.L., Scobey, T., Ge, X.-Y., Donaldson, E.F., et al. (2015). A SARS-like cluster of circulating bat coronaviruses shows potential for human emergence. Nat Med 21, 1508–1513.

Menachery, V.D., Yount, B.L., Sims, A.C., Debbink, K., Agnihothram, S.S., Gralinski, L.E., Graham, R.L., Scobey, T., Plante, J.A., Royal, S.R., et al. (2016). SARS-like WIV1-CoV poised for human emergence. Proc Natl Acad Sci USA 113, 3048–3053.

Motozono, C., Toyoda, M., Zahradnik, J., Saito, A., Nasser, H., Tan, T.S., Ngare, I., Kimura, I., Uriu, K., Kosugi, Y., et al. (2021). SARS-CoV-2 spike L452R variant evades cellular immunity and increases infectivity. Cell Host & Microbe 29, 1124–1136.e11.

Nathan, A., Rossin, E.J., Kaseke, C., Park, R.J., Khatri, A., Koundakjian, D., Urbach, J.M., Singh, N.K., Bashirova, A., Tano-Menka, R., et al. (2021). Structure-guided T cell vaccine design for SARS-CoV-2 variants and sarbecoviruses. Cell 184, 4401–4413.e10.

Nelde, A., Bilich, T., Heitmann, J.S., Maringer, Y., Salih, H.R., Roerden, M., Lübke, M., Bauer, J., Rieth, J., Wacker, M., et al. (2021). SARS-CoV-2-derived peptides define heterologous and COVID-19-induced T cell recognition. Nat Immunol 22, 74–85.

Painter, M.M., Mathew, D., Goel, R.R., Apostolidis, S.A., Pattekar, A., Kuthuru, O., Baxter, A.E., Herati, R.S., Oldridge, D.A., Gouma, S., et al. (2021). Rapid induction of antigen-specific CD4+ T cells is associated with coordinated humoral and cellular immune responses to SARS-CoV-2 mRNA vaccination. Immunity S1074761321003083.

Peng, Y., Mentzer, A.J., Liu, G., Yao, X., Yin, Z., Dong, D., Dejnirattisai, W., Rostron, T., Supasa, P., Liu, C., et al. (2020). Broad and strong memory CD4+ and CD8+ T cells induced by SARS-CoV-2 in UK convalescent individuals following COVID-19. Nat Immunol 21, 1336–1345.

Planas, D., Saunders, N., Maes, P., Guivel-Benhassine, F., Planchais, C., Buchrieser, J., Bolland, W.-H., Porrot, F., Staropoli, I., Lemoine, F., et al. (2021). Considerable escape of SARS-CoV-2 Omicron to antibody neutralization. Nature d41586-021-03827–2.

Poran, A., Harjanto, D., Malloy, M., Arieta, C.M., Rothenberg, D.A., Lenkala, D., van Buuren, M.M., Addona, T.A., Rooney, M.S., Srinivasan, L., et al. (2020). Sequence-based prediction of SARS-CoV-2 vaccine targets using a mass spectrometry-based bioinformatics predictor identifies immunogenic T cell epitopes. Genome Med 12, 70.

Prachar, M., Justesen, S., Steen-Jensen, D.B., Thorgrimsen, S., Jurgons, E., Winther, O., and Bagger, F.O. (2020). Identification and validation of 174 COVID-19 vaccine candidate epitopes reveals low performance of common epitope prediction tools. Sci Rep 10, 20465.

Prendecki, M., Clarke, C., Edwards, H., McIntyre, S., Mortimer, P., Gleeson, S., Martin, P., Thomson, T., Randell, P., Shah, A., et al. (2021). Humoral and T-cell responses to SARS-CoV-2 vaccination in patients receiving immunosuppression. Ann Rheum Dis annrheumdis-2021-220626.

Puranik, A., Lenehan, P.J., Silvert, E., Niesen, M.J.M., Corchado-Garcia, J., O’Horo, J.C., Virk, A., Swift, M.D., Halamka, J., Badley, A.D., et al. Comparison of two highly-effective mRNA vaccines for COVID-19 during periods of Alpha and Delta variant prevalence. 29.

Redd, A.D., Nardin, A., Kared, H., Bloch, E.M., Abel, B., Pekosz, A., Laeyendecker, O., Fehlings, M., Quinn, T.C., and Tobian, A.A. (2021). Minimal cross-over between mutations associated with Omicron variant of SARS-CoV-2 and CD8+ T cell epitopes identified in COVID-19 convalescent individuals (Immunology).

Rha, M.-S., Jeong, H.W., Ko, J.-H., Choi, S.J., Seo, I.-H., Lee, J.S., Sa, M., Kim, A.R., Joo, E.-J., Ahn, J.Y., et al. (2021). PD-1-Expressing SARS-CoV-2-Specific CD8+ T Cells Are Not Exhausted, but Functional in Patients with COVID-19. Immunity 54, 44–52.e3.

Roederer, M., Nozzi, J.L., and Nason, M.C. (2011). SPICE: Exploration and analysis of post-cytometric complex multivariate datasets. Cytometry 79A, 167–174.

Rössler, A., Riepler, L., Bante, D., Laer, D. von, and Kimpel, J. (2021). SARS-CoV-2 B.1.1.529 variant (Omicron) evades neutralization by sera from vaccinated and convalescent individuals (Infectious Diseases (except HIV/AIDS)).

Sahin, U., Muik, A., Derhovanessian, E., Vogler, I., Kranz, L.M., Vormehr, M., Baum, A., Pascal, K., Quandt, J., Maurus, D., et al. (2020). COVID-19 vaccine BNT162b1 elicits human antibody and TH1 T cell responses. Nature 586, 594–599.

Schulien, I., Kemming, J., Oberhardt, V., Wild, K., Seidel, L.M., Killmer, S., Sagar, Daul, F., Salvat Lago, M., Decker, A., et al. (2021). Characterization of pre-existing and induced SARS-CoV-2-specific CD8+ T cells. Nat Med 27, 78–85.

Sette, A., and Crotty, S. (2021). Adaptive immunity to SARS-CoV-2 and COVID-19. Cell 184, 861–880.

Shomuradova, A.S., Vagida, M.S., Sheetikov, S.A., Zornikova, K.V., Kiryukhin, D., Titov, A., Peshkova, I.O., Khmelevskaya, A., Dianov, D.V., Malasheva, M., et al. (2020). SARS-CoV-2 Epitopes Are Recognized by a Public and Diverse Repertoire of Human T Cell Receptors. Immunity 53, 1245–1257.e5.

Stamatatos, L., Czartoski, J., Wan, Y.-H., Homad, L.J., Rubin, V., Glantz, H., Neradilek, M., Seydoux, E., Jennewein, M.F., MacCamy, A.J., et al. (2021). mRNA vaccination boosts cross-variant neutralizing antibodies elicited by SARS-CoV-2 infection. Science 372, 1413–1418.

Tan, A.T., Linster, M., Tan, C.W., Le Bert, N., Chia, W.N., Kunasegaran, K., Zhuang, Y., Tham, C.Y.L., Chia, A., Smith, G.J.D., et al. (2021). Early induction of functional SARS-CoV-2-specific T cells associates with rapid viral clearance and mild disease in COVID-19 patients. Cell Reports 34, 108728.

Tario, J.D., Muirhead, K.A., Pan, D., Munson, M.E., and Wallace, P.K. (2011). Tracking Immune Cell Proliferation and Cytotoxic Potential Using Flow Cytometry. In Flow Cytometry Protocols, T.S. Hawley, and R.G. Hawley, eds. (Totowa, NJ: Humana Press), pp. 119–164.

Tarke, A., Sidney, J., Methot, N., Yu, E.D., Zhang, Y., Dan, J.M., Goodwin, B., Rubiro, P., Sutherland, A., Wang, E., et al. (2021a). Impact of SARS-CoV-2 variants on the total CD4+ and CD8+ T cell reactivity in infected or vaccinated individuals. Cell Reports Medicine 2, 100355.

Tarke, A., Sidney, J., Kidd, C.K., Dan, J.M., Ramirez, S.I., Yu, E.D., Mateus, J., da Silva Antunes, R., Moore, E., Rubiro, P., et al. (2021b). Comprehensive analysis of T cell immunodominance and immunoprevalence of SARS-CoV-2 epitopes in COVID-19 cases. Cell Reports Medicine 2, 100204.

Tegally, H., Wilkinson, E., Lessells, R.J., Giandhari, J., Pillay, S., Msomi, N., Mlisana, K., Bhiman, J.N., von Gottberg, A., Walaza, S., et al. (2021). Sixteen novel lineages of SARS-CoV-2 in South Africa. Nat Med 27, 440–446.

Voloch, C.M., Gerber, A.L., Leitão, I. de C., Galliez, R.M., and Faffe, D.S. (2021). Genomic Characterization of a Novel SARS-CoV-2 Lineage from Rio de Janeiro, Brazil. Journal of Virology 95, 5.

Wacharapluesadee, S., Tan, C.W., Maneeorn, P., Duengkae, P., Zhu, F., Joyjinda, Y., Kaewpom, T., Chia, W.N., Ampoot, W., Lim, B.L., et al. (2021). Evidence for SARS-CoV-2 related coronaviruses circulating in bats and pangolins in Southeast Asia. Nat Commun 12, 972.

Wang, P., Nair, M.S., Liu, L., Iketani, S., Luo, Y., Guo, Y., Wang, M., Yu, J., Zhang, B., Kwong, P.D., et al. (2021). Antibody resistance of SARS-CoV-2 variants B.1.351 and B.1.1.7. Nature 593, 130–135.

Ward, J.H. (1963). Hierarchical Grouping to Optimize an Objective Function. Journal of the American Statistical Association 58, 236–244.

Weiskopf, D., Schmitz, K.S., Raadsen, M.P., Grifoni, A., Okba, N.M.A., Endeman, H., van den Akker, J.P.C., Molenkamp, R., Koopmans, M.P.G., van Gorp, E.C.M., et al. (2020). Phenotype and kinetics of SARS-CoV-2–specific T cells in COVID-19 patients with acute respiratory distress syndrome. Sci. Immunol. 5, eabd2071.

Wibmer, C.K., Ayres, F., Hermanus, T., Madzivhandila, M., Kgagudi, P., Oosthuysen, B., Lambson, B.E., de Oliveira, T., Vermeulen, M., van der Berg, K., et al. (2021). SARS-CoV-2 501Y.V2 escapes neutralization by South African COVID-19 donor plasma. Nat Med 27, 622–625.

Wu, F., Zhao, S., Yu, B., Chen, Y.-M., Wang, W., Song, Z.-G., Hu, Y., Tao, Z.-W., Tian, J.-H., Pei, Y.-Y., et al. (2020). A new coronavirus associated with human respiratory disease in China. Nature 579, 265–269.

Yazdanpanah, Y., French COVID cohort investigators and study group, Diallo, A., Paul, C., Mercier, N., Le Mestre, S., Petrov-Sanchez, V., Andrejak, C., Chirouze, C., Malvy, D., et al. (2021). Impact on disease mortality of clinical, biological, and virological characteristics at hospital admission and overtime in COVID-19 patients. J Med Virol 93, 2149–2159.

Yin, W., Gorvel, L., Zurawski, S., Li, D., Ni, L., Duluc, D., Upchurch, K., Kim, J., Gu, C., Ouedraogo, R., et al. (2016). Functional Specialty of CD40 and Dendritic Cell Surface Lectins for Exogenous Antigen Presentation to CD8+ and CD4+ T Cells. EBioMedicine 5, 46–58.

Zhou, P., Yang, X.-L., Wang, X.-G., Hu, B., Zhang, L., Zhang, W., Si, H.-R., Zhu, Y., Li, B., Huang, C.-L., et al. (2020). A pneumonia outbreak associated with a new coronavirus of probable bat origin. Nature 579, 270–273.

Zurawski, G., Shen, X., Zurawski, S., Tomaras, G.D., Montefiori, D.C., Roederer, M., Ferrari, G., Lacabaratz, C., Klucar, P., Wang, Z., et al. (2017). Superiority in Rhesus Macaques of Targeting HIV-1 Env gp140 to CD40 versus LOX-1 in Combination with Replication-Competent NYVAC-KC for Induction of Env-Specific Antibody and T Cell Responses. J Virol 91.

