## Supplementary figures for "Design, immunogenicity, and efficacy of a pan-sarbecovirus dendritic-cell targeting vaccine"

**French Cohort Study Group**

Laurent ABEL (Inserm UMR 1163, Paris, France), Claire ANDREJAK (CHU Amiens, France), François ANGOULVANT (Hôpital Necker, Paris, France), Delphine BACHELET, Krishna BHAVSAR, Lila BOUADMA, Anissa CHAIR, Camille COUFFIGNAL, Charlene DA SILVEIRA, Marie-Pierre DEBRAY, Diane DESCAMPS, Xavier DUVAL, Philippine ELOY, Marina ESPOSITO-FARESE, Nadia ETTALHAOUI, Nathalie GAULT, Jade GHOSN, Isabelle GORENNE, Isabelle HOFFMANN, Ouifiya KAFIF, Sabrina KALI, Antoine KHALIL, Cédric LAOUÉNAN, Samira LARIBI, Minh LE, Quentin LE HINGRAT, François-Xavier LESCURE, Jean Christophe LUCET, France MENTRÉ, Jimmy Mullaert, Nathan PEIFFER-SMADJA, Gilles PEYTAVIN, Carine ROY, Marion SCHNEIDER, Nassima SI MOHAMMED, Lysa TAGHERSET, Coralie TARDIVON, Marie-Capucine TELLIER, Jean-François TIMSIT, Théo TRIOUX, Sarah TUBIANA, Benoit VISSEAUX, Yazdan YAZDANPANAH (Hôpital Bichat, Paris, France), Romain BASMACI, Olivier PICONE (Hôpital Louis Mourier, Colombes, France), Sylvie BEHILILL, Sylvie VAN DER WERF, Vincent ENOUF, Hugo MOUQUET (Pasteur Institute, Paris, France), Marine BELUZE (F-CRIN Partners Platform, Paris, France), Dehbia BENKERROU, Céline DORIVAL, François TÉOULÉ, Amina MEZIANE (Inserm UMR 1136, Paris, France), François BOMPART (Drugs for Neglected Diseases initiative, Geneva, Switzerland), Maude BOUSCAMBERT (Inserm UMR 1111, Lyon, France), Minerva CERVANTES-GONZALEZ, Eric d'ORTENZIO, Oriane PUÉCHAL, Caroline SEMAILLE (REACTing, Paris, France), Catherine CHIROUZE (CHRU Jean Minjoz, Besançon, France), Alexandra COELHO (Inserm UMR 1018, Paris, France), Sandrine COUFFIN-CADIERGUES, Hélène ESPEROU, Ikram HOUAS, Salma JAAFOURA, Aurélie PAPADOPOULOS (Inserm sponsor, Paris, France), Dominique DEPLANQUE (Hôpital Calmette, Lille, France), Mathilde DESVALLÉE, Coralie KHAN (Inserm UMR 1219, Bordeaux, France), Alpha DIALLO, Marie BARTOLI, Soizic LE MESTRE, Noémie MERCIER, Christelle PAUL, Ventzislava PETROV-SANCHEZ (ANRS, Paris, France), Alphonsine DIOUF, Alexandre HOCTIN, Marina MAMBERT (Inserm UMR 1018, Paris, France), François DUBOS (CHU Lille, France), Manuel ETIENNE (CHU Rouen, France), Alexandre GAYMARD (Inserm UMR 1111, Lyon, France), Tristan GIGANTE, Morgane GILG, Bénédicte ROSSIGNOL (F-CRIN INI-CRCT, Nancy, France), Jérémie GUEDJ, Hervé LE NAGARD, Guillaume LINGAS, Nadège NEANT (Inserm UMR 1137, Paris, France), Jean-Sébastien HULOT (Hôpital Européen Georges Pompidou, Paris, France), Florentia KAGUELIDOU, Justine PAGES (Hôpital Robert Debré, Paris, France), Yves LEVY, Aurélie WIEDEMANN (Vaccine Research Insitute (VRI), Inserm UMR 955, Créteil, France), Claire LEVY-MARCHAL (F-CRIN INI-CRCT, Paris, France), Bruno LINA, Manuel ROSA-CALATRAVA, Olivier TERRIER (Inserm UMR 1111, Lyon, France), Denis MALVY (CHU Bordeaux, France), Marion NORET (RENARCI, Annecy, France), Patrick ROSSIGNOL (CHU Nancy, France), Christelle TUAL, Aurélie VEISLINGER (Inserm CIC-1414, Rennes, France), Noémie VANEL (Hôpital la Timone, Marseille, France)

**Table S1. CD40.CoV2 vaccine region characteristics**

| Characteristics | vS1 (S125-250) | vRBD (S318-541) | vS2 (S1056-1209) | vN2 (N276-411) | Global |
|---|---|---|---|---|---|
| Lengh (aa) | 126 | 224 | 154 | 136 | 640 |
| Number of potential SARS-CoV-2 CD8 T-cell epitopes (predicted) | 535 | 908 | 458 | 412 | 2313 |
| % Class-I Coverage (A, B, C) | 99% (100-97-100) | 99% (100-97-100) | 98% (100-94-100) | 96% (97-94-100) | 100% |
| Number of potential SARS-CoV-2 CD4 T-cell epitopes (predicted) | 712 | 1166 | 548 | 559 | 2985 |
| % Class-II HLA coverage (DR, DP, DQ) | 89% (96-100-75) | 100% | 87% (84-67-100) | 83% (88-44-95) | 100% |
| Number of potential SARS-CoV-2 linear B-cell epitopes (predicted) | 3 | 8 | 3 | 3 | 17 |

**Table S4. Vaccine region homology between β and α coronaviruses**

| Virus strain | Accession number | Genus | Subgenus | Virus type | SARS-CoV-2 (MN908947.3) homology (%) | | | |
|---|---|---|---|---|---|---|---|---|
| | | | | | vS1 (125-250) | vRBD (318-541) | vS2 (1056-1209) | vN2 (276-411) |
| SARS-CoV-2 VOC Alpha (UK) | MZ344997.1 | Betacoronavirus | Sarbecovirus | SARS-CoV-2 | 99,2 | 99,6 | 99,4 | 100,0 |
| SARS-CoV-2 VOC beta (SA) | MW598419.1 | Betacoronavirus | Sarbecovirus | SARS-CoV-2 | 96,8 | 98,7 | 100,0 | 100,0 |
| SARS-CoV-2 VOC Gamma (P.1 Brazil) | MZ169911.1 | Betacoronavirus | Sarbecovirus | SARS-CoV-2 | 98,4 | 98,7 | 99,4 | 100,0 |
| SARS-CoV-2 VOC Delta (India) | MZ359841.1 | Betacoronavirus | Sarbecovirus | SARS-CoV-2 | 96,8 | 99,1 | 100,0 | 99,3 |
| Bat SC2r-CoV RaTG13 | MN996532 | Betacoronavirus | Sarbecovirus | SC2r-CoV | 99,2 | 90,2 | 99,4 | 100,0 |
| Bat SC2r-CoV RacCS203 | MW251308 | Betacoronavirus | Sarbecovirus | SC2r-CoV | 38,1 | 63,8 | 98,1 | 96,3 |
| Pangolin SC2r-CoV GX-P4L | MT040333 | Betacoronavirus | Sarbecovirus | SC2r-CoV | 92,1 | 86,6 | 97,4 | 94,9 |
| Pangolin SC2r-CoV GX-P5L | MT040335 | Betacoronavirus | Sarbecovirus | SC2r-CoV | 92,1 | 86,6 | 98,1 | 95,6 |
| Bat SC2r-CoV ZC45 | MG772933 | Betacoronavirus | Sarbecovirus | SC2r-CoV | 57,1 | 65,6 | 95,5 | 96,3 |
| Bat SC2r-CoV ZXC21 | MG772934 | Betacoronavirus | Sarbecovirus | SC2r-CoV | 56,3 | 66,1 | 95,5 | 96,3 |
| Bat SC2r-CoV Rc-o319 | LC556375 | Betacoronavirus | Sarbecovirus | SC2r-CoV | 36,5 | 73,7 | 90,9 | 94,1 |
| Bat SC1r-CoV WIV1 | KF367457 | Betacoronavirus | Sarbecovirus | SC1r-CoV | 46,0 | 74,6 | 92,9 | 91,9 |
| Bat CoV WIV16 | KT444582 | Betacoronavirus | Sarbecovirus | SC1r-CoV | 45,2 | 74,6 | 92,9 | 91,9 |
| Bat CoV Rs3367 | KC881006 | Betacoronavirus | Sarbecovirus | SC1r-CoV | 46,0 | 74,6 | 92,2 | 91,9 |
| Bat CoV LYRa11 | KF569996 | Betacoronavirus | Sarbecovirus | SC1r-CoV | 46,8 | 72,8 | 92,9 | 91,9 |
| Bat SC1r-CoV Cp/Yunnan2011 | JX993988 | Betacoronavirus | Sarbecovirus | SC1r-CoV | 42,1 | 64,3 | 94,2 | 92,6 |
| Bat CoV Rs-YN2018B | MK211376 | Betacoronavirus | Sarbecovirus | SC1r-CoV | 45,2 | 74,1 | 92,9 | 91,9 |
| Bat CoV Rs7327 | KY417151 | Betacoronavirus | Sarbecovirus | SC1r-CoV | 46,0 | 74,1 | 93,5 | 91,9 |
| Bat CoV RsSHC014 | KC881005 | Betacoronavirus | Sarbecovirus | SC1r-CoV | 46,0 | 75,0 | 92,9 | 91,9 |
| Bat CoV Rs4231 | KY417146 | Betacoronavirus | Sarbecovirus | SC1r-CoV | 45,2 | 75,0 | 92,2 | 91,9 |
| Bat CoV Rs4084 | KY417144 | Betacoronavirus | Sarbecovirus | SC1r-CoV | 46,0 | 74,6 | 92,9 | 91,2 |
| Bat CoV Rs4081 | KY417143.1 | Betacoronavirus | Sarbecovirus | SC1r-CoV | 41,3 | 64,3 | 94,8 | 91,9 |
| Bat CoV Rs672 | FJ588686.1 | Betacoronavirus | Sarbecovirus | SC1r-CoV | 40,5 | 63,8 | 94,8 | 91,9 |
| Bat CoV Rs4237 | KY417147.1 | Betacoronavirus | Sarbecovirus | SC1r-CoV | 42,1 | 64,7 | 94,8 | 91,9 |
| Bat SC1r-CoV YNLF_31C | KP886808 | Betacoronavirus | Sarbecovirus | SC1r-CoV | 43,7 | 65,2 | 92,9 | 91,2 |
| Bat SC1r-CoV Rp3 | DQ071615 | Betacoronavirus | Sarbecovirus | SC1r-CoV | 42,1 | 65,2 | 93,5 | 91,9 |
| BtRl-BetaCoV/SC2018 | MK211374.1 | Betacoronavirus | Sarbecovirus | SC1r-CoV | 39,7 | 65,6 | 94,2 | 91,2 |
| Bat SC1r-CoV Rf1 | DQ412042 | Betacoronavirus | Sarbecovirus | SC1r-CoV | 44,4 | 65,2 | 92,2 | 90,4 |
| Bat SC1r-CoV HeB2013 | KJ473812 | Betacoronavirus | Sarbecovirus | SC1r-CoV | 44,4 | 65,2 | 91,6 | 91,9 |
| Bat SC1r-CoV Rp/Shaanxi2011 | JX993987 | Betacoronavirus | Sarbecovirus | SC1r-CoV | 40,5 | 66,1 | 92,9 | 91,2 |
| Bat SC1r-CoV Rm1 | DQ412043 | Betacoronavirus | Sarbecovirus | SC1r-CoV | 39,7 | 65,2 | 94,2 | 89,7 |
| Bat SC1r-CoV HuB2013 | KJ473814 | Betacoronavirus | Sarbecovirus | SC1r-CoV | 40,5 | 65,6 | 94,2 | 91,9 |
| Bat SC1r-CoV HKU3-1 | DQ022305 | Betacoronavirus | Sarbecovirus | SC1r-CoV | 41,3 | 65,2 | 94,8 | 91,9 |
| Bat SC1r-CoV Longquan-140 | KF294457 | Betacoronavirus | Sarbecovirus | SC1r-CoV | 42,1 | 64,7 | 94,8 | 91,9 |
| Bat SC1r-CoV BM48-31/BGR/2008 | NC_014470 | Betacoronavirus | Sarbecovirus | SCr-Cov | 36,5 | 68,8 | 86,4 | 93,4 |
| Bat SC1r-CoV BtKY72 | KY352407 | Betacoronavirus | Sarbecovirus | SCr-Cov | 38,1 | 73,2 | 87,0 | 93,4 |
| SARS-CoV-1 (Tor2) | NC_004718 | Betacoronavirus | Sarbecovirus | SARS-CoV-1 | 44,4 | 72,8 | 93,5 | 91,9 |
| OC43 (strain ATCC VR-759) | NC_006213 | Betacoronavirus | Embecovirus | CoV | 14,3 | 20,1 | 38,3 | 25,2 |
| HKU1 | NC_006577 | Betacoronavirus | Embecovirus | CoV | 13,5 | 19,2 | 32,1 | 23,7 |
| MERS (HCoV-EMC/2012) | NC_019843 | Betacoronavirus | Merbecovirus | MERS-CoV | 14,7 | 16,1 | 31,9 | 37,0 |
| NL63 | NC_005831 | Alphacoronavirus | Setracovirus | CoV | 13,5 | 13,8 | 22,2 | 19,9 |
| 229E | NC_002645 | Alphacoronavirus | Duvinacovirus | CoV | 6,3 | 12,9 | 25,1 | 16,2 |

**Table S5. Vaccine CD4 and CD8 T-cell epitopes with 100% homology with 38 sarbecoviruses**

| Position | Sequence | HLA restriction | Type |
|---|---|---|---|
| S1056-1063 | APHGVVFL | B*07 | CD8 |
| S1089-1096 | FPREGVFV | B*5101 | CD8 |
| S1137-1145 | VYDPLQPEL | A*2402 | CD8 |
| N305-314 | AQFAPSASAF | B*1501 | CD8 |
| N306-315 | QFAPSASAFF | A*2402 | CD8 |
| N307-315 | FAPSASAFF | B*3501 | CD8 |
| N308-317 | APSASAFFGM | B*0702 | CD8 |
| N310-319 | SASAFFGMSR | A*6801 | CD8 |
| N311-319 | ASAFFGMSR | A*1101 / A*6801 | CD8 |
| N301-315 | WPQIAQFAPSASAFF | | CD4 |
| N306-320 | QFAPSASAFFGMSRI | | CD4 |

**Figure S1. Clinical symptom scores for CD40.CoV2 vaccinated and mock-vaccinated hCD40/K18-hACE2 mice from days 0 to 12 post-inoculation with SARS-CoV-2.** The clinical score, ranging from 0 to 4, was monitored daily until day 12 post-inoculation (dpi) and was calculated according to the presence of four symptoms: eye closure, ruffled fur, hunched posture, and labored breathing.

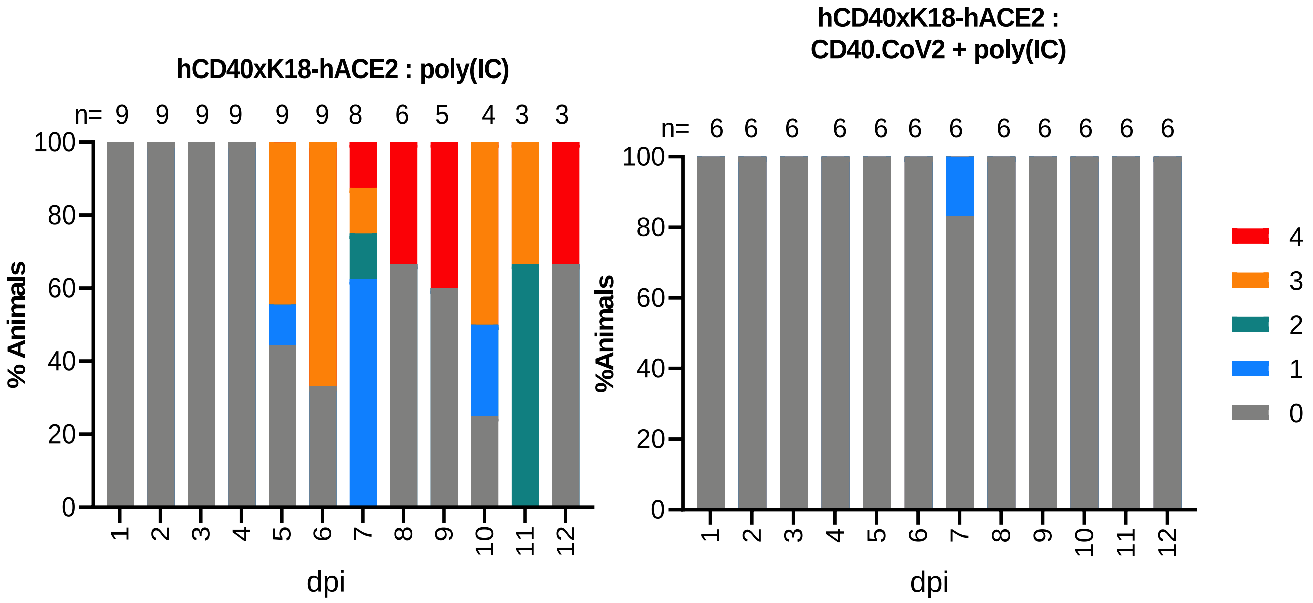

**Figure S2. IgG responses against MERS and common cold coronaviruses in CD40.CoV2 vaccinated and mock-vaccinated hCD40/K18-hACE2 mice.** Level of IgG antibodies (AU) binding to spike proteins from MERS and common cold coronaviruses in vaccinated and mock-vaccinated animals before vaccination (baseline, -2 days post-vaccination [dpv], n = 9-12 animals per group), after completion of the vaccination schedule (28 dpv, n = 9-12 animals per group), and 40 dpv (i.e., day 12 pi time point, n = 3-5 animals per group). The medians [Min-Max] are shown. Grey dashed lines represent the prime and boost vaccination. The red dashed line represents SARS-CoV2 inoculation.

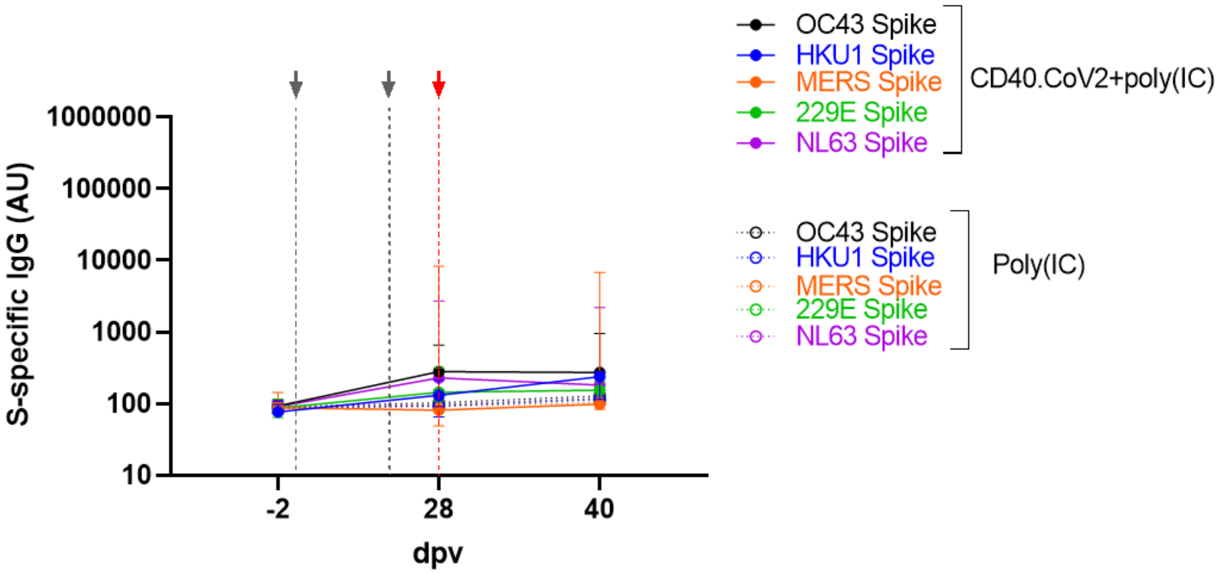

**Figure S3. Induction of cross-reactive and neutralizing IgG antibody responses by the CD40.CoV2 vaccine in a second replicated animal experiment.** Level of IgG antibodies (AU) binding to Wuhan and VOC SARS-CoV-2 RBD proteins (A) and Spike proteins from MERS and common cold coronaviruses (B) in mock-vaccinated and vaccinated animals before vaccination (baseline, -2 days post-vaccination [dpv], n = 10-12 animals per group), after completion of the vaccination schedule (28 dpv, n = 10-12 animals per group), and 40 dpv (i.e., day 12 pi time point, n = 3 animals per group). (C) Levels of IgG antibodies (AU) binding to SARS-CoV-2 (solid lines) and SARS-CoV-1 (dashed lines) S proteins in mock-vaccinated (grey) and vaccinated (blue) animals at -2, 28, and 40 dpv. Medians [Min-Max] are shown. Grey dashed lines represent the prime and boost vaccination. The red dashed line represents SARS-CoV-2 inoculation. Neutralizing activity of RBD antibodies (Unit/mL) (D) and S antibodies (Unit/mL) (E) in mock-vaccinated (grey) and vaccinated (blue) animals post-vaccination (open circle) and post-infection (solid circle). Because mock-vaccinated animals have no SARS-CoV-2 antibody responses before infection, neutralization activity was evaluated in sera from only three mock-vaccinated animals for which samples before and after infection were available. Medians ± IQRs are shown. Thirty plasma samples from unvaccinated mice were used to determine the positivity threshold, defined as the whole units/mL value immediately above the concentration of the highest sample for RBD (i.e., 8 units/mL) and Spike (i.e., 4 units/mL) proteins.

.

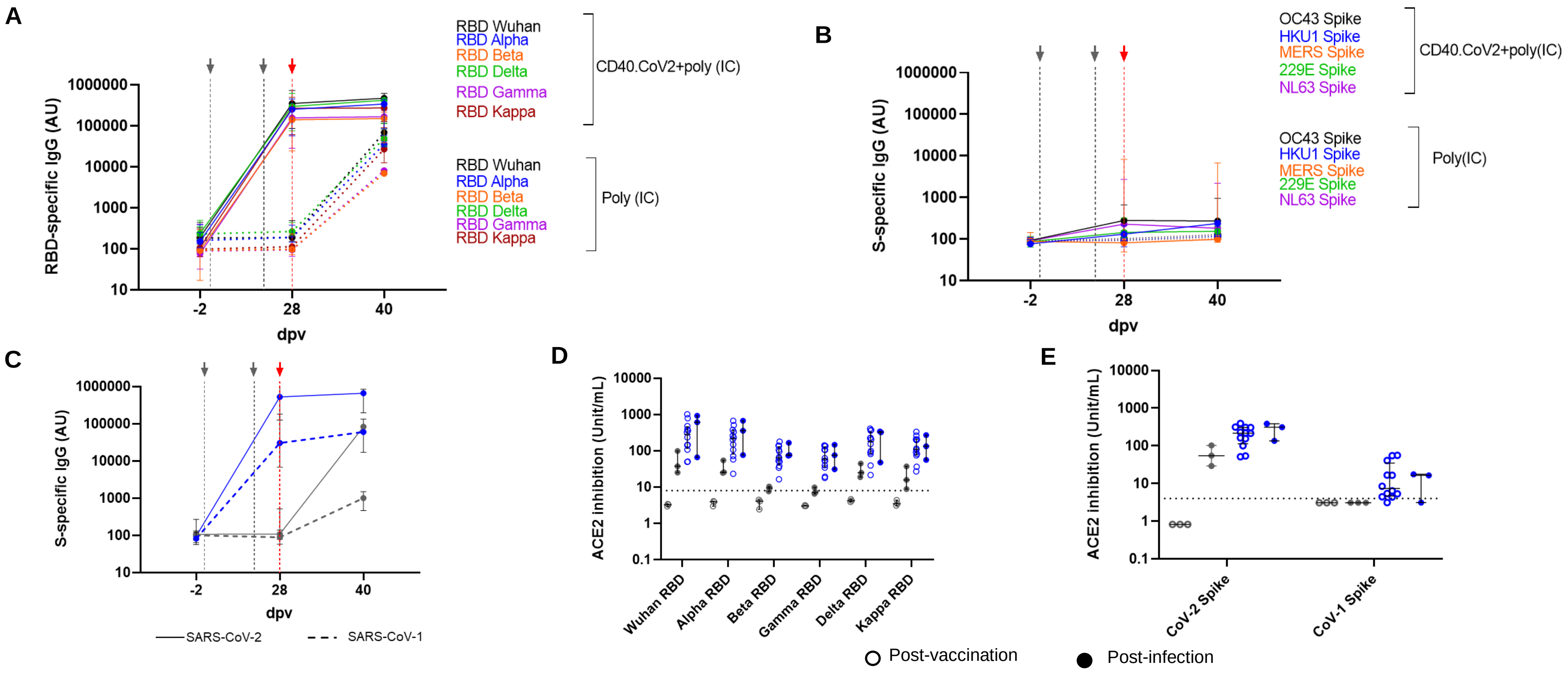

**Figure S4. Production of TNF and IL-2 by SARS-CoV-2-specific CD4⁺ and CD8⁺ T cells from convalescent COVID-19 patients after *in-vitro* stimulation with the CD40.CoV2 vaccine.** Frequency of TNF⁺ and IL-2⁺ SARS-CoV2 specific CD4⁺ T cells (A) or specific CD8⁺ T cells (B) from convalescent COVID-19 patients (n = 14) stimulated or not with the CD40.CoV2 vaccine (1 nM) on D0 and re-stimulated with various vOLPs (vRBD, vS1, vS2, or vN2) or cont.OLPs (Gpz or N1-N2, grey) on D8 (1 µg/ml). Median values ± IQRs are shown. Friedman and Dunn's multiple comparison tests were used for statistical analysis (*P < 0.05, **P < 0.01, ***P < 0.001, ****P < 0.0001, ns: not significant).

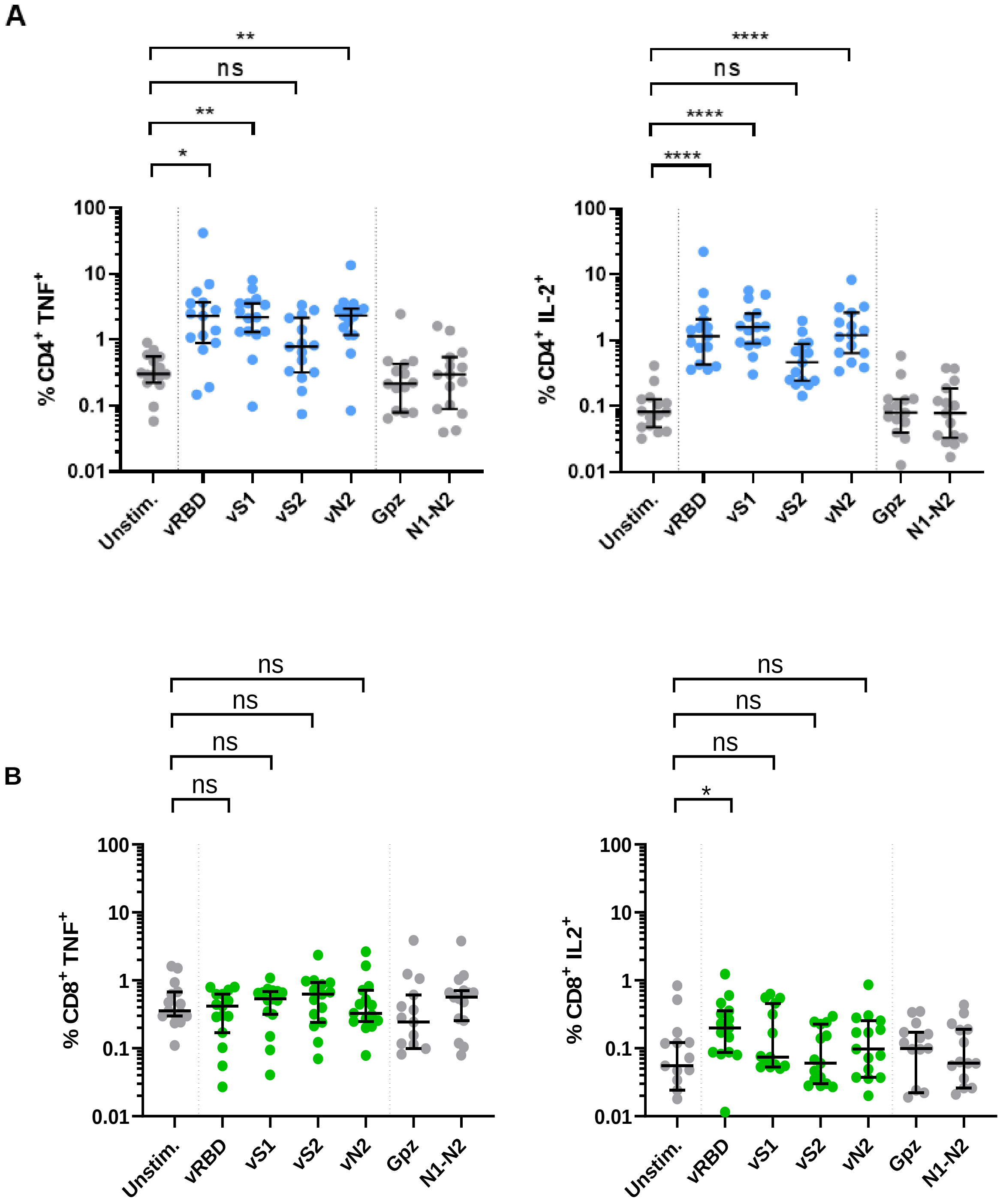

**Figure S5. Simple linear regression between SARS-CoV-1- and SARS-CoV-2-specific CD4⁺ and CD8⁺ T-cell responses.** Correlation between the frequency of total cytokines (IFN-γ ± IL-2 ± TNF) secreted by specific CD4⁺ (A) or CD8⁺ (B) T cells after stimulation with the CD40.CoV2 vaccine (1 nM) on D0 and re-stimulation with OLPs representing the sequence of S1, vRBD, and vN2 from either the SARS CoV-1 or SARS CoV-2 virus (1 µg/mL). Pearson's test was used for statistical significance. Simple linear regression is represented as a solid line and the 95% confidence interval as dashed lines.

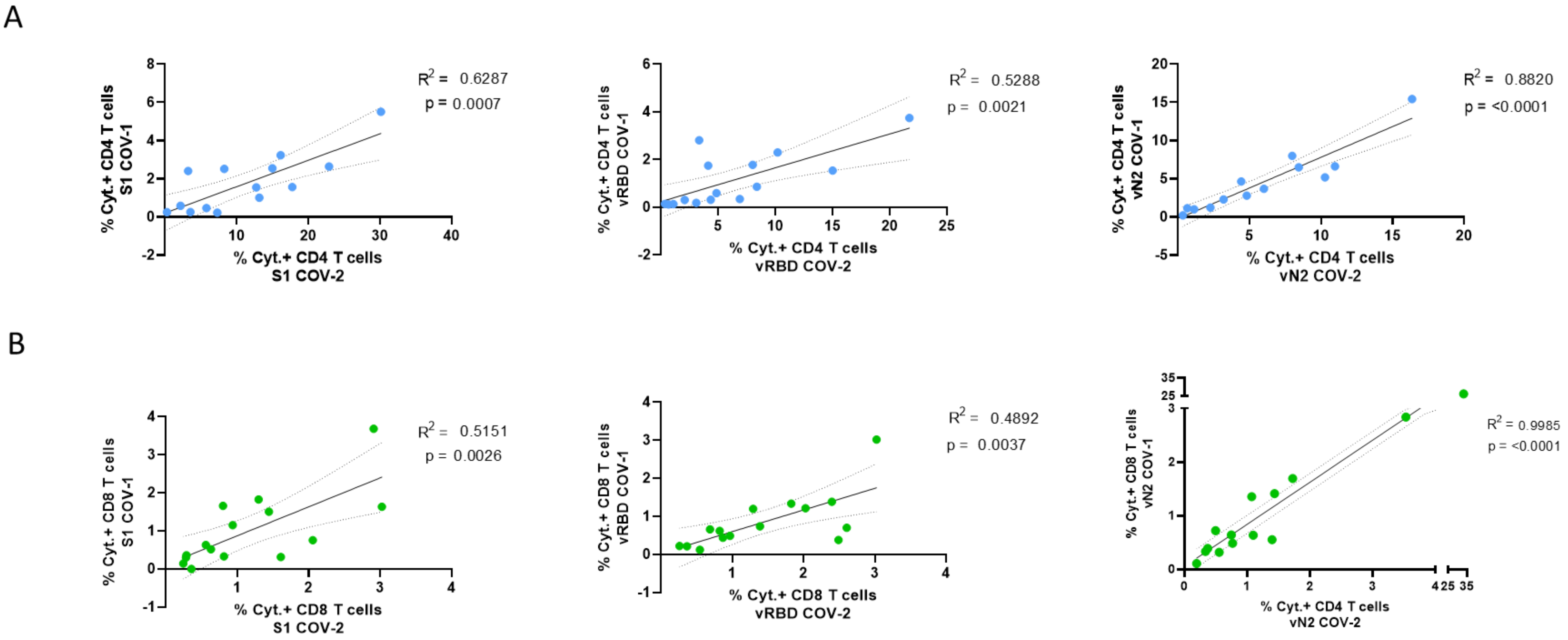

### Table S2

| Vaccine Antigen | Position | Sequence | HLA restriction | Source | B.1.1.7 (Alpha Variant) | B.1.351 (Beta Variant) | P.1 (Gamma Variant) | B.1.617.2 (Delta Variant) | SARS-CoV-1 | Sequence homology (%) with SARS-CoV-1 | Sequence homology (mean %) with 38 sarbecoviruses |
|---|---|---|---|---|---|---|---|---|---|---|---|
| vS1 | S142-150 | GVYYHKNNK | A*0301 | Tarke et al., Cell Rep Medicine, 2021 | GVY---HKNNK | GVY--YHKNNK | GVY--YHKNNK | DVY--YHKNNK | AVSKPMGTQTH | 11,1 | 30,0 |
| | S144-152 | YYHKNNKSW | A*2402 | Tarke et al., Cell Rep Medicine, 2021 | Y---HKNNKSW | Y--YHKNNKSW | Y--YHKNNKSW | Y--YHKNNKSW | SKPMGTQTHTM | 0,0 | 33,6 |
|  | S151-160 | SWMESEFRVY | A*2902 | Tarke et al., Cell Rep Medicine, 2021 | SWMESEFRVY | SWMESEFRVY | SWMESEFRVY | SWMESG--VY | TM------IF | 0,0 | 37,8 |
| | S159-168 | VYSSANNCTF | A*2402 | Tarke et al., Cell Rep Medicine, 2021 | VYSSANNCTF | VYSSANNCTF | VYSSANNCTF | VYSSANNCTF | IFDNAFNCTF | 50,0 | 67,8 |
| | S162-170 | SANNCTFEY | A*2902<br>B*3501 | Tarke et al., Cell Rep Medicine, 2021<br>Tarke et al., Cell Rep Medicine, 2021 | SANNCTFEY | SANNCTFEY | SANNCTFEY | SANNCTFEY | NAFNCTFEY | 77,8 | 68,8 |
| | S168-177 | FEYVSQPFLM | B*4001 | Tarke et al., Cell Rep Medicine, 2021 | FEYVSQPFLM | FEYVSQPFLM | FEYVSQPFLM | FEYVSQPFLM | FEYISDAFSL | 50,0 | 46,2 |
| | S169-177 | EYVSQPFLM | A*2402 | Hu et al., Genes & Diseases, 2021<br>Tarke et al., Cell Rep Medicine, 2021 | EYVSQPFLM | EYVSQPFLM | EYVSQPFLM | EYVSQPFLM | EYISDAFSL | 44,4 | 45,6 |
| | S192-200 | FVFKNIDGY | A*2902<br>A*2601 | Tarke et al., Cell Rep Medicine, 2021<br>Tarke et al., Cell Rep Medicine, 2021 | FVFKNIDGY | FVFKNIDGY | FVFKNIDGY | FVFKNIDGY | FVFKNKDGF | 77,8 | 73,9 |
| | S193-201 | VFKNIDGYF | A*2402 | Hu et al., Genes & Diseases, 2021<br>Tarke et al., Cell Rep Medicine, 2021 | VFKNIDGYF | VFKNIDGYF | VFKNIDGYF | VFKNIDGYF | VFKNKDGFL | 66,7 | 69,4 |
| | S202-210 | KIYSKHTPI | B*0801 | Tarke et al., Cell Rep Medicine, 2021 | KIYSKHTPI | KIYSKHTPI | KIYSKHTPI | KIYSKHTPI | YVYKGYQPI | 33,3 | 50,8 |
| | S208-216 | TPINLVRDL | B*0702 | Tarke et al., Cell Rep Medicine, 2021 | TPINLVRDL | TPINLVRGL | TPINLVRDL | TPINLVRDL | QPIDVVRDL | 66,7 | 54,7 |
| | S216-223 | LPQGFSAL | B*0801<br>B*0702 | Tarke et al., Cell Rep Medicine, 2021<br>Tarke et al., Cell Rep Medicine, 2021 | LPQGFSAL | LPQGFSAL | LPQGFSAL | LPQGFSAL | LPSGFNTL | 62,5 | 72,0 |
| | S224-231 | EPLVDLPI | B*5101 | Tarke et al., Cell Rep Medicine, 2021 | EPLVDLPI | EPLVDLPI | EPLVDLPI | EPLVDLPI | KPIFKLPL | 37,5 | 52,4 |
| | S229-238 | LPIGINITRF | B*0702<br>B*3501 | Tarke et al., Cell Rep Medicine, 2021<br>Tarke et al., Cell Rep Medicine, 2021 | LPIGINITRF | LPIGINITRF | LPIGINITRF | LPIGINITRF | LPLGINITNF | 80,0 | 79,7 |
| | S233-241 | INITRFQTL | B*0801 | Tarke et al., Cell Rep Medicine, 2021 | INITRFQTL | INITRFQTL | INITRFQTL | INITRFQTL | INITNFRAI | 55,6 | 65,5 |
| | S234-241 | NITRFQTL | B*0801 | Tarke et al., Cell Rep Medicine, 2021 | NITRFQTL | NITRFQTL | NITRFQTL | NITRFQTL | NITNFRAI | 50,0 | 62,8 |
| vRBD | S321-329 | QPTESIVRF | B*3501<br>B*0702 | Tarke et al., Cell Rep Medicine, 2021<br>Tarke et al., Cell Rep Medicine, 2021 | QPTESIVRF | QPTESIVRF | QPTESIVRF | QPTESIVRF | VPSGDVVRF | 44,4 | 58,0 |
| | S323-335 | TESIVRFPNITNL | B*4001 | Tarke et al., Cell Rep Medicine, 2021 | TESIVRFPNITNL | TESIVRFPNITNL | TESIVRFPNITNL | TESIVRFPNITNL | SGDVVRFPNITNL | 69,2 | 73,4 |
| | S328-338 | RFPNITNLCPF | A*2402 | Tarke et al., Cell Rep Medicine, 2021 | RFPNITNLCPF | RFPNITNLCPF | RFPNITNLCPF | RFPNITNLCPF | RFPNITNLCPF | 100,0 | 96,3 |
| | S339-347 | GEVFNATRF | B*4402 / B*4403<br>B*4001 | Tarke et al., Cell Rep Medicine, 2021<br>Tarke et al., Cell Rep Medicine, 2021 | GEVFNATRF | GEVFNATRF | GEVFNATRF | GEVFNATRF | GEVFNATKF | 88,9 | 80,5 |
| | S340-351 | EVFNATRFASVY | A*2601 | Tarke et al., Cell Rep Medicine, 2021 | EVFNATRFASVY | EVFNATRFASVY | EVFNATRFASVY | EVFNATRFASVY | EVFNATKFPSVY | 83,3 | 81,3 |
| | S343-351 | NATRFASVY | B*3501 | Tarke et al., Cell Rep Medicine, 2021 | NATRFASVY | NATRFASVY | NATRFASVY | NATRFASVY | NATKFPSVY | 77,8 | 80,5 |
| | S346-355 | RFASVYAWNR | A*3101 | Tarke et al., Cell Rep Medicine, 2021 | RFASVYAWNR | RFASVYAWNR | RFASVYAWNR | RFASVYAWNR | KFPSVYAWER | 70,0 | 77,8 |
| | S347-355 | FASVYAWNR | A*6801 | Tarke et al., Cell Rep Medicine, 2021 | FASVYAWNR | FASVYAWNR | FASVYAWNR | FASVYAWNR | FPSVYAWER | 77,8 | 80,2 |
| | S349-357 | SVYAWNRKR | A*0301 | Tarke et al., Cell Rep Medicine, 2021 | SVYAWNRKR | SVYAWNRKR | SVYAWNRKR | SVYAWNRKR | SVYAWERKK | 77,8 | 77,8 |
| | S351-358 | YAWNRKRI | B*5101 | Tarke et al., Cell Rep Medicine, 2021 | YAWNRKRI | YAWNRKRI | YAWNRKRI | YAWNRKRI | YAWERKKI | 75,0 | 77,7 |
| | S361-369 | CVADYSVLY | A*2902<br>A*2601 | Tarke et al., Cell Rep Medicine, 2021<br>Tarke et al., Cell Rep Medicine, 2021 | CVADYSVLY | CVADYSVLY | CVADYSVLY | CVADYSVLY | CVADYSVLY | 100,0 | 91,9 |
| | S370-378 | NSASFSTFK | A*6801 | Tarke et al., Cell Rep Medicine, 2021 | NSASFSTFK | NSASFSTFK | NSASFSTFK | NSASFSTFK | NSTFFSTFK | 77,8 | 88,0 |
| | S378-386 | KCYGVSPTK | A*0301 | Ferretti et al., Immunity, 2020<br>Tarke et al., Cell Rep Medicine, 2021 | KCYGVSPTK | KCYGVSPTK | KCYGVSPTK | KCYGVSPTK | KCYGVSATK | 88,9 | 89,8 |
| | S386-395 | KLNDLCFTNV | A*0201 | Poran et al., Genome Medicine, 2020 | KLNDLCFTNV | KLNDLCFTNV | KLNDLCFTNV | KLNDLCFTNV | KLNDLCFSNV | 90,0 | 87,0 |
| | S394-403 | NVYADSFVIR | A*6801 | Tarke et al., Cell Rep Medicine, 2021 | NVYADSFVIR | NVYADSFVIR | NVYADSFVIR | NVYADSFVIR | NVYADSFVVK | 80,0 | 77,0 |
| | S400-408 | FVIRGDEVR | A*6801 | Tarke et al., Cell Rep Medicine, 2021 | FVIRGDEVR | FVIRGDEVR | FVIRGDEVR | FVIRGDEVR | FVVKGDDVR | 66,7 | 72,1 |
| | S409-417 | QIAPGQTGK | A*6801 | Tarke et al., Cell Rep Medicine, 2021 | QIAPGQTGK | QIAPGQTGN | QIAPGQTGT | QIAPGQTGK | QIAPGQTGV | 88,9 | 81,1 |
| | S417-425 | KIADYNYKL | A*0201 | Shomuradova et al., Immunity, 2020 | KIADYNYKL | NIADYNYKL | TIADYNYKL | KIADYNYKL | VIADYNYKL | 88,9 | 89,8 |
| | S424-433 | KLPDDFTGCV | A*0201 | Shomuradova et al., Immunity, 2020 | KLPDDFTGCV | KLPDDFTGCV | KLPDDFTGCV | KLPDDFTGCV | KLPDDFMGCV | 90,0 | 97,8 |
| | S442-451 | DSKVGGNYNY | A*2601 | Tarke et al., Cell Rep Medicine, 2021 | DSKVGGNYNY | DSKVGGNYNY | DSKVGGNYNY | DSKVGGNYNY | DATSTGNYNY | 60,0 | 45,1 |
| | S444-453 | KVGGNYNYLY | A*2902 | Tarke et al., Cell Rep Medicine, 2021 | KVGGNYNYLY | KVGGNYNYLY | KVGGNYNYLY | KVGGNYNYRY | TSTGNYNYKY | 60,0 | 45,1 |
| | S445-453 | VGGNYNYLY | A*2902 | Tarke et al., Cell Rep Medicine, 2021 | VGGNYNYLY | VGGNYNYLY | VGGNYNYLY | VGGNYNYRY | STGNYNYKY | 66,7 | 48,3 |
| | S448-456 | NYNYLYRLF | A*2402 | Hu et al., Genes & Diseases, 2021<br>Gao et al., IScience, 2021<br>Motozono et al. Cell Host & Microbes, 2021<br>Kared et al., JCI, 2021<br>Tarke et al., Cell Rep Medicine, 2021 | NYNYLYRLF | NYNYLYRLF | NYNYLYRLF | NYNYRYRLF | NYNYKYRYL | 66,7 | 55,6 |
| | S449-457 | YNYLYRLFR | A*3101 | Tarke et al., Cell Rep Medicine, 2021 | YNYLYRLFR | YNYLYRLFR | YNYLYRLFR | YNYRYRLFR | YNYKYRYLR | 66,7 | 61,0 |
| | S454-462 | RLFRKSNLK | A*11<br>A*3101<br>A*0301 | Kared et al., JCI, 2021<br>Tarke et al., Cell Rep Medicine, 2021<br>Tarke et al., Cell Rep Medicine, 2021 | RLFRKSNLK | RLFRKSNLK | RLFRKSNLK | RLFRKSNLK | RYLRHGKLR | 33,3 | 56,2 |
| | S458-466 | KSNLKPFER | A*3101 | Tarke et al., Cell Rep Medicine, 2021 | KSNLKPFER | KSNLKPFER | KSNLKPFER | KSNLKPFER | HGKLRPFER | 55,6 | 71,5 |
| | S462-472 | KPFERDISTEI | B*0702 | Tarke et al., Cell Rep Medicine, 2021 | KPFERDISTEI | KPFERDISTEI | KPFERDISTEI | KPFERDISTEI | RPFERDISNVP | 63,6 | 69,3 |
| | S464-473 | FERDISTEIY | B*4403 | Tarke et al., Cell Rep Medicine, 2021 | FERDISTEIY | FERDISTEIY | FERDISTEIY | FERDISTEIY | FERDISNVPF | 60,0 | 62,4 |
| | S464-472 | FERDISTEI | B*4001 | Tarke et al., Cell Rep Medicine, 2021 | FERDISTEI | FERDISTEI | FERDISTEI | FERDISTEI | FERDISNVP | 66,7 | 65,8 |
| | S495-503 | YGFQPTNGV | B*5101 | Tarke et al., Cell Rep Medicine, 2021 | YGFQPTYGV | YGFQPTYGV | YGFQPTYGV | YGFQPTNGV | YGFYTTTGI | 55,6 | 55,0 |
| | S506-515 | QPYRVVVLSF | B*0702 | Tarke et al., Cell Rep Medicine, 2021 | QPYRVVVLSF | QPYRVVVLSF | QPYRVVVLSF | QPYRVVVLSF | QPYRVVVLSF | 100,0 | 90,3 |
| | S506-513 | QPYRVVVL | B*0801<br>B*0702 | Tarke et al., Cell Rep Medicine, 2021<br>Tarke et al., Cell Rep Medicine, 2021 | QPYRVVVL | QPYRVVVL | QPYRVVVL | QPYRVVVL | QPYRVVVL | 100,0 | 87,8 |
| vS2 | S1056-1063 | APHGVVFL | B*07 | Kared et al., JCI, 2021 | APHGVVFL | APHGVVFL | APHGVVFL | APHGVVFL | APHGVVFL | 100,0 | 100,0 |
| | S1059-1067 | GVVFLHVTY | A*3201 | Tarke et al., Cell Rep Medicine, 2021 | GVVFLHVTY | GVVFLHVTY | GVVFLHVTY | GVVFLHVTY | GVVFLHVTY | 100,0 | 99,7 |
| | S1060-1068 | VVFLHVTYV | A*02<br>A*0201<br>A*0206 | Kared et al., JCI, 2021<br>Tarke et al., Cell Rep Medicine, 2021<br>Tarke et al., Cell Rep Medicine, 2021 | VVFLHVTYV | VVFLHVTYV | VVFLHVTYV | VVFLHVTYV | VVFLHVTYV | 100,0 | 98,8 |
| | S1064-1073 | HVTYVPAQEK | A*6801 | Tarke et al., Cell Rep Medicine, 2021 | HVTYVPAQEK | HVTYVPAQEK | HVTYVPAQEK | HVTYVPAQEK | HVTYVPSQER | 80,0 | 85,4 |
| | S1065-1073 | VTYVPAQEK | A*3001<br>A*0301 | Tarke et al., Cell Rep Medicine, 2021<br>Tarke et al., Cell Rep Medicine, 2021 | VTYVPAQEK | VTYVPAQEK | VTYVPAQEK | VTYVPAQEK | VTYVPSQER | 77,8 | 83,8 |
| | S1089-1096 | FPREGVFV | B*5101 | Tarke et al., Cell Rep Medicine, 2021 | FPREGVFV | FPREGVFV | FPREGVFV | FPREGVFV | FPREGVFV | 100,0 | 100,0 |
| | S1091-1102 | REGVFVSNGTHW | B*4403 | Tarke et al., Cell Rep Medicine, 2021 | REGVFVSNGTHW | REGVFVSNGTHW | REGVFVSNGTHW | REGVFVSNGTHW | REGVFVFNGTSW | 83,3 | 91,4 |
| | S1093-1102 | GVFVSNGTHW | B*5701 | Tarke et al., Cell Rep Medicine, 2021 | GVFVSNGTHW | GVFVSNGTHW | GVFVSNGTHW | GVFVSNGTHW | GVFVFNGTSW | 80,0 | 89,7 |
| | S1094-1103 | VFVSNGTHWF | A*2402 | Tarke et al., Cell Rep Medicine, 2021 | VFVSNGTHWF | VFVSNGTHWF | VFVSNGTHWF | VFVSNGTHWF | VFVFNGTSWF | 80,0 | 89,7 |
| | S1095-1103 | FVSNGTHWF | A*2601<br>B*3501 | Tarke et al., Cell Rep Medicine, 2021<br>Tarke et al., Cell Rep Medicine, 2021 | FVSNGTHWF | FVSNGTHWF | FVSNGTHWF | FVSNGTHWF | FVFNGTSWF | 77,8 | 88,6 |
| | S1099-1107 | GTHWFVTQR | A*11<br>A*3101 | Kared et al., JCI, 2021<br>Tarke et al., Cell Rep Medicine, 2021 | GTHWFVTQR | GTHWFVTQR | GTHWFVTQR | GTHWFVTQR | GTSWFITQR | 77,8 | 83,5 |
| | S1101-1109 | HWFVTQRNF | A*2402 | Hu et al., Genes & Diseases, 2021<br>Tarke et al., Cell Rep Medicine, 2021 | HWFVTQRNF | HWFVTQRNF | HWFVTQRNF | HWFVTQRNF | SWFITQRNF | 77,8 | 84,1 |
| | S1102-1110 | WFVTQRNFY | A*2902 | Tarke et al., Cell Rep Medicine, 2021 | WFVTQRNFY | WFVTQRNFY | WFVTQRNFY | WFVTQRNFY | WFITQRNFF | 77,8 | 88,0 |
| | S1136-1149 | TVYDPLQPELDSFK | A*0301 | Tarke et al., Cell Rep Medicine, 2021 | TVYDPLQPELDSFK | TVYDPLQPELDSFK | TVYDPLQPELDSFK | TVYDPLQPELDSFK | TVYDPLQPELDSFK | 100,0 | 99,6 |
| | S1137-1148 | VYDPLQPELDSF | A*2402 | Tarke et al., Cell Rep Medicine, 2021 | VYDPLQPELDSF | VYDPLQPELDSF | VYDPLQPELDSF | VYDPLQPELDSF | VYDPLQPELDSF | 100,0 | 99,5 |
| | S1137-1145 | VYDPLQPEL | A*2402 | Hu et al., Genes & Diseases, 2021 | VYDPLQPEL | VYDPLQPEL | VYDPLQPEL | VYDPLQPEL | VYDPLQPEL | 100,0 | 100,0 |
| | S1147-1155 | SFKEELDKY | A*2902 | Tarke et al., Cell Rep Medicine, 2021 | SFKEELDKY | SFKEELDKY | SFKEELDKY | SFKEELDKY | SFKEELDKY | 100,0 | 98,8 |
| | S1173-1181 | NASVVNIQK | A*6801 | Tarke et al., Cell Rep Medicine, 2021 | NASVVNIQK | NASVVNIQK | NASFVNIQK | NASVVNIQK | NASVVNIQK | 100,0 | 97,6 |
| | S1181-1179 | KEIDRLNEV | B*4402 / B*4403<br>B*4001 | Tarke et al., Cell Rep Medicine, 2021<br>Tarke et al., Cell Rep Medicine, 2021 | KEIDRLNEV | KEIDRLNEV | KEIDRLNEV | KEIDRLNEV | KEIDRLNEV | 100,0 | 97,0 |
| | S1185-1193 | RLNEVAKNL | A*0201 | Shomuradova et al., Immunity, 2020 | RLNEVAKNL | RLNEVAKNL | RLNEVAKNL | RLNEVAKNL | RLNEVAKNL | 100,0 | 97,0 |
| | S1192-1200 | NLNESLIDL | A*0201 | Shomuradova et al., Immunity, 2020 | NLNESLIDL | NLNESLIDL | NLNESLIDL | NLNESLIDL | NLNESLIDL | 100,0 | 98,8 |
| | S1194-1203 | NESLIDLQEL | B*4001 | Tarke et al., Cell Rep Medicine, 2021 | NESLIDLQEL | NESLIDLQEL | NESLIDLQEL | NESLIDLQEL | NESLIDLQEL | 100,0 | 99,2 |
| | S1201-1209 | QELGKYEQY | B*4402 / B*4403 | Tarke et al., Cell Rep Medicine, 2021 | QELGKYEQY | QELGKYEQY | QELGKYEQY | QELGKYEQY | QELGKYEQY | 100,0 | 99,7 |
| vN2 | N289-298 | QELIRQGTDY | B*4402 | Tarke et al., Cell Rep Medicine, 2021 | QELIRQGTDY | QELIRQGTDY | QELIRQGTDY | QELIRQGTDY | QDLIRQGTDY | 90,0 | 93,8 |
| | N290-298 | ELIRQGTDY | A*2601 | Tarke et al., Cell Rep Medicine, 2021 | ELIRQGTDY | ELIRQGTDY | ELIRQGTDY | ELIRQGTDY | DLIRQGTDY | 88,9 | 93,1 |
| | N297-307 | DYKHWPQIAQF | A*24 | Nelde et al., 2020 | DYKHWPQIAQF | DYKHWPQIAQF | DYKHWPQIAQF | DYKHWPQIAQF | DYKHWPQIAQF | 100,0 | 98,5 |
| | N305-314 | AQFAPSASAF | B*1501 | Schulien et al., Nat Med, 2020 | AQFAPSASAF | AQFAPSASAF | AQFAPSASAF | AQFAPSASAF | AQFAPSASAF | 100,0 | 100,0 |
| | N306-315 | QFAPSASAFF | A*2402 | Tarke et al., Cell Rep Medicine, 2021 | QFAPSASAFF | QFAPSASAFF | QFAPSASAFF | QFAPSASAFF | QFAPSASAFF | 100,0 | 100,0 |
| | N307-315 | FAPSASAFF | B*3501 | Tarke et al., Cell Rep Medicine, 2021 | FAPSASAFF | FAPSASAFF | FAPSASAFF | FAPSASAFF | FAPSASAFF | 100,0 | 100,0 |
| | N308-317 | APSASAFFGM | B*0702 | Tarke et al., Cell Rep Medicine, 2021 | APSASAFFGM | APSASAFFGM | APSASAFFGM | APSASAFFGM | APSASAFFGM | 100,0 | 100,0 |
| | N310-319 | SASAFFGMSR | A*6801 | Tarke et al., Cell Rep Medicine, 2021 | SASAFFGMSR | SASAFFGMSR | SASAFFGMSR | SASAFFGMSR | SASAFFGMSR | 100,0 | 100,0 |
| | N311-319 | ASAFFGMSR | A*1101<br>A*6801 | Ferretti et al., Immunity, 2020<br>Tarke et al., Cell Rep Medicine, 2021 | ASAFFGMSR | ASAFFGMSR | ASAFFGMSR | ASAFFGMSR | ASAFFGMSR | 100,0 | 100,0 |
| | N316-324 | GMSRIGMEV | A*0201 | Schulien et al., Nat Med, 2020<br>Hu et al., Genes & Diseases, 2021<br>Tarke et al., Cell Rep Medicine, 2021 | GMSRIGMEV | GMSRIGMEV | GMSRIGMEV | GMSRIGMEV | GMSRIGMEV | 100,0 | 99,7 |
| | N322-331 | MEVTPSGTWL | B*4001 | Schulien et al., Nat Med, 2020<br>Peng et al., Nat Immunol, 2020<br>Nelde et al., 2020<br>Tarke et al., Cell Rep Medicine, 2021 | MEVTPSGTWL | MEVTPSGTWL | MEVTPSGTWL | MEVTPSGTWL | MEVTPSGTWL | 100,0 | 98,9 |
| | N322-330 | MEVTPSGTW | B*4402 / B*4403 | Schulien et al., Nat Med, 2020 | MEVTPSGTW | MEVTPSGTW | MEVTPSGTW | MEVTPSGTW | MEVTPSGTW | 100,0 | 98,8 |
| | N323-333 | EVTPSGTWLTY | A*2601 | Tarke et al., Cell Rep Medicine, 2021 | EVTPSGTWLTY | EVTPSGTWLTY | EVTPSGTWLTY | EVTPSGTWLTY | EVTPSGTWLTY | 100,0 | 98,8 |
| | N324-333 | VTPSGTWLTY | A*3002 | Tarke et al., Cell Rep Medicine, 2021 | VTPSGTWLTY | VTPSGTWLTY | VTPSGTWLTY | VTPSGTWLTY | VTPSGTWLTY | 100,0 | 98,6 |
| | N325-333 | TPSGTWLTY | B*3501 | Tarke et al., Cell Rep Medicine, 2021 | TPSGTWLTY | TPSGTWLTY | TPSGTWLTY | TPSGTWLTY | TPSGTWLTY | 100,0 | 98,5 |
| | N338-346 | KLDDKDPNF | A*0201 | Poran et al., Genome Medicine, 2020<br>Hu et al., Genes & Diseases, 2021 | KLDDKDPNF | KLDDKDPNF | KLDDKDPNF | KLDDKDPNF | KLDDKDPQF | 88,9 | 91,3 |
| | N360-369 | YKTFPPTEPK | A*6801 | Tarke et al., Cell Rep Medicine, 2021 | YKTFPPTEPK | YKTFPPTEPK | YKTFPPTEPK | YKTFPPTEPK | YKTFPPTEPK | 100,0 | 99,2 |
| | N361-374 | KTFPPTEPKKDKKK | A*0301 | Tarke et al., Cell Rep Medicine, 2021 | KTFPPTEPKKDKKK | KTFPPTEPKKDKKK | KTFPPTEPKKDKKK | KTFPPTEPKKDKKK | KTFPPTEPKKDKKK | 100,0 | 99,4 |
| | N361-370 | KTFPPTEPKK | A*03<br>A*0301 / A*1101 | Nelde et al., 2020<br>Tarke et al., Cell Rep Medicine, 2021 | KTFPPTEPKK | KTFPPTEPKK | KTFPPTEPKK | KTFPPTEPKK | KTFPPTEPKK | 100,0 | 99,2 |
| | N361-369 | KTFPPTEPK | A*0301 / A*1101<br>A*0301 / A*1101<br>A*0301 / A*1101<br>A*0301 / A*1101<br>A*03 / A*11 | Ferretti et al., Immunity, 2020<br>Peng et al., Nat Immunol, 2020<br>Hu et al., Genes & Diseases, 2021<br>Gao et al., IScience, 2021<br>Kared et al., JCI, 2021 | KTFPPTEPK | KTFPPTEPK | KTFPPTEPK | KTFPPTEPK | KTFPPTEPK | 100,0 | 99,1 |
| | N395-403 | LPAADLDDF | A*6801<br>B*3501 | Tarke et al., Cell Rep Medicine, 2021<br>Tarke et al., Cell Rep Medicine, 2021 | LPAADLDDF | LPAADLDDF | LPAADLDDF | LPAADLDDF | LPAADMDDF | 88,9 | 91,9 |

Mutations and deletions from original Wuhan SARS-CoV-2 sequence are highlighted in red
Mutated CD8 T-cell epitopes no more able to bind to MHC-restriction molecule for SARS-CoV-2 VOC or SARS-CoV-1 are colored in grey

### Table S3

| Vaccine Antigen | Position | Sequence | Source | B.1.1.7 (Alpha Variant) | B.1.351 (Beta Variant) | P.1 (Gamma Variant) | B.1.617.2 (Delta Variant) | SARS-CoV-1 | Sequence homology (%) with SARS-CoV-1 | Sequence homology (mean %) with 38 sarbecoviruses |
|---|---|---|---|---|---|---|---|---|---|---|
| vS1 | S126-140 | VVIKVCEFQFCNDPF | Mateus et al., Science, 2020<br>Tarke et al., Cell Rep Medicine, 2021 | VVIKVCEFQFCNDPF | VVIKVCEFQFCNDPF | VVIKVCEFQFCNYPF | VVIKVCEFQFCNDPF | VVIRACNFELCDNPF | 53,3 | 57,3 |
| | S131-145 | CEFQFCNDPFLGVYY | Mateus et al., Science, 2020<br>Tarke et al., Cell Rep Medicine, 2021 | CEFQFCNDPFLGVY--- | CEFQFCNDPFLGVY--Y | CEFQFCNYPFLGVY--Y | CEFQFCNDPFLDVY--Y | CNFELCDNPFFAVSKPM | 40,0 | 49,2 |
| | S141-155 | LGVYYHKNNKSWMES | Mateus et al., Science, 2020<br>Tarke et al., Cell Rep Medicine, 2021 | LGVY---HKNNKSWMES | LGVY--YHKNNKSWMES | LGVY--YHKNNKSWMES | LDVY--YHKNNKSWMES | FAVSKPMGTQTHTM--- | 6,7 | 31,5 |
|  | S146-160 | HKNNKSWMESEFRVY | Tarke et al., Cell Rep Medicine, 2021 | HKNNKSWMESEFRVY | HKNNKSWMESEFRVY | HKNNKSWMESEFRVY | HKNNKSWMESG--VY | GTQTHTM------IF | 0,0 | 33,7 |
| | S161-175 | SSANNCTFEYVSQPF | Tarke et al., Cell Rep Medicine, 2021 | SSANNCTFEYVSQPF | SSANNCTFEYVSQPF | SSANNCTFEYVSQPF | SSANNCTFEYVSQPF | DNAFNCTFEYISDAF | 60,0 | 60,5 |
| | S166-180 | CTFEYVSQPFLMDLE | Peng et al., Nat Immunol, 2020<br>Mateus et al., Science, 2020<br>Tarke et al., Cell Rep Medicine, 2021 | CTFEYVSQPFLMDLE | CTFEYVSQPFLMDLE | CTFEYVSQPFLMDLE | CTFEYVSQPFLMDLE | CTFEYISDAFSLDVS | 53,3 | 53,7 |
| | S171-185 | VSQPFLMDLEGKQGN | Tarke et al., Cell Rep Medicine, 2021 | VSQPFLMDLEGKQGN | VSQPFLMDLEGKQGN | VSQPFLMDLEGKQGN | VSQPFLMDLEGKQGN | ISDAFSLDVSEKSGN | 40,0 | 49,5 |
| | S176-190 | LMDLEGKQGNFKNLR | Tarke et al., Cell Rep Medicine, 2021 | LMDLEGKQGNFKNLR | LMDLEGKQGNFKNLR | LMDLEGKQGNFKNLS | LMDLEGKQGNFKNLR | SLDVSEKSGNFKHLR | 53,3 | 58,0 |
| | S181-195 | GKQGNFKNLREFVFK | Tarke et al., Cell Rep Medicine, 2021 | GKQGNFKNREFVFK | GKQGNFKNREFVFK | GKQGNFKNSEFVFK | GKQGNFKNREFVFK | EKSGNFKHLREFVFK | 80,0 | 74,8 |
| | S191-205 | EFVFKNIDGYFKIYS | Tarke et al., Cell Rep Medicine, 2021 | EFVFKNIDGYFKIYS | EFVFKNIDGYFKIYS | EFVFKNIDGYFKIYS | EFVFKNIDGYFKIYS | EFVFKNKDGFLYVYK | 60,0 | 67,7 |
| | S196-210 | NIDGYFKIYSKHTPI | Mateus et al., Science, 2020<br>Tarke et al., Cell Rep Medicine, 2021 | NIDGYFKIYSKHTPI | NIDGYFKIYSKHTPI | NIDGYFKIYSKHTPI | NIDGYFKIYSKHTPI | NKDGFLYVYKGYQPI | 40,0 | 54,4 |
| | S201-215 | FKIYSKHTPINLVRD | Tarke et al., Cell Rep Medicine, 2021 | FKIYSKHTPINLVRD | FKIYSKHTPINLVRG | FKIYSKHTPINLVRD | FKIYSKHTPINLVRD | LYVYKGYQPIDVVRD | 40,0 | 47,0 |
| | S206-220 | KHTPINLVRDLPQGF | Tarke et al., Cell Rep Medicine, 2021 | KHTPINLVRDLPQGF | KHTPINLVRGLPQGF | KHTPINLVRDLPQGF | KHTPINLVRDLPQGF | GYQPIDVVRDLPSGF | 60,0 | 55,3 |
| | S211-225 | NLVRDLPQGFSALEP | Tarke et al., Cell Rep Medicine, 2021 | NLVRDLPQGFSALEP | NLVRGLPQGFSALEP | NLVRDLPQGFSALEP | NLVRDLPQGFSALEP | DVVRDLPSGFNTLKP | 60,0 | 60,7 |
| | S216-230 | LPQGFSALEPLVDLP | Mateus et al., Science, 2020<br>Tarke et al., Cell Rep Medicine, 2021 | LPQGFSALEPLVDLP | LPQGFSALEPLVDLP | LPQGFSALEPLVDLP | LPQGFSALEPLVDLP | LPSGFNTLKPIFKLP | 53,3 | 64,3 |
| | S221-235 | SALEPLVDLPIGINI | Tarke et al., Cell Rep Medicine, 2021 | SALEPLVDLPIGINI | SALEPLVDLPIGINI | SALEPLVDLPIGINI | SALEPLVDLPIGINI | NTLKPIFKLPLGINI | 53,3 | 66,5 |
| | S231-245 | IGINITRFQTLLALH | Mateus et al., Science, 2020<br>Tarke et al., Cell Rep Medicine, 2021 | IGINITRFQTLLALH | IGINITRFQTL---H | IGINITRFQTLLALH | IGINITRFQTLLALH | LGINITNFRAILT-- | 46,7 | 56,0 |
| | S235-249 | ITRFQTLLALHRSYL | Nelde et al., 2020 | ITRFQTLLALHRSYL | ITRFQTL---HRSYL | ITRFQTLLALHRSYL | ITRFQTLLALHRSYL | ITNFRAILT---AFS | 26,7 | 41,1 |
| | S236-250 | TRFQTLLALHRSYLT | Mateus et al., Science, 2020<br>Tarke et al., Cell Rep Medicine, 2021 | TRFQTLLALHRSYLT | TRFQTL---HRSYLT | TRFQTLLALHRSYLT | TRFQTLLALHRSYLT | TNFRAILT---AFSP | 20,0 | 35,7 |
| vRBD | S321-335 | QPTESIVRFPNITNL | Mateus et al., Science, 2020 | QPTESIVRFPNITNL | QPTESIVRFPNITNL | QPTESIVRFPNITNL | QPTESIVRFPNITNL | VPSGDVVRFPNITNL | 66,7 | 72,1 |
| | S326-340 | IVRFPNITNLCPFGE | Mateus et al., Science, 2020<br>Tarke et al., Cell Rep Medicine, 2021 | IVRFPNITNLCPFGE | IVRFPNITNLCPFGE | IVRFPNITNLCPFGE | IVRFPNITNLCPFGE | VVRFPNITNLCPFGE | 93,3 | 84,3 |
| | S336-350 | CPFGEVFNATRFASV | Mateus et al., Science, 2020<br>Tarke et al., Cell Rep Medicine, 2021 | CPFGEVFNATRFASV | CPFGEVFNATRFASV | CPFGEVFNATRFASV | CPFGEVFNATRFASV | CPFGEVFNATKFPSV | 86,7 | 81,6 |
| | S341-355 | VFNATRFASVYAWNR | Mateus et al., Science, 2020 | VFNATRFASVYAWNR | VFNATRFASVYAWNR | VFNATRFASVYAWNR | VFNATRFASVYAWNR | VFNATKFPSVYAWER | 80,0 | 83,1 |
| | S346-360 | RFASVYAWNRKRISN | Tarke et al., Cell Rep Medicine, 2021 | RFASVYAWNRKRISN | RFASVYAWNRKRISN | RFASVYAWNRKRISN | RFASVYAWNRKRISN | KFPSVYAWERKKISN | 73,3 | 75,1 |
| | S351-365 | YAWNRKRISNCVADY | Peng et al., Nat Immunol, 2020<br>Tarke et al., Cell Rep Medicine, 2021 | YAWNRKRISNCVADY | YAWNRKRISNCVADY | YAWNRKRISNCVADY | YAWNRKRISNCVADY | YAWERKKISNCVADY | 86,7 | 84,1 |
| | S356-370 | KRISNCVADYSVLYN | Tarke et al., Cell Rep Medicine, 2021 | KRISNCVADYSVLYN | KRISNCVADYSVLYN | KRISNCVADYSVLYN | KRISNCVADYSVLYN | KKISNCVADYSVLYN | 93,3 | 85,0 |
| | S361-375 | CVADYSVLYNSASFS | Tarke et al., Cell Rep Medicine, 2021 | CVADYSVLYNSASFS | CVADYSVLYNSASFS | CVADYSVLYNSASFS | CVADYSVLYNSASFS | CVADYSVLYNSTFFS | 86,7 | 88,8 |
| | S366-380 | SVLYNSASFSTFKCY | Tarke et al., Cell Rep Medicine, 2021 | SVLYNSASFSTFKCY | SVLYNSASFSTFKCY | SVLYNSASFSTFKCY | SVLYNSASFSTFKCY | SVLYNSTFFSTFKCY | 86,7 | 88,5 |
| | S371-385 | SASFSTFKCYGVSPT | Tarke et al., Cell Rep Medicine, 2021 | SASFSTFKCYGVSPT | SASFSTFKCYGVSPT | SASFSTFKCYGVSPT | SASFSTFKCYGVSPT | STFFSTFKCYGVSAT | 80,0 | 87,6 |
| | S376-390 | TFKCYGVSPTKLNDL | Tarke et al., Cell Rep Medicine, 2021 | TFKCYGVSPTKLNDL | TFKCYGVSPTKLNDL | TFKCYGVSPTKLNDL | TFKCYGVSPTKLNDL | TFKCYGVSATKLNDL | 93,3 | 90,8 |
| | S381-395 | GVSPTKLNDLCFTNV | Peng et al., Nat Immunol, 2020 | GVSPTKLNDLCFTNV | GVSPTKLNDLCFTNV | GVSPTKLNDLCFTNV | GVSPTKLNDLCFTNV | GVSATKLNDLCFSNV | 86,7 | 86,1 |
| | S386-400 | KLNDLCFTNVYADSF | Tarke et al., Cell Rep Medicine, 2021 | KLNDLCFTNVYADSF | KLNDLCFTNVYADSF | KLNDLCFTNVYADSF | KLNDLCFTNVYADSF | KLNDLCFSNVYADSF | 93,3 | 87,9 |
| | S396-410 | YADSFVIRGDEVRQI | Tarke et al., Cell Rep Medicine, 2021 | YADSFVIRGDEVRQI | YADSFVIRGDEVRQI | YADSFVIRGDEVRQI | YADSFVIRGDEVRQI | YADSFVVKGDDVRQI | 80,0 | 77,1 |
| | S401-415 | VIRGDEVRQIAPGQT | Tarke et al., Cell Rep Medicine, 2021 | VIRGDEVRQIAPGQT | VIRGDEVRQIAPGQT | VIRGDEVRQIAPGQT | VIRGDEVRQIAPGQT | VVKGDDVRQIAPGQT | 80,0 | 78,0 |
| | S416-430 | GKIADYNYKLPDDFT | Mateus et al., Science, 2020 | GKIADYNYKLPDDFT | GNIADYNYKLPDDFT | GTIADYNYKLPDDFT | GKIADYNYKLPDDFT | GVIADYNYKLPDDFM | 86,7 | 92,4 |
| | S421-435 | YNYKLPDDFTGCVIA | Mateus et al., Science, 2020 | YNYKLPDDFTGCVIA | YNYKLPDDFTGCVIA | YNYKLPDDFTGCVIA | YNYKLPDDFTGCVIA | YNYKLPDDFMGCVLA | 86,7 | 96,2 |
| | S431-445 | GCVIAWNSNNLDSKV | Mateus et al., Science, 2020 | GCVIAWNSNNLDSKV | GCVIAWNSNNLDSKV | GCVIAWNSNNLDSKV | GCVIAWNSNNLDSKV | GCVLAWNTRNIDATS | 53,3 | 60,9 |
| | S436-450 | WNSNNLDSKVGGNYN | Tarke et al., Cell Rep Medicine, 2021 | WNSNNLDSKVGGNYN | WNSNNLDSKVGGNYN | WNSNNLDSKVGGNYN | WNSNNLDSKVGGNYN | WNTRNIDATSTGNYN | 53,3 | 43,6 |
| | S446-460 | GGNYNYLYRLFRKSN | Mateus et al., Science, 2020 | GGNYNYLYRLFRKSN | GGNYNYLYRLFRKSN | GGNYNYLYRLFRKSN | GGNYNYRYRLFRKSN | TGNYNYKYRYLRHGK | 53,3 | 51,0 |
| | S461-475 | LKPFERDISTEIYQA | Tarke et al., Cell Rep Medicine, 2021 | LKPFERDISTEIYQA | LKPFERDISTEIYQA | LKPFERDISTEIYQA | LKPFERDISTEIYQA | LRPFERDISNVPFSP | 53,3 | 62,0 |
|  | S471-485 | EIYQAGSTPCNGVEG | Tarke et al., Cell Rep Medicine, 2021 | EIYQAGSTPC-----NGVEG | EIYQAGSTPC-----NGVKG | EIYQAGSTPC-----NGVKG | EIYQAGSKPC-----NGVEG | VPFSPDGKPCTPPALNC--- | 20,0 | 31,5 |
| | S486-500 | FNCYFPLQSYGFQPT | Mateus et al., Science, 2020<br>Tarke et al., Cell Rep Medicine, 2021 | FNCYFPLQSYGFQPT | FNCYFPLQSYGFQPT | FNCYFPLQSYGFQPT | FNCYFPLQSYGFQPT | ---YWPLNDYGFYTT | 46,7 | 44,3 |
| | S506-520 | QPYRVVVLSFELLHA | Tarke et al., Cell Rep Medicine, 2021 | QPYRVVVLSFELLHA | QPYRVVVLSFELLHA | QPYRVVVLSFELLHA | QPYRVVVLSFELLHA | QPYRVVVLSFELLNA | 93,3 | 87,2 |
| | S511-525 | VVLSFELLHAPATVC | Peng et al., Nat Immunol, 2020<br>Tarke et al., Cell Rep Medicine, 2021 | VVLSFELLHAPATVC | VVLSFELLHAPATVC | VVLSFELLHAPATVC | VVLSFELLHAPATVC | VVLSFELLNAPATVC | 93,3 | 93,7 |
| vS2 | S1056-1070 | APHGVVFLHVTYVPA | Mateus et al., Science, 2020<br>Tarke et al., Cell Rep Medicine, 2021 | APHGVVFLHVTYVPA | APHGVVFLHVTYVPA | APHGVVFLHVTYVPA | APHGVVFLHVTYVPA | APHGVVFLHVTYVPS | 93,3 | 94,1 |
| | S1076-1090 | TTAPAICHDGKAHFP | Mateus et al., Science, 2020 | TTAPAICHDGKAHFP | TTAPAICHDGKAHFP | TTAPAICHDGKAHFP | TTAPAICHDGKAHFP | TTAPAICHEGKAYFP | 86,7 | 89,5 |
| | S1086-1100 | KAHFPREGVFVSNGT | Mateus et al., Science, 2020 | KAHFPREGVFVSNGT | KAHFPREGVFVSNGT | KAHFPREGVFVSNGT | KAHFPREGVFVSNGT | KAYFPREGVFVFNGT | 86,7 | 93,0 |
| | S1091-1105 | REGVFVSNGTHWFVT | Mateus et al., Science, 2020 | REGVFVSNGTHWFVT | REGVFVSNGTHWFVT | REGVFVSNGTHWFVT | REGVFVSNGTHWFVT | REGVFVFNGTSWFIT | 80,0 | 87,9 |
| | S1101-1115 | HWFVTQRNFYEPQII | Mateus et al., Science, 2020<br>Tarke et al., Cell Rep Medicine, 2021 | HWFVTQRNFYEPQII | HWFVTQRNFYEPQII | HWFVTQRNFYEPQII | HWFVTQRNFYEPQII | SWFITQRNFFSPQII | 73,3 | 82,5 |
| | S1106-1120 | QRNFYEPQIITTDNT | Tarke et al., Cell Rep Medicine, 2021 | QRNFYEPQIITTHNT | QRNFYEPQIITTDNT | QRNFYEPQIITTDNT | QRNFYEPQIITTDNT | QRNFFSPQIITTDNT | 86,7 | 91,9 |
| | S1121-1135 | FVSGNCDVVIGIVNN | Tarke et al., Cell Rep Medicine, 2021 | FVSGNCDVVIGIVNN | FVSGNCDVVIGIVNN | FVSGNCDVVIGIVNN | FVSGNCDVVIGIVNN | FVSGNCDVVIGIINN | 93,3 | 89,7 |
| | S1131-1145 | GIVNNTVYDPLQPEL | Tarke et al., Cell Rep Medicine, 2021 | GIVNNTVYDPLQPEL | GIVNNTVYDPLQPEL | GIVNNTVYDPLQPEL | GIVNNTVYDPLQPEL | GIINNTVYDPLQPEL | 93,3 | 95,1 |
| | S1141-1155 | LQPELDSFKEELDKY | Tarke et al., Cell Rep Medicine, 2021 | LQPELDSFKEELDKY | LQPELDSFKEELDKY | LQPELDSFKEELDKY | LQPELDSFKEELDKY | LQPELDSFKEELDKY | 100,0 | 98,9 |
| | S1151-1165 | ELDKYFKNHTSPDVD | Mateus et al., Science, 2020<br>Tarke et al., Cell Rep Medicine, 2021 | ELDKYFKNHTSPDVD | ELDKYFKNHTSPDVD | ELDKYFKNHTSPDVD | ELDKYFKNHTSPDVD | ELDKYFKNHTSPDVD | 100,0 | 98,9 |
| | S1166-1180 | LGDISGINASVVNIQ | Mateus et al., Science, 2020 | LGDISGINASVVNIQ | LGDISGINASVVNIQ | LGDISGINASFVNIQ | LGDISGINASVVNIQ | LGDISGINASVVNIQ | 100,0 | 97,3 |
| | S1171-1185 | GINASVVNIQKEIDR | Mateus et al., Science, 2020<br>Tarke et al., Cell Rep Medicine, 2021 | GINASVVNIQKEIDR | GINASVVNIQKEIDR | GINASFVNIQKEIDR | GINASVVNIQKEIDR | GINASVVNIQKEIDR | 100,0 | 97,1 |
| | S1186-1200 | LNEVAKNLNESLIDL | Mateus et al., Science, 2020 | LNEVAKNLNESLIDL | LNEVAKNLNESLIDL | LNEVAKNLNESLIDL | LNEVAKNLNESLIDL | LNEVAKNLNESLIDL | 100,0 | 98,2 |
| vN2 | N276-290 | RRGPEQTQGNFGDQE | Tarke et al., Cell Rep Medicine, 2021 | RRGPEQTQGNFGDQE | RRGPEQTQGNFGDQE | RRGPEQTQGNFGDQE | RRGPEQTQGNFGDQE | RRGPEQTQGNFGDQD | 93,3 | 96,6 |
| | N286-300 | FGDQELIRQGTDYKH | Tarke et al., Cell Rep Medicine, 2021 | FGDQELIRQGTDYKH | FGDQELIRQGTDYKH | FGDQELIRQGTDYKH | FGDQELIRQGTDYKH | FGDQDLIRQGTDYKH | 93,3 | 95,3 |
| | N291-305 | LIRQGTDYKHWPQIA | Le Bert et al., Nature, 2020<br>Tarke et al., Cell Rep Medicine, 2021 | LIRQGTDYKHWPQIA | LIRQGTDYKHWPQIA | LIRQGTDYKHWPQIA | LIRQGTDYKHWPQIA | LIRQGTDYKHWPQIA | 100,0 | 98,0 |
| | N296-310 | TDYKHWPQIAQFAPS | Tarke et al., Cell Rep Medicine, 2021 | TDYKHWPQIAQFAPS | TDYKHWPQIAQFAPS | TDYKHWPQIAQFAPS | TDYKHWPQIAQFAPS | TDYKHWPQIAQFAPS | 100,0 | 98,7 |
| | N298-312 | YKHWPQIAQFAPSAS | Nelde et al., 2020 | YKHWPQIAQFAPSAS | YKHWPQIAQFAPSAS | YKHWPQIAQFAPSAS | YKHWPQIAQFAPSAS | YKHWPQIAQFAPSAS | 100,0 | 99,5 |
| | N301-315 | WPQIAQFAPSASAFF | Le Bert et al., Nature, 2020<br>Tarke et al., Cell Rep Medicine, 2021 | WPQIAQFAPSASAFF | WPQIAQFAPSASAFF | WPQIAQFAPSASAFF | WPQIAQFAPSASAFF | WPQIAQFAPSASAFF | 100,0 | 100,0 |
| | N306-320 | QFAPSASAFFGMSRI | Tarke et al., Cell Rep Medicine, 2021 | QFAPSASAFFGMSRI | QFAPSASAFFGMSRI | QFAPSASAFFGMSRI | QFAPSASAFFGMSRI | QFAPSASAFFGMSRI | 100,0 | 100,0 |
| | N311-325 | ASAFFGMSRIGMEVT | Nelde et al., 2020<br>Tarke et al., Cell Rep Medicine, 2021 | ASAFFGMSRIGMEVT | ASAFFGMSRIGMEVT | ASAFFGMSRIGMEVT | ASAFFGMSRIGMEVT | ASAFFGMSRIGMEVT | 100,0 | 99,8 |
| | N321-340 | GMEVTPSGTWLTYTGAIKLD | Le Bert et al., Nature, 2020<br>Tarke et al., Cell Rep Medicine, 2021 | GMEVTPSGTWLTYTGAIKLD | GMEVTPSGTWLTYTGAIKLD | GMEVTPSGTWLTYTGAIKLD | GMEVTPSGTWLTYTGAIKLD | GMEVTPSGTWLTYHGAIKLD | 95,0 | 95,3 |
| | N326-340 | PSGTWLTYTGAIKLD | Mateus et al., Science, 2020<br>Tarke et al., Cell Rep Medicine, 2021 | PSGTWLTYTGAIKLD | PSGTWLTYTGAIKLD | PSGTWLTYTGAIKLD | PSGTWLTYTGAIKLD | PSGTWLTYHGAIKLD | 93,3 | 93,9 |
| | N328-342 | GTWLTYTGAIKLDDK | Nelde et al., 2020 | GTWLTYTGAIKLDDK | GTWLTYTGAIKLDDK | GTWLTYTGAIKLDDK | GTWLTYTGAIKLDDK | GTWLTYHGAIKLDDK | 93,3 | 94,6 |
| | N329-346 | TWLTYTGAIKLDDKDPNF | Peng et al., Nat Immunol, 2020 | TWLTYTGAIKLDDKDPNF | TWLTYTGAIKLDDKDPNF | TWLTYTGAIKLDDKDPNF | TWLTYTGAIKLDDKDPNF | TWLTYHGAIKLDDKDPQF | 88,9 | 91,1 |
| | N331-345 | LTYTGAIKLDDKDPN | Tarke et al., Cell Rep Medicine, 2021 | LTYTGAIKLDDKDPN | LTYTGAIKLDDKDPN | LTYTGAIKLDDKDPN | LTYTGAIKLDDKDPN | LTYHGAIKLDDKDPQ | 86,7 | 89,4 |
| | N336-350 | AIKLDDKDPNFKDQV | Tarke et al., Cell Rep Medicine, 2021 | AIKLDDKDPNFKDQV | AIKLDDKDPNFKDQV | AIKLDDKDPNFKDQV | AIKLDDKDPNFKDQV | AIKLDDKDPQFKDNV | 86,7 | 89,5 |
| | N344-361 | PNFKDQVILLNKHIDAYK | Peng et al., Nat Immunol, 2020 | PNFKDQVILLNKHIDAYK | PNFKDQVILLNKHIDAYK | PNFKDQVILLNKHIDAYK | PNFKDQVILLNKHIDAYK | PQFKDNVILLNKHIDAYK | 88,9 | 90,8 |
| | N346-360 | FKDQVILLNKHIDAY | Tarke et al., Cell Rep Medicine, 2021 | FKDQVILLNKHIDAY | FKDQVILLNKHIDAY | FKDQVILLNKHIDAY | FKDQVILLNKHIDAY | FKDNVILLNKHIDAY | 93,3 | 94,2 |
| | N351-365 | ILLNKHIDAYKTFPP | Tarke et al., Cell Rep Medicine, 2021 | ILLNKHIDAYKTFPP | ILLNKHIDAYKTFPP | ILLNKHIDAYKTFPP | ILLNKHIDAYKTFPP | ILLNKHIDAYKTFPP | 100,0 | 99,1 |
| | N356-370 | HIDAYKTFPPTEPKK | Tarke et al., Cell Rep Medicine, 2021 | HIDAYKTFPPTEPKK | HIDAYKTFPPTEPKK | HIDAYKTFPPTEPKK | HIDAYKTFPPTEPKK | HIDAYKTFPPTEPKK | 100,0 | 99,5 |
| | N386-400 | QKKQQTVTLLPAADL | Tarke et al., Cell Rep Medicine, 2021 | QKKQQTVTLLPAADL | QKKQQTVTLLPAADL | QKKQQTVTLLPAADL | QKKQQTVTLLPAADL | QKKQPTVTLLPAADM | 86,7 | 89,7 |
| | N391-405 | TVTLLPAADLDDFSK | Tarke et al., Cell Rep Medicine, 2021 | TVTLLPAADLDDFSK | TVTLLPAADLDDFSK | TVTLLPAADLDDFSK | TVTLLPAADLDDFSK | TVTLLPAADMDDFSR | 86,7 | 90,6 |
| | N397-411 | AADLDDFSKQLQQSM | Nelde et al., 2020 | AADLDDFSKQLQQSM | AADLDDFSKQLQQSM | AADLDDFSKQLQQSM | AADLDDFSKQLQQSM | AADMDDFSRQLQNSM | 80,0 | 85,9 |

Mutations and deletions from original Wuhan SARS-CoV-2 sequence are highlighted in red
